# Systemic degradation of repressive transcription factors gates gene expression and cell fate specification

**DOI:** 10.64898/2026.06.16.732780

**Authors:** Predrag Jevtić, Samuel R. Witus, Devlon M. McCloud, Zhi Yang, Ana Milunović Jevtić, Heegwang Roh, Michael Rapé

## Abstract

While proteasomes are best known for eliminating defective proteins or turning off signaling pathways, they also enable crucial cellular activities. Critical among these, proteasomes allow cells to initiate gene expression, but underlying targets and regulatory mechanisms remain poorly understood. Here, we report that proteasomes drive the systemic degradation of repressive transcription factors to eject TLE/Groucho-family co-repressors from chromatin and thereby constantly free transcription start sites for activator binding. This circuitry requires the E3 ligase SCF^FBXL14^, which modifies its targets dependent on presentation by TLEs, but independently of their identity. The continuous cycling of co-repressors off chromatin, as achieved by systemic turnover of a protein family, is essential for stem cells to translate developmental cues into lineage-specific gene expression, and it is disrupted by cancer mutations in *TLE1* that impair SCF^FBXL14^-recruitment. We conclude that systemic degradation of repressive transcription factors establishes co-repressor dynamics required for genes expression and cell fate specification.

## Introduction

Ubiquitin- and proteasome-dependent degradation provides cells with a powerful mechanism to clear aberrant proteins or turn off signaling pathways ^1,2^. These proteolytic events are typically regulated and depend on conformational changes or posttranslational modifications that mark specific proteins for recognition by their cognate E3 ubiquitin ligases ^3^. However, cells also rely on proteasomes to turn on critical functions ^3–6^, but the targets and molecular mechanisms underlying such regulation are often poorly understood.

Illustrating this complexity, proteasomes play opposing roles in controlling the gene expression programs that orchestrate cell division, differentiation, and survival. In a well-understood inhibitory capacity, proteasomes target transcription factors, co-activators, or RNA polymerase subunits to turn off transcriptional events ^7–12^. However, proteasomes are also broadly required for cells to elicit new gene expression ^13^. Proteasomes can exert this function by eliminating select inhibitory proteins, such as IκBα, which releases transcription factors for chromatin binding ^14^. Degradation of specific transcription factors, such as c-MYC, can also facilitate the progression from transcription initiation to elongation or enable cells to re-direct co-activators and RNA polymerase to new targets ^15–20^. While these observations revealed dual roles of proteasomes in transcriptional control, protein degradation is broadly required for mammalian gene expression and key regulatory circuits likely remain to be discovered.

Most studies dissecting the proteolytic control of gene expression focused on activating factors that help RNA polymerase bind specific transcription start sites. Equally important, however, are repressive transcription factors that prevent inappropriate or untimely gene expression. Such proteins include members of the HES, FOX, TLX and NKX2 families that are essential for accurate cell fate specification and mutated in developmental diseases and cancer ^21–26^. Repressive transcription factors act by recruiting complexes, referred to as co-repressors ^27–29^, that in turn attract chromatin modifiers to restrict the access of activators and RNA polymerase to promoters or enhancers ^29,30^. Most co-repressors are expressed at high levels and form oligomeric assemblies that bind DNA by recognizing transcription factors and nucleosomes at the same time ^30–36^. While the abundance and multivalent chromatin association of co-repressors enables cells to stably shut off transcription, initiation of new gene expression programs requires cells to overcome such regulation through mechanisms that are still ill-defined.

The release of co-repressors from chromatin is particularly important during development, when activation of lineage-specific gene expression drives the formation of the many cell types that comprise metazoan organisms. Accordingly, despite their stable association with DNA *in vitro*, co-repressors and their effectors are highly responsive to differentiation signals ^37–40^. Limiting co-repressor dynamics derails tissue formation, as documented for the TLE proteins that act down-stream of NOTCH, WNT, BMP, or growth factor signaling. Overexpression of TLE1 in mice inhibits forebrain development and induces formation of lung tumors ^41–43^, while in human patients, high TLE levels are seen in several cancers with progenitor-like transcriptional signatures and correlate with poor prognosis ^44–48^. In synovial sarcoma characterized by neural crest-like gene expression, TLE1 accumulation is now a major diagnostic biomarker ^46^. To explain the discrepancy between stable co-repressor binding to DNA *in vitro* and their dynamic behavior in cells, it has been proposed that co-repressors are actively dislodged from chromatin by transcriptional activators ^49–54^. How transcriptional activators, which are typically expressed at low levels, could dissociate an abundant complex that binds chromatin in a multivalent fashion remains to be explained.

Here, we show that proteasomes drive systemic degradation of repressive transcription factors to constantly eject co-repressors of the TLE/Groucho family from chromatin and thereby open genomic sites for activator binding. This regulation relies on the E3 ligase SCF^FBXL14^, which targets all WRPW- and EH1-motif containing repressive transcription factors dependent on their presentation by TLEs and independent of their actual identity. By continually cycling TLEs off chromatin, as accomplished by coordinated turnover of a protein family, cells circumvent the need for co-repressor displacement and instead routinely open promoters for activator binding. This dynamic regulation is essential for stem cells to translate developmental cues into accurate cell fate decisions, and it is disrupted by cancer mutations in *TLE1* that rewire gene expression similar to TLE1-overexpressing synovial sarcoma. We conclude that proteasomes systemically target repressive transcription factors to establish co-repressor dynamics required for lineage-specific gene expression and cell fate specification.

## Results

### TLE co-repressors recruit the E3 ligase SCF^FBXL14^

Co-repressors of the TLE-family are conserved regulators of gene expression that are recruited to genomic sites by multivalent interactions with nucleosomes and many repressive transcription factors ^30^. They act downstream of Notch, WNT, BMP or EGFR signaling, which are among the most important developmental pathways. Overexpression of TLEs disrupts tissue formation and is frequently observed in cancers characterized by precursor-like transcriptional signatures ^41–43,46,47,55^. These observations suggested that TLE proteins are subject to important regulatory mechanisms that remain to be discovered.

To isolate regulators of TLE function, we appended FLAG-epitope tags to the endogenous *TLE3* loci of 293T cells and determined its interactors by affinity-purification and mass spectrometry. TLE3 bound other TLE family members as well as expected co-repressor effectors, such as HDACs and GSK3 kinase ^56–60^. In addition, TLE3 engaged the E3 ligases SCF^FBXL14^, GID complex, and DTX4 (**Figure 1A**), while ubiquitylation enzymes suggested to regulate TLEs in other cell types were not detected ^61–63^. Among these E3 ligases, SCF^FBXL14^ quickly emerged as a candidate for important regulation: immunoprecipitation showed that its specificity factor FBXL14 efficiently bound all full-length TLEs, but barely any other protein (**Figure 1B**). Moreover, the interaction of FBXL14 with TLEs appeared to be specific, as it was disrupted by deleting a conserved extension to the leucine-rich repeats in FBXL14 (FBXL14^ΔCTE^) (**Figure S1A**). Similar to FBXL14, mutation a C-terminal extension to the leucine-rich repeats in the related FBXL17 had abolished binding events that were important for its role as a dimerization quality control enzyme ^64^.

**Figure 1:**
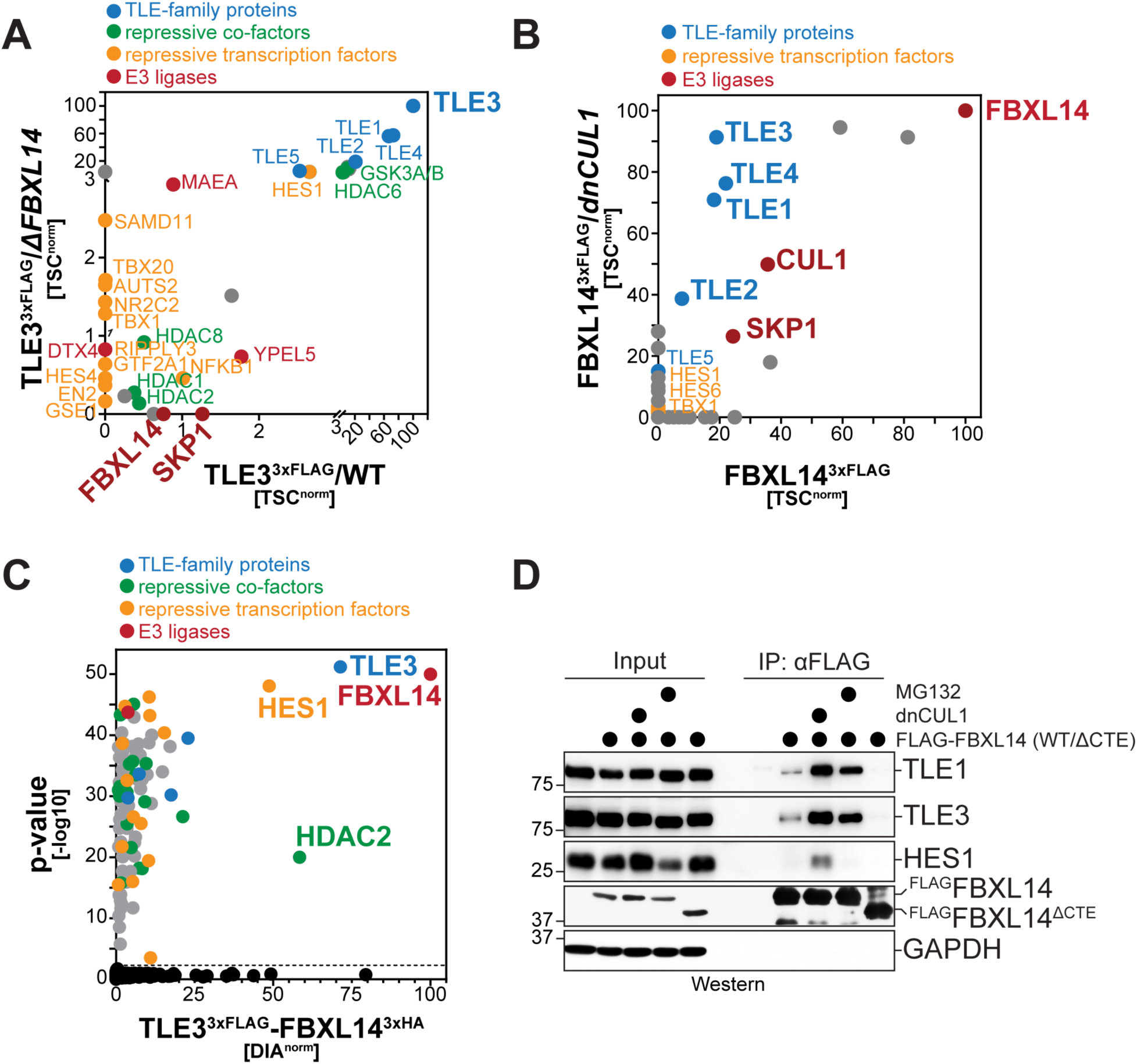
TLE co-repressors bind the E3 ligase SCF^FBXL14^. **A.** Endogenous TLE3 binds the CUL1-RING E3 ligase adaptor FBXL14. All TLE3 loci in either wildtype 293T cells (WT) or 293T cells deleted of *FBXL14* (*ΔFBXL14*) were fused to 3xFLAG epitope tags using CRISPR/Cas9-mediated genome editing. TLE3 was affinity-purified from cell lysates and binding partners were determined by mass spectrometry. **B.** FBXL14 binds all members of the TLE family of co-repressors. FBXL14^3xFLAG^ was affinity-purified from either WT cells or from cells expressing a dominant-negative variant of CUL1 (dnCUL1) and binding partners were determined by mass spectrometry. **C.** TLE3 incorporates FBXL14 into active co-repressor complexes. Sequential affinity-purifications of TLE3^3xFLAG^ and FBXL14^3xHA^ were analyzed for interactors by mass spectrometry. **D.** Validation of proteomic analyses by immunoprecipitation and Western blotting. Full-length FBXL14 or a variant lacking its carboxy-terminal extension (CTE) were affinity-purified from 293T cells and analyzed for endogenous binding partners by Western blotting using specific antibodies.

In contrast to abundant co-repressor effectors, we detected only few of its chromatin-targeting repressive transcription factors in affinity-purifications of endogenous TLE3. Having noted that TLE3 binds SCF^FBXL14^, we considered the possibility that ubiquitylation might displace transcription factors from TLEs and therefore repeated endogenous TLE3 immunoprecipitations in cells lacking *FBXL14*. Deletion of *FBXL14* broadly increased the association of TLE3 with several repressive transcription factors, while it did not affect recognition of other binding partners (**Figure 1A**). Affinity-purification of FBXL14 from cells with impaired E3 ligase activity similarly improved capture of transcription factors (**Figure 1B**). TLE3 therefore recruits SCF^FBXL14^, which in turn appears to limit the binding of TLE3 to chromatin-recruiting transcription factors.

To assess if TLE3 and FBXL14 bind transcription factors independently of each other or at the same time, we purified TLE3 and then enriched for complexes containing FBXL14 through a second immunoprecipitation step. Cells were supplemented with MLN4924 to block the degradation of associated proteins. These experiments revealed that 293T cells possess a major complex comprised of TLE3, FBXL14, the repressive transcription factor HES1, and HDAC2 (**Figure 1C**). Albeit at lower levels, TLE3-FBXL14 complexes also captured other HDACs, chromatin remodelers, as well as many repressive transcription factors. TLE3 therefore engages SCF^FBXL14^ at the same time as it binds transcription factors and effectors, indicating that TLE3 incorporates the E3 ligase into active co-repressor complexes.

Validating these proteomic studies, analysis of FBXL14-immunoprecipitates by Western blotting confirmed its association with TLE1 and TLE3 in a manner enhanced by dominant-negative CUL1 or proteasome inhibition (**Figure 1D**). E3 ligase or proteasome inhibition improved the interaction between FBXL14 and TLEs without impacting TLE levels. As we had seen by mass spectrometry, deletion of its CTE abolished the interaction of FBXL14 with TLE proteins. We also detected endogenous HES1 in complex with FBXL14, but only if substrate release from the E3 ligase was impaired by dominant-negative CUL1 (**Figure 1D**). As expected from its predominant association with a gene expression regulator, FBXL14 localized to the cell nucleus, but loss of TLE binding upon deletion of its CTE redirected FBXL14 to the cytoplasm (**Figure S1B**). These experiments confirmed that TLE co-repressors bind SCF^FBXL14^ and further suggested that these interactions occur with high efficiency and specificity.

SCF^FBXL14^ is an E3 ligase required for fly, zebrafish and frog development ^65–67^. Mirroring TLE1 overexpression ^43^, low levels of its specificity factor FBXL14 correlate with poor prognosis in cancers ^31^. SCF^FBXL14^ had previously been shown to target inositol-1,4,5-trisphosphate-gated calcium channels and regulators of the epithelial-mesenchymal transition ^68–72^. Its efficient interaction with TLEs suggested that SCF^FBXL14^ may also control co-repressor abundance or function, and we set out to reveal the basis for the underlying regulatory circuit.

### TLEs form compact hetero-dodecameric complexes with FBXL14-SKP1

As a first step towards understanding how SCF^FBXL14^ regulates co-repressors, we investigated the structural basis of the interaction between FBXL14 and TLE1, a family member that is crucial for cell fate specification and tumorigenesis ^43,73^. To facilitate structure determination, we generated ‘mini-TLE1’ by replacing the intrinsically disordered GP, CcN, and SP domains in TLE1 with a short Gly/Ser-rich linker to connect the N-terminal Q-domain to C-terminal WD40 repeats (**Figure 2A**). Previous work had identified the Q- and WD40-domains as critical for TLE function ^30,74^, and most inactivating TLE mutations obtained from genetic screens were found in these domains ^75^. Mini-TLE strongly bound purified FBXL14 (**Figure 2B; Figure S2A**), showing that it contained the motifs mediating E3 ligase recognition.

**Figure 2:**
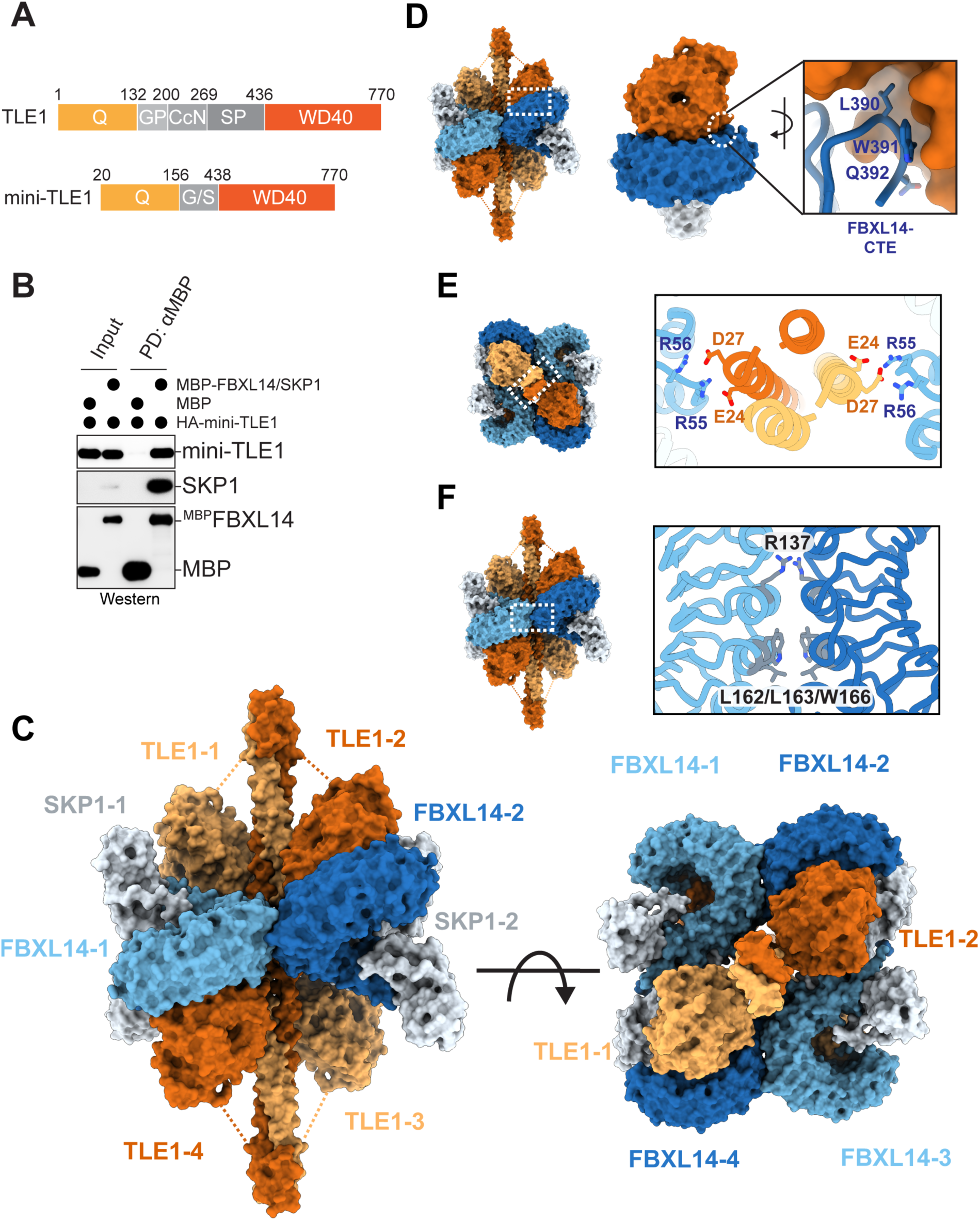
TLE1 forms compact oligomers with FBXL14-SKP1. **A.** Domain organization of either full-length TLE1 or ‘mini-TLE1’ that was used for structural analyses. **B.** Recombinant mini-TLE1 strongly binds purified FBXL14 *in vitro*, as assessed by αMBP-pulldown using MBP or ^MBP^FBXL14 as baits. Binding was analyzed by Western blotting. **C.** Cryo-EM structure of the complex between mini-TLE1 (orange), FBXL14 (blue), and SKP1 (grey). **D.** The LRR-domain of FBXL14 (blue) cradles the WD40 of TLE1 (orange). Inset shows the CTE of FBXL14 that nestles into a hydrophobic pocket on the periphery of the TLE1 WD40. **E.** A salt-bridge connects the FBXL14 LRR domain to the TLE1 Q-domain in the center of the structure. **F.** FBXL14 LRR-domains dimerize through backside binding.

Using cryo-EM, we obtained a structure of the complex between FBXL14, SKP1 and mini-TLE1 with an overall resolution of 3.9Å and a higher local resolution of 3.2Å achieved for a critical region encompassing FBXL14-SKP1 and the WD40 domains of TLE1 (**Figure 2C**; **Figure S2B-F**). At the center of this assembly is a TLE1 tetramer formed by head-to-head binding of dimeric building blocks. Each dimer unit depends on an extended coiled coil of the Q-domain, which is consistent with crystal structures of this region ^76^. Despite their connection to the Q-domain via a flexible linker, the WD40-repeats of mini-TLE1 localize around the central part of the Q-domain tetramer. The leucine-rich repeats (LRR) of the four FBXL14 subunits cradle each WD40, with the FBXL14-CTE fitting into a hydrophobic pocket of TLE1’s WD40 at the periphery of this interaction surface (**Figure 2D**). An Arg-clamp at the convex side of the LRRs forms additional salt-bridges with acidic residues of TLE1’s Q-domain (**Figure 2E**). This arrangement places all LRR domains of FBXL14 in a central plane perpendicular to the TLE1 coiled-coil. Locking this architecture in place, each FBXL14 molecule dimerizes in a back-to-back fashion with an FBXL14 subunit bound to the opposing TLE dimer (**Figure 2F**).

To validate the cryo-EM analysis, we generated FBXL14 and TLE1 variants with mutations at each interface. We first mutated residues at the surface of the FBXL14-LRRs that engage the TLE1 WD40 and found that the resulting variants failed to immunoprecipitate TLE1 from cells and were unable to bind TLE1 *in vitro* (**Figure 3A-C**). Reciprocal mutation of TLE1 residues similarly disrupted recognition of FBXL14 in cells and *in vitro* (**Figure S3A, B**). We next mutated residues in the CTE of FBXL14. Akin to deletion of the entire CTE, the FBXL14^386–391/A^ variant lost binding to TLE1 in cells and *in vitro* (**Figure 3D-F**). Mutation of FBXL14 residues that mediate the back-to-back interaction between LRRs also disrupted complex formation (**Figure 3G**). The association of LRRs is templated by TLE1 tetramerization, and TLE1 variants designed to prevent oligomerization were impaired in FBXL14 capture (**Figure S3C**). FBXL14-mutations that abolished the Arg-based salt-bridges with the TLE1 Q-domain did not abrogate complex assembly (**Figure S3D, E**). Thus, FBXL14, SKP1 and TLE1 form compact hetero-dodecameric complexes that are stabilized by interactions at multiple interfaces. Importantly, interface residues in TLE1 that are required for the interaction with FBXL14 are invariant in metazoan organisms expressing FBXL14, while they diverge in *C. elegans* that is missing a clear FBXL14 homolog (**Figure S3F**).

**Figure 3:**
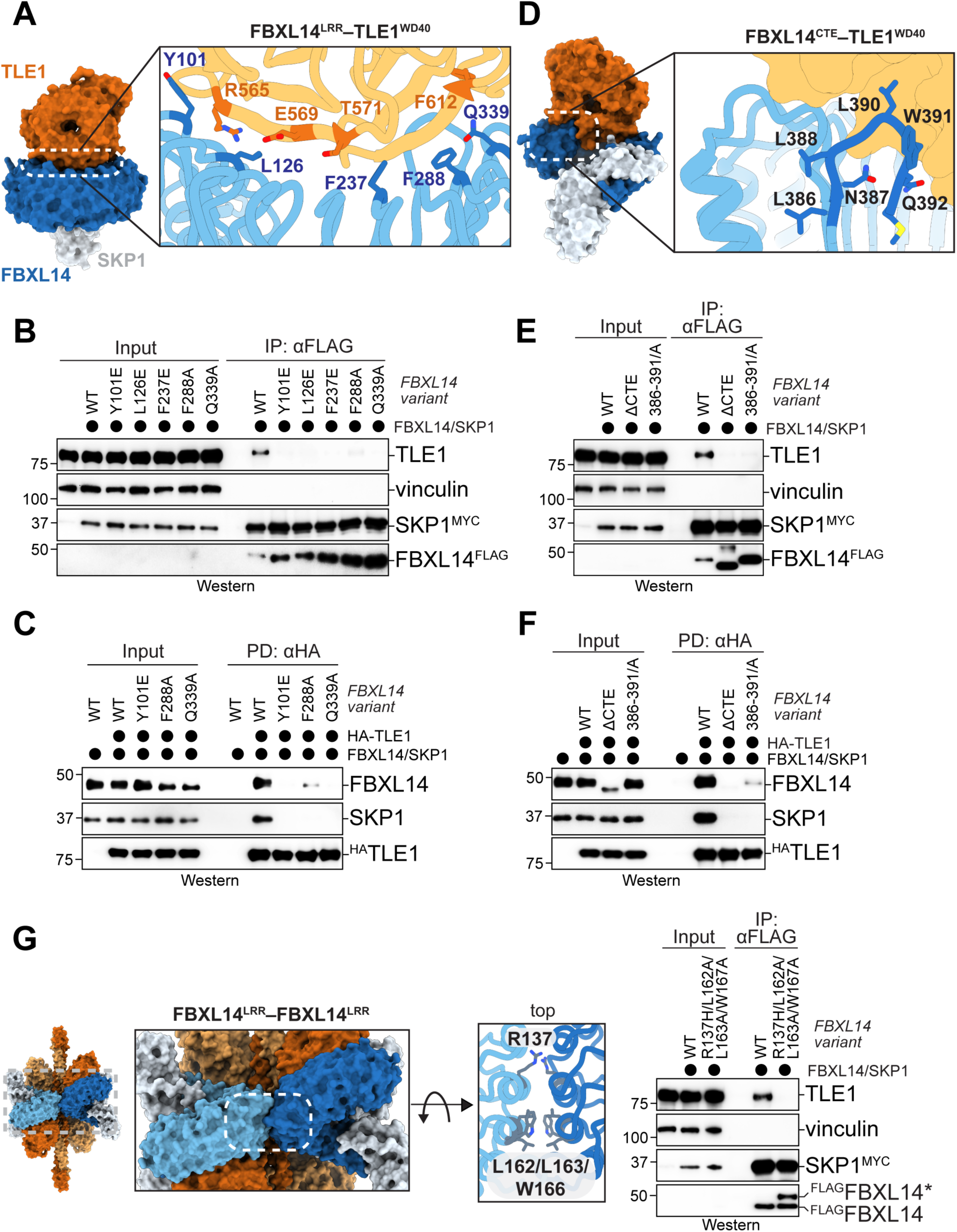
Multivalent interaction between TLE1 and FBXL14. **A.** Zoom into the interface between a TLE1 WD40 (orange) and an FBXL14 LRR domain (blue), highlighting conserved residues. **B.** Mutation of WD40/LRR-interface residues in FBXL14 disrupts binding to TLE1 in cells. WT FBXL14 or variants with altered interface residues were affinity-purified and bound endogenous TLE1 was detected by Western blotting. **C.** Mutation of WD40/LRR-interface residues in FBXL14 disrupts complex formation *in vitro*. WT- or mutant FBXL14 variants were immobilized and incubated with recombinant TLE1. Binding was visualized after gel electrophoresis by Western blotting. **D.** Structural depiction of the interaction between the FBXL14-CTE and a hydrophobic pocket at the periphery of the TLE1 WD40. Important residues are highlighted in the inset. **E.** Mutation of the five CTE residues in FBXL14 to Ala disrupts interaction with TLE1 in cells, as shown by immunoprecipitation of WT-FBXL14, FBXL14^ΔCTE^ (entire CTE deleted), or FBXL14^386–291/A^ (all CTE residues mutated to Ala). Co-purifying endogenous TLE1 was detected by Western blotting. **F.** Reciprocal immunoprecipitation of TLE1^3HA^ *in vitro* shows loss of binding to recombinant FBXL14 mutants that either lack or have a mutated CTE. **G.** Mutation of residues at the interface between two FBXL14 LRR domains disrupts formation of the TLE1-FBXL14 complex in cells. WT- or mutant FBXL14^3FLAG^ were immunoprecipitated and co-purifying endogenous TLE1 was detected by Western blotting.

In addition to providing structural insight into a conserved E3 ligase-corepressor complex, our findings immediately suggested TLE1-functions under control of ubiquitylation. The TLE1 Q-domain encircled by FBXL14 molecules also recognizes TCF transcription factors that are important for WNT signaling ^76^. Thus, TLE proteins may either bind TCF proteins or FBXL14 but not both at the same time, and indeed, we did not find TCFs in sequential FBXL14-TLE3 affinity-purifications (**Figure 1D**). By contrast, TLEs engage repressive transcription factors with WRPW or EH1 motifs through their WD40 domains, which remain accessible in the presence of FBXL14. Accordingly, we had observed ternary complexes formed by TLE3, FBXL14, and such transcription factors (**Figure 1C**). SCF^FBXL14^ therefore likely regulates TLEs that are bound to WRPW-or EH1-motif transcription factors, a large family of proteins that stabilize the association of co-repressors with specific sites on chromatin.

### SCF^FBXL14^ targets TLE-bound transcription factors for degradation

By modelling CUL1, RBX1, and a ubiquitin-charged E2 enzyme into our structure, we noticed a gap of ∼60Å between TLE1 and the catalytic center of SCF^FBXL14^ (**Figure 4A**). Given the compact nature of the complex, SCF^FBXL14^ is unlikely to efficiently ubiquitylate TLE1 in this configuration. Indeed, *FBXL14* deletion did not protect TLE1 stability reporters from degradation (**Figure S4A**), and loss of *FBXL14* did not increase the abundance of endogenous TLE1 in stem cells (**Figure S4B**). These observations do not exclude the possibility that SCF^FBXL14^ may target TLE proteins in response to signals that change complex composition or architecture.

**Figure 4:**
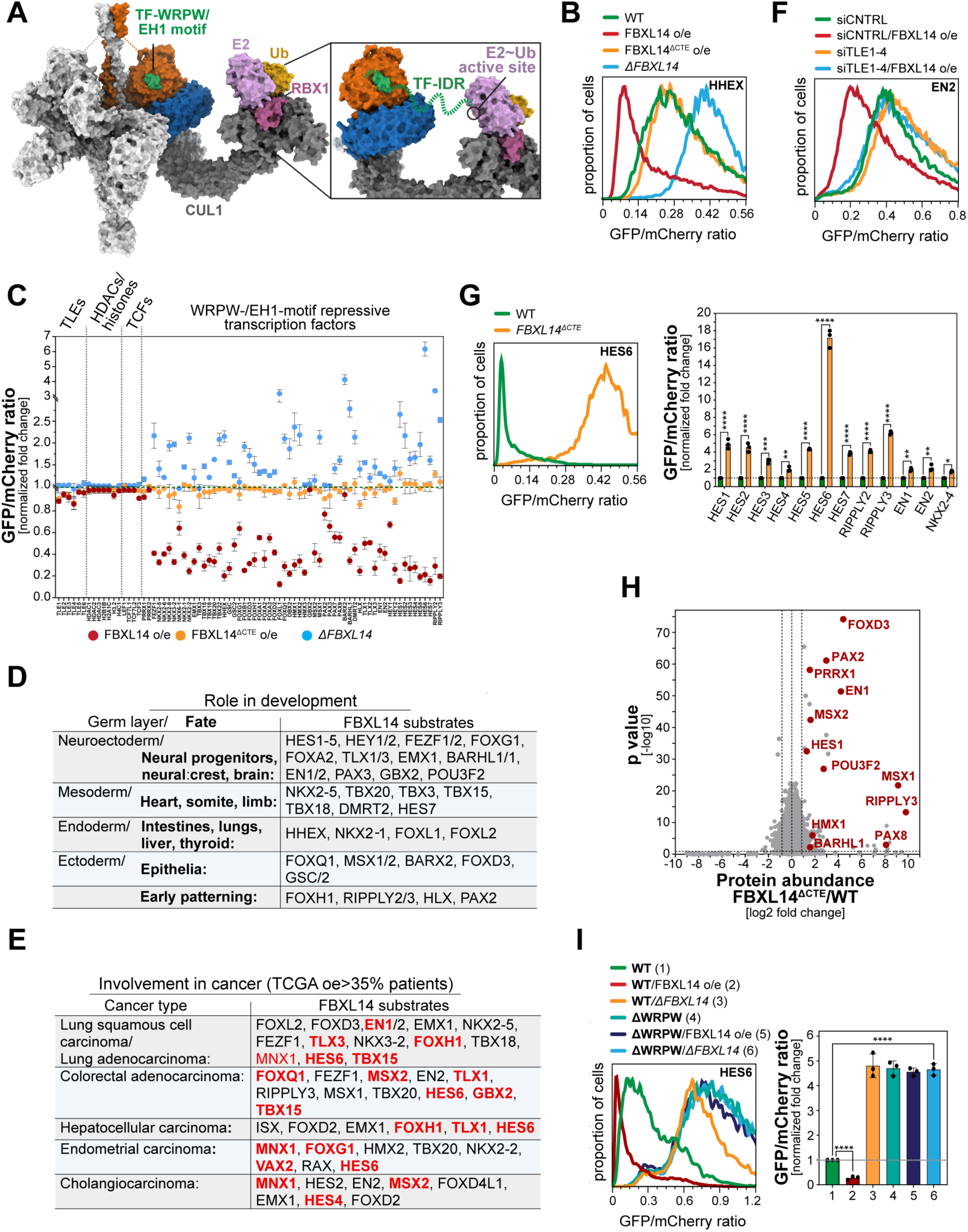
TLEs present repressive transcription factors for ubiquitylation by SCF^FBXL14^. **A.** Structural model of the full SCF^FBXL14^ bound to TLE1. For simplicity, only a single CUL1/RBX1 module is shown. In green, a TF peptide was modelled into its binding position on the TLE1 WD40 domain, based on X-ray structures (PDBs: 1LDK, 2CE9) and AlphaFold modeling. The inset shows how disordered regions in proximity to the WRPW- or EH1-motifs in transcription factors could bridge the gap from the WD40 domain to the active site of a ubiquitin-charged E2 enzymes. **B.** Flow cytometry measurements of stability reporters of the HHEX transcription factors in WT cells (green); cells overexpressing WT-FBXL14 (red); cells overexpressing TLE-binding deficient FBXL14^ΔCTE^ (orange); or cells deleted of *FBXL14* (*ΔFBXL14*; blue). **C.** WRPW- and EH1-motif transcription factors are targeted by FBXL14 dependent on TLEs. Reporter assays described above were performed for indicated proteins and change in protein abundance relative to WT cells was quantified. n=3 independent experiments. **D.** SCF^FBXL14^-substrates with important roles in tissue formation. **E.** SCF^FBXL14^ substrates overexpressed in the TCGA. Red targets have been classified as oncogenes. **F.** Depletion of all full-length TLEs (TLE1-TLE4) prevents transcription factor degradation induced by FBXL14 overexpression. Note that TLEs were partially depleted by siRNAs; complete deletion of four *TLE* genes was not viable. Stability reporters were monitored in the indicated cell lines by flow cytometry, as described above. **G.** Deletion of the CTE in the endogenous *FBXL14* loci to create cells that cannot form any TLE-FBXL14 complexes stabilizes both WRPW- and EH1-motif containing transcription factors, as shown for several stability reporters using flow cytometry. Quantification of n=3 independent experiments is shown on the right. **H.** Deletion of the CTE in endogenous *FBXL14* of iPSCs results in accumulation of several WRPW- and EH1-motif containing repressive transcription factors, as determined by data-independent acquisition mass spectrometry. **I.** Deletion of WRPW- or EH1-motifs protects transcription factors from degradation through SCF^FBXL14^. Stability reporters for the WRPW-motif transcription factor HES6 and the EH1-motif transcription factor EN2 were analyzed by flow cytometry in either WT cells; cells overexpressing FBXL14; or *ΔFBXL14* cells. Quantification of three independent experiments is shown on the right. In all experiments, data are represented from n=3 independent experiments +/- SD.

While TLEs were positioned distant from the catalytic center of the E3 ligase, the WRPW-or EH1-engaging surface of their WD40 domain was oriented towards the E2 enzyme bound to SCF^FBXL14^ (**Figure 4A**). As WRPW and EH1 motifs are typically embedded in regions of intrinsic disorder (**Figure S4C**), these transcription factors may be able to bridge the gap between their binding site and the catalytic center of SCF^FBXL14^. To test this notion, we measured the stability of a large collection of WRPW- or EH1 motif transcription factors using flow cytometry ^17,77–79^. Strikingly, all transcription factors that engage the WD40 repeats of TLEs were depleted upon FBXL14 overexpression and stabilized by *FBXL14* deletion (**Figure 4B, C**). Overexpression of FBXL14^ΔCTE^, which does not bind TLEs, did not elicit broad transcription factor turnover, and histones, TCFs, or HDACs that associate with TLEs outside of their WD40 repeats were not affected by changes in FBXL14 abundance or function (**Figure 4C**). The degradation of WRPW-and EH1-motif transcription factors was blocked by inhibition of Cullin-RING E3 ligases or proteasomes (**Figure S4D**). Thus, SCF^FBXL14^ targets all tested WRPW- and EH1-motif containing transcription factors, irrespective of their actual identity, for degradation. These SCF^FBXL14^ substrates include crucial regulators of cell fate specification and organ development (**Figure 4D**), and many are overexpressed in cancers or have been characterized as potent oncogenes (**Figure 4E**).

The apparently systemic degradation of WRPW- and EH1-motif containing transcription factors could be explained if binding to the TLE WD40 domain, a property shared by all these proteins, would be required and sufficient for recognition by SCF^FBXL14^. In line with this notion, depletion of all full-length TLEs abrogated the ability of FBXL14 to broadly elicit transcription factor degradation (**Figure 4F; Figure S4E**). Deletion of the CTE in endogenous *FBXL14* to create cells that cannot form TLE-FBXL14 complexes also protected stability reporters from degradation and increased the abundance of several endogenous WRPW- and EH1-motif transcription factors (**Figure 4G, H; Figure S4F**). As a consequence of failed turnover, multiple transcription factors became trapped in complexes with TLEs if cells were unable to recruit SCF^FBXL14^ to these co-repressors (**Figure S4G**). Finally, deletion of WRPW- or EH1-motifs that mediate binding to TLEs protected all tested transcription factors from their degradation dependent on SCF^FBXL14^ (**Figure 4I; Figure S4H**).

Together, these results showed that SCF^FBXL14^ broadly targets WRPW- and EH1-motif transcription factors for proteasomal degradation. This regulation is dependent on co-repressor presentation, but independent of protein identity, resulting in the systemic turnover of a family of repressive transcription factors that recruit the co-repressors to specific sites on chromatin. Many of these SCF^FBXL14^ substrates have critical roles in tissue formation or have been characterized as potent oncogenes ^30,75,80,81^, suggesting that systemic transcription factor degradation may play an important role in development and disease.

### SCF^FBXL14^ targets chromatin-bound transcription factors

To understand the physiological role of systemic transcription factor turnover, we dissected where and when cells target these proteins for degradation. SCF^FBXL14^ belongs to the family of CUL1-RING E3 ligases that often recognize proteins marked for degradation by posttranslational modifications, such as phosphorylation ^82,83^. To test if transcription factors were subject to similar regulation, we first reconstituted substrate binding by SCF^FBXL14^ using recombinant proteins. As expected from our structure, transcription factors did not interact with FBXL14 directly, but were recruited to the E3 ligase through TLE1 (**Figure S5Α**). Transcription factor variants that lacked WRPW or EH1 motifs and cannot engage TLE1 were not recognized by FBXL14. This observation validated that SCF^FBXL14^ recognizes transcription factors presented by TLEs, but posttranslational modifications did not seem to be required to mediate such interactions.

Based on these observations, we synthesized peptides that contained WRPW- or EH1-motifs as well as Lys residues as potential ubiquitin acceptor sites. In line with our binding studies, the degron peptides were ubiquitylated by SCF^FBXL14^ in a TLE1-dependent, but phosphorylation-independent, manner (**Figure 5Α**). Mutation of R534 in the WD40 repeat of TLE1, which abrogates recognition of all tested WRPW- or EH1-motifs ^42,75^, prevented ubiquitylation through SCF^FBXL14^ (**Figure 5A**). In a similar manner, mutations in TLE1 or FBXL14 that impaired the interaction between these proteins reduced transcription factor modification (**Figure 5B**; **Figure S5B, C**). As seen for other SCF-family E3 ligases ^84^, SCF^FBXL14^ acted most effectively in the presence of distinct E2 enzymes for chain initiation and elongation, respectively (**Figure S5D**), and it decorated its targets with K48-linked chains that mediate proteasomal degradation (**Figure S5E**). SCF^FBXL14^ therefore ubiquitylates transcription factors dependent on their presentation by TLEs, but without requiring posttranslational modifications.

**Figure 5:**
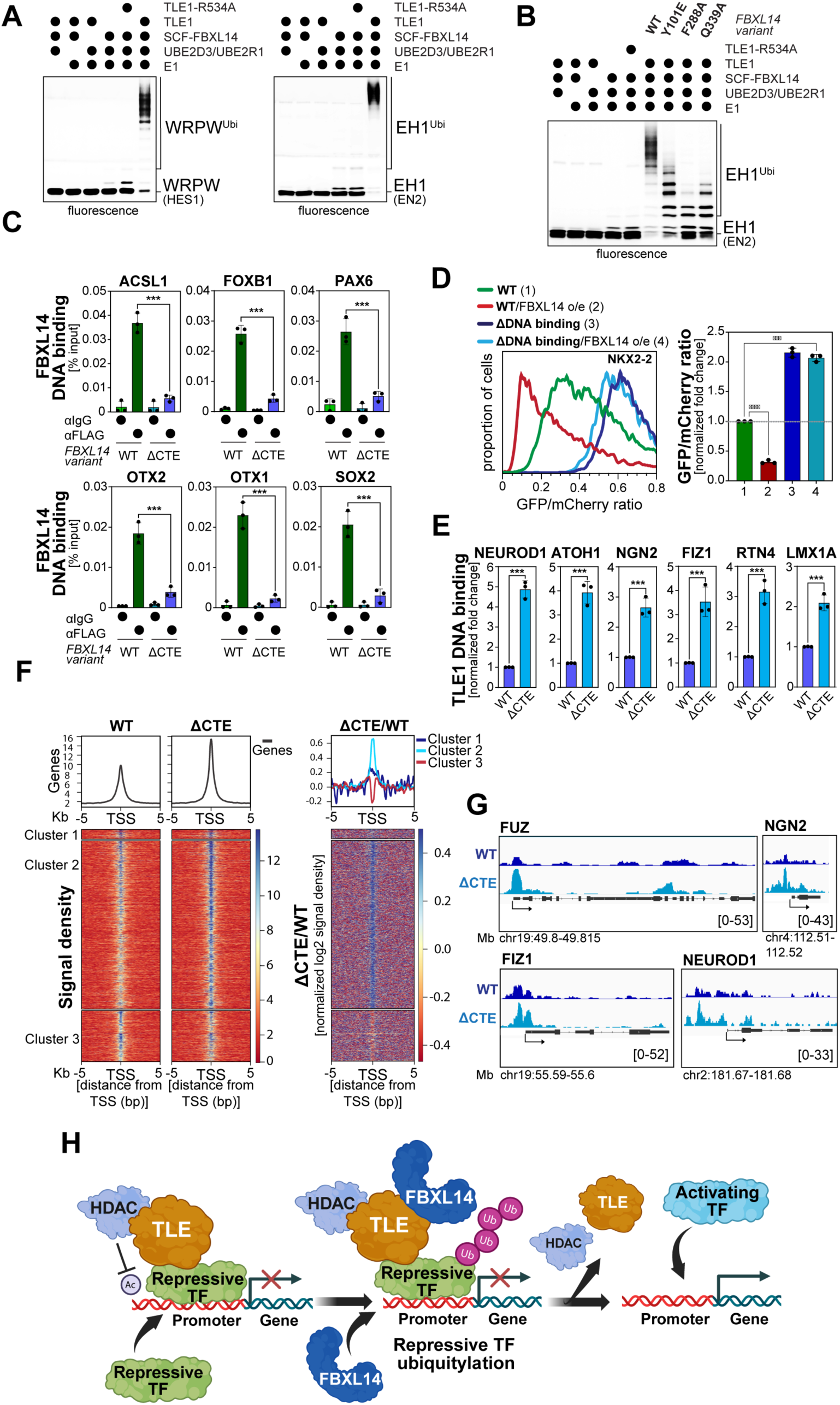
SCF^FBXL14^ releases TLE proteins from chromatin. **A.** SCF^FBXL14^ ubiquitylates transcription factors dependent on their presentation by TLEs. TAMRA-labeled WRPW- and a EH1-motif peptides derived from SCF^FBXL14^-targets (HES1, EN2) were incubated with recombinant E1, E2 (UBE2D3, UBE2R1), SCF^FBXL14^, and ubiquitin. As indicated, either WT-TLE1 or TLE1^R534A^ was added. Ubiquitylation was monitored by gel electrophoresis and fluorescence detection. **B.** Mutation of FBXL14-residues at the interface with TLE1 impedes ubiquitylation of transcription factor degrons. A TAMRA-labeled EH1 motif peptide derived from the SCF^FBXL14^ substrate EN2 was incubated with recombinant E1, E2 (UBE2D3, UBE2R1), SCF^FBXL14^, TLE1 and ubiquitin. As indicated, either WT-FBXL14 or FBXL14 variants impaired in TLE binding were added. TLE1^R534A^ was used as an additional control. Ubiquitylation was monitored as described above. **C.** FBXL14 is recruited to known TLE-binding sites on chromatin. iPSCs stably expressing endogenously FLAG-tagged FBXL14 or FBXL14^ΔCTE^, respectively, were subjected to αIgG (control) or αFLAG chromatin-immunoprecipitation, and co-purifying DNA was detected by qPCR. DNA sequences were selected based on known TLE-binding sites ^103^. **D.** Deletion of DNA binding motifs in repressive transcription factors protects them against degradation by SCF^FBXL14^. Stability of an example transcription factor, NKX2-2, was monitored by flow cytometry, as described above. Either WT-NKX2-2 or a variant with its DNA-binding domain deleted were analyzed, and FBXL14 was overexpressed as indicated. **E.** SCF^FBXL14^ restricts chromatin binding of TLEs. Binding of endogenous TLE1 to known sites on chromatin was investigated in iPSCs either expressing WT-FBXL14 or FBXL14^ΔCTE^ from the endogenous *FBXL14* loci, using Cut&Run-qPCR analysis. In all experiments, data are represented from n=3 independent experiments +/- SD. **F.** Cut&Run-seq of chromatin association of endogenous TLE1 in either WT-iPSCs or iPSCs that expressed FBXL14^ΔCTE^ from the endogenous *FBXL14* loci. Right panel shows a difference map between both experiments. **G.** Genome browser tracks of TLE1 chromatin binding for example binding sites in either WT-iPSCs or iPSCs that expressed FBXL14^ΔCTE^ from the endogenous *FBXL14* loci. **H.** Model of recurrent TLE recruitment to chromatin, transcription factor degradation through SCF^FBXL14^ and TLE release.

Previous studies found that TLEs bind transcription factors more stably on chromatin than in solution, dependent on conformational changes in TLEs that are induced by their association with nucleosomes ^30^. If TLEs indeed present transcription factors to SCF^FBXL14^, as revealed by our structural and biochemical investigations, then a more stable interaction between these partners could result in localized degradation on chromatin. In line with this notion, chromatin immuno-precipitations (ChIP) found FBXL14 to bind the same genomic sites that are known to be occupied by TLE proteins (**Figure 5C**). Deletion of its CTE strongly diminished FBXL14’s ability to engage chromatin, showing that the E3 ligase is recruited to DNA through its interactions with TLEs. Conversely, deleting DNA-binding domains in transcription factors to block their chromatin association also prevented their SCF^FBXL14^-dependent turnover (**Figure 5D**), without impacting their nuclear localization (**Figure S5F**). Rather than relying on posttranslational modifications, SCF^FBXL14^ therefore appears to target transcription factors that form complexes with TLEs on DNA.

As transcription factors lock TLEs onto chromatin, their degradation might release the co-repressors from DNA. We tested this notion by performing Cut&Run-qPCR of endogenous TLE1 in WT- and *FBXL14^ΔCTE^* iPSCs that were induced to undergo neuronal differentiation and hence were primed to exchange co-repressors with lineage-specific activators. *FBXL14^ΔCTE^* iPSCs that cannot ubiquitylate transcription factors retained TLE1 on chromatin (**Figure 5E**). Whole genome Cut&Run sequencing showed that SCF^FBXL14^-inhibition led to an accumulation of TLE1 across many genomic loci (**Figure 5F, G**), most of which were transcription start sites that overlapped with regions that during differentiation are bound by lineage-specific activators (**Figure S5G**). The broad effects of SCF^FBXL14^ inactivation onto chromatin binding of TLEs highlighted the ability of this E3 ligase to target many transcription factors, as expected for systemic protein degradation. Moreover, SCF^FBXL14^ acts by limiting the amplitude of the TLE1 signal, and not by preventing spreading of TLE1 to non-physiological sites, which highlights that the E3 ligase restricts physiological transcription factor function rather than targeting aberrant proteins. We conclude that TLEs broadly present DNA-bound transcription factors for SCF^FBXL14^-dependent ubiquitylation and degradation, which releases TLEs from chromatin and routinely opens transcription start sites for potential activator binding (**Figure 5H**).

### Transcription factor degradation is required for cell fate specification

Given the importance of TLEs for developmental signaling, we wished to determine if systemic transcription factor degradation is required for differentiation events that depend on exchange of co-repressors with lineage-specific activators. We therefore subjected WT- and *FBXL14^ΔCTE^-*iPSCs to neural conversion, as described above, and followed their differentiation into either neuronal or neural crest progenitors. RNAseq of differentiating iPSCs revealed that FBXL14 inactivation downregulated nearly all markers of neural progenitors (**Figure 6A**). Rather than remaining pluripotent or dying, *FBXL14^ΔCTE^-*iPSCs upregulated the expression of neural crest markers, indicative of a switch in cell fate. We confirmed the change from neuronal to neural crest fate for select markers by qRT-PCR (**Figure 6B**), microscopy (**Figure 6C**), and Western blot analysis (**Figure S6A**). We made similar observations when we acutely depleted FBXL14 from iPSCs (**Figure S6B-D**) or engineered hESCs to express endogenous *FBXL14^ΔCTE^* (**Figure S6E-G**). Rescue experiments highlighted the specificity of these depletions and showed that FBXL14 must interact with TLEs to control cell fate: re-expression of WT-FBXL14, but not any TLE-binding mutant, restored neuronal differentiation in *FBXL14^ΔCTE^-*iPSCs (**Figure S6H**).

**Figure 6:**
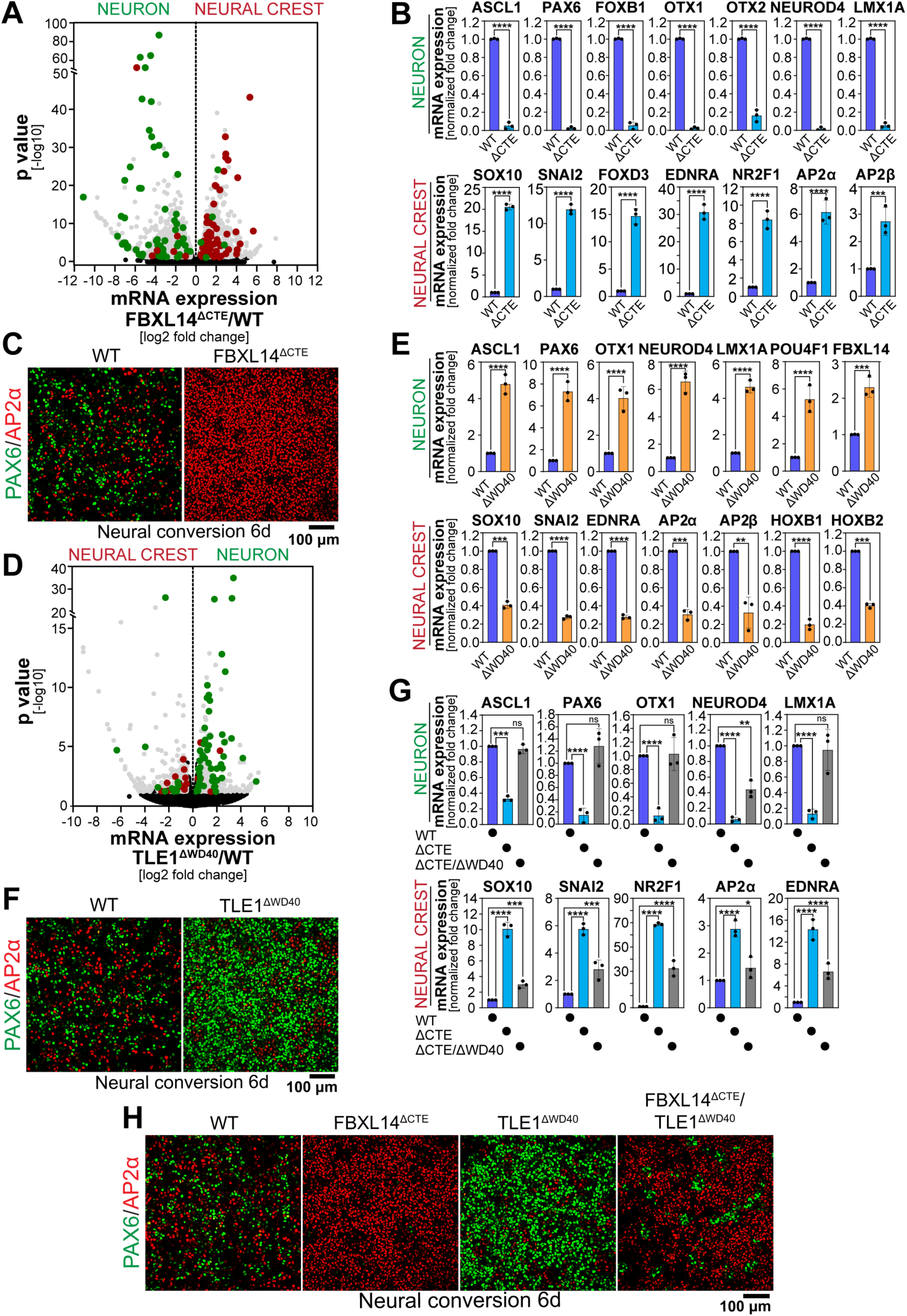
SCF^FBXL14^ controls cell fate in a TLE-dependent manner. **A.** RNAseq experiments in either WT-iPSCs or *FBXL14^ΔCTE^::*iPSCs subjected to neural conversion show downregulation of neural markers and simultaneous upregulation of neural crest markers if FXBL14 cannot bind TLEs. **B.** Confirmation of the cell fate switch from a neural to a neural crest fate upon loss of TLE binding by FBXL14 in iPSCs subjected to neural conversion, as assessed by qRT-PCR. **C.** Confirmation of the cell fate switch from a neural to a neural crest fate upon loss of TLE binding by FBXL14 in iPSCs subjected to neural conversion, as assessed by immunofluorescence microscopy against a neural (PAX6; green) and neural crest (AP2α; red) marker. **D.** RNAseq experiments in either WT-iPSCs or *TLE1^ΔWD40^::*iPSCs subjected to neural conversion show downregulation of neural crest markers and simultaneous upregulation of neural markers if TLEs cannot bind WRPW- or EH1-motif containing repressive transcription factors. **E.** Confirmation of the cell fate switch from a neural crest to a neural fate upon loss of transcription factor binding by TLE1 in iPSCs subjected to neural conversion, as assessed by qRT-PCR. **F.** Confirmation of the cell fate switch from a neural crest to a neural fate upon loss of transcription factor binding by TLEs in iPSCs subjected to neural conversion, as assessed by immunofluorescence microscopy against a neural (PAX6; green) and neural crest (AP2α; red) marker. **G.** SCF^FBXL14^ controls cell fate through TLEs. WT-iPSCs, *FBXL14^ΔCTE^*::iPSCs, or *FBXL14^ΔCTE^; TLE1^ΔWD40^*::iPSCs were subjected to neural conversion. Expression of neural markers (top row) or neural crest markers (bottom row) was assessed by qRT-PCR. In all experiments, data are represented from n=3 independent experiments +/- SD. **H.** WT-iPSCs, *FBXL14^ΔCTE^*::iPSCs, or *FBXL14^ΔCTE^; TLE1^ΔWD40^*::iPSCs were subjected to neural conversion. Expression of neural markers (PAX6; green) or neural crest markers (AP2α; red) was assessed by immunofluorescence microscopy.

To further assess whether SCF^FBXL14^ acts through TLEs, we inactivated TLE1 in iPSCs by excising the WD40 domain from its endogenous loci. *TLE1^ΔWD40^-*iPSCs still possess other TLE proteins and hence reduce, but do not eliminate, gene repression. *TLE1^ΔWD40^-*iPSCs subjected to neural conversion behaved opposite to SCF^FBXL14^: TLE1 inactivation enhanced neuronal differentiation while inhibiting neural crest specification, as measured by RNAseq, qRT-PCR, and microscopy (**Figure 6D-F**). These observations are consistent with results in *ΔTLE1* mice that showed delayed appearance of neural crest-derived melanocytes ^73^. We then inactivated FBXL14 and TLE1 at the same time (*FBXL14^ΔCTE^; TLE1^ΔWD40^*) and, strikingly, found that *TLE1* inactivation restored neuronal and neural crest specification of *FBXL14^ΔCTE^* iPSCs to wildtype levels (**Figure 6G, H**). SCF^FBXL14^ therefore controls cell fate by limiting TLE-dependent gene repression, showing that systemic degradation of repressive transcription factors and the constant release of TLEs from chromatin are critical for lineage-specific gene expression and cell fate specification.

### Cancer mutations in TLE1 impede cell fate specification

The transcriptional rewiring of *FBXL14^ΔCTE^* iPSCs was reminiscent of synovial sarcoma, a cancer that is diagnosed using TLE1 as a biomarker and characterized by a neural crest-like expression signature ^85^. Other tumors with neural crest-like gene expression also show high TLE1 levels ^86,87^, but whether disrupting systemic transcription factor degradation could result in similar phenotypes as TLE1 overexpression is not known. Intriguingly, we found by searching the TCGA that residues in TLE1 at the interface with FBXL14 are mutated in cancer (**Figure 7A**). These TLE1 cancer variants failed to engage FBXL14 in cells (**Figure 7B**), and as shown for TLE1^R610S^, did not bind FBXL14 *in vitro* (**Figure 7C**). TLE1^R610S^ was also strongly impaired in supporting transcription factor ubiquitylation through SCF^FBXL14^ (**Figure 7D**).

**Figure 7:**
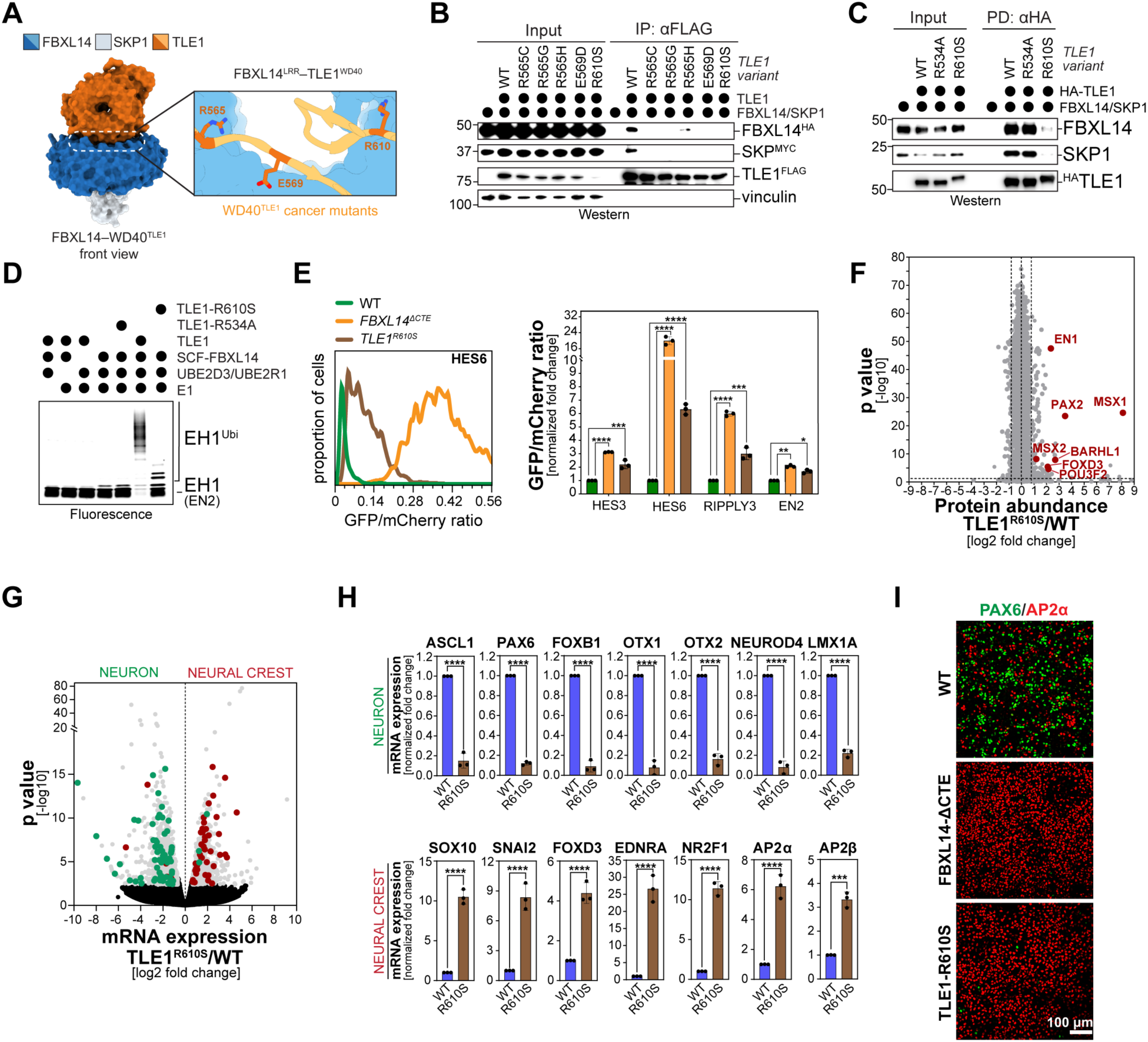
Cancer mutations in TLE1 disrupt regulation of cell fate through SCF^FBXL14^. **A.** Cancer mutations in *TLE1* reported in the TCGA localize to the interface with FBXL14’s LRR domain. **B.** Cancer mutations in TLE1 disrupt binding to FBXL14, as revealed by affinity-purification of TLE1^FLAG^. Co-purifying FBXL14 was detected by Western blotting. **C.** The TLE1^R610S^ variant is unable to bind FBXL14 *in vitro*. Recombinant TLE1^HA^ variants (WT-TLE1; TLE1^R534A^, defective in transcription factor binding; or TLE1^R610S^, mutated in cancer) were immobilized on αHA-resin and incubated with either purified FBXL14/SKP1 complexes. Binding of FBXL14 was detected after pulldown by Western blotting. **D.** The TLE1^R610S^ variant found in cancer is strongly impaired in promoting transcription factor ubiquitylation *in vitro*. A TAMRA-labeled EH1 motif peptide derived from the SCF^FBXL14^ substrate EN2 was incubated with recombinant E1, E2 (UBE2D3, UBE2R1), SCF^FBXL14^, TLE1 and ubiquitin. As indicated, either WT-TLE1 or TLE^1R610S^ were added. TLE1^R534A^ was used as an additional control. Ubiquitylation was monitored as described above. **E.** Cancer mutations in TLE1 stabilize SCF^FBXL14^ substrates. Stability reporters of various SCF^FBXL14^ substrates were investigated by flow cytometry in WT-iPSCs or iPSCs carrying endogenous *FBXL14^ΔCTE^* or *TLE1^R610S^* mutations. Left panel shows a single example for the WRPW-motif transcription factor HES6, while right panel shows quantification of n=3 independent experiments for both WRPW- and EH1-motif containing transcription factors. **F.** Cancer mutations in TLE1 increase abundance of oncogenic transcription factors. Protein abundance in either WT-iPSCs or iPSCs expressing endogenous TLE1^R610S^ was determined by DIA mass spectrometry. **G.** Cancer mutations in TLE1 rewire cell fate specification. WT-iPSCs or iPSCs expressing endogenous TLE1^R610S^ were subjected to neural specification and analyzed by RNAseq. Markers for neural differentiation are shown in green; markers for neural crest specification are shown in red. **H.** Confirmation of the neural to neural crest switch in cell fate caused by the TLE1^R510S^ mutation, as shown by qRT-PCR. In all experiments, data are represented from n=3 independent experiments +/- SD. **I.** Confirmation of the neural to neural crest switch in cell fate caused by the TLE1^R510S^ mutation, as shown by immunofluorescence microscopy of a neural marker (PAX6; green) and a neural crest marker (AP2α; red).

To assess cancer variants at the endogenous level, we introduced the R610S mutation into the *TLE1* loci of iPCSs. Flow cytometry assays revealed stabilization of many transcription factor targets of SCF^FBXL14^, albeit to a less dramatic extent as seen upon *FBXL14* deletion (**Figure 7E**). Whole cell proteomics accordingly showed that several endogenous SCF^FBXL14^-substrates, including the oncogenic EN1, MSX2, or POU3F2, increased in abundance in *TLE1^R610S^* iPSCs (**Figure 7F**). When subjected to neural conversion, *TLE1^R610S^-*iPSCs were strongly biased towards a neural crest fate, which was seen by RNAseq, qRT-PCR and microscopy (**Figure 7G-I**). Thus, *TLE1* mutations in cancer disrupt recognition of many transcription factors by SCF^FBXL14^ and divert differentiation to neural crest-like progenitors. This change in gene expression is highly similar to synovial sarcoma, which suggests that systemic transcription factor degradation ensures both tissue formation and homeostasis.

## Discussion

Proteasomes have long been known to broadly enable cells to initiate gene expression, but most work had focused on specific transcription factors that must be released from inhibitory complexes or need to be turned over for RNA polymerase to move from initiation to elongation states ^17,88,89^. These studies did not uncover the general function of protein turnover in enabling gene expression. Here, we show that systemic degradation of repressive transcription factors constantly releases TLE co-repressors from DNA, thereby freeing many transcription start sites for potential activator binding and enabling cells to initiate gene expression programs essential for cell fate specification. Central to this regulation is the E3 ligase SCF^FBXL14^, which systemically targets the WRPW- and EH1-motif transcription factors that recruit TLE co-repressors to chromatin sites critical for developmental gene expression.

SCF^FBXL14^ only acts on transcription factors that can bind DNA, suggesting that it predominantly functions by releasing TLEs. In line with this notion, ChIP experiments found FBXL14 at the same genomic sites as TLEs, dependent on FBXL14’s ability to integrate into co-repressor complexes. Previous work had revealed that TLEs bind transcription factors much more stably on DNA ^30^, and our structures now show that these interactions broadly present transcription factors to SCF^FBXL14^ for ubiquitylation and degradation. Following degradation of transcription factors and subsequent release of TLEs from chromatin, TLEs likely dissociate from FBXL14 and are thus free to bind new transcription factors to be delivered to new genomic sites. Our work therefore suggests that TLEs constantly cycle on and off chromatin. While more work is needed to understand how SCF^FBXL14^ recognizes transcription factors on chromatin, these results show that gene repression through TLEs is much more dynamic than previously appreciated.

Systemic degradation of repressive transcription factors has important implications for gene expression control. When developmental signals call for initiation of lineage-specific transcription, activators must gain access to sites on chromatin that are kept off in stem cells by co-repressors. Previous models suggested that activators displace repressors, potentially enabled by posttranslational modifications that stabilize activators or reduce the co-repressor affinity to chromatin. However, activators accumulate to much lower levels than co-repressors, and the latter associate with chromatin through multivalent interactions that are rarely disrupted by select posttranslational modifications. Providing an alternative model, our work suggests that activators do not need to displace repressors from chromatin. Rather, the constant release of co-repressors from chromatin by systemic transcription factor turnover constantly frees transcription start sites for activator binding. While this regulation is energetically costly and requires cells to continuously synthesize new transcription factors, it endows cells with the flexibility required for the rapid cell fate specification events that characterize early metazoan development.

The targets of SCF^FBXL14^-dependent degradation include crucial developmental regulators, such as HES proteins that control stem cell pluripotency and neurogenesis ^90^; TLXs that are key to patterning of the central nervous system ^91^; FOXG1 that is crucial for telencephalon development ^92^; the FOXA family of transcription factors that drive endoderm formation ^93^; FOXQ1 that controls epithelial differentiation ^94^; or NKX2 proteins that guide lung or heart development ^95^. Inactivation of these SCF^FBXL14^ targets leads to a wide range of developmental diseases, while gain-of-function mutations or overexpression have been linked to tumorigenesis. Accordingly, we found that SCF^FBXL14^ is a central regulator of stem cell differentiation whose deletion causes a striking switch in cell fate. Downregulation of FBXL14 homologs in flies, frogs or zebrafish also resulted in patterning defects ^65,66,96,97^, suggesting that similar to its structure, its developmental functions are conserved. By contrast, overexpression of TLE1 induces lung cancer in mice ^43^, and it correlates with poor patient outcomes ^98^. In synovial sarcoma, where TLE1 is used as biomarker ^46,47^, high TLE1 levels induce a neural-crest like gene expression program ^44,45^. Expanding on these results, we found that *TLE1* mutations in cancer that disrupt FBXL14 binding rewire differentiation towards a neural crest-like fate. Systemic transcription factor degradation therefore not only controls developmental gene expression but also protects organisms from tumorigenesis.

Similar to TLEs, Polycomb and NCoR co-repressors are important developmental regulators that are recruited to chromatin by transcription factors subject to proteasomal degradation ^99–102^. Rather than being constantly degraded, Polycomb- or NCoR-recruiting transcription factors are turned over in response to ligand binding or posttranslational modifications, and these differences in degradation coincide with distinct biological roles: TLEs are short-range co-repressors that primarily act through transient chromatin deacetylation and compaction and do not impose a heritable epigenetic state ^27^. Accordingly, TLE-mediated repression is highly reversible and exquisitely sensitive to upstream signaling, enabling rapid switching between repression and activation during development. In contrast, Polycomb complexes establish self-reinforcing chromatin domains that confer developmental stability, often persisting independently of initiating transcription factors and resisting acute signaling inputs. Thus, TLE repression is optimized for plasticity and poised for transcriptional reprogramming, whereas Polycomb repression prioritizes robustness and epigenetic memory. We propose that it is systemic transcription factor degradation to release TLE co-repressors from chromatin that installs such dynamic regulation at the heart of human development and tissue homeostasis.

## Resource Availability

### Lead Contact

Further information and requests for reagents and resources should be directed to Michael Rapé.

### Materials Availability

All plasmids and cell lines generated in this work can be requested from the lead contact’s lab and will be freely shared. All antibodies, chemicals, and most cell lines used in this study are commercially available.

### Data Availability

Original proteomics data obtained by mass spectrometry were uploaded to the ProteomeXchange Consortium via the PRIDE repository with the dataset identifier PXD076746 and to be released upon publication. RNAseq and Cut&Run seq data have been uploaded to the GEO repository with GEO accessions GSE327629, GSE327631, GSE327633, and GSE327699 and to be released upon publication. Cryo-EM maps and atomic coordinates for the FBXL14/SKP1/mini-TLE1 complex have been deposited to the EMDB and wwPDB to be released upon publication, with the accession codes EMD-75774, EMD-75775, and PDB: 11KL. No original code was reported in this work. Additional information to reanalyze data reported in this work paper is available from the Lead Contact upon request.

## Acknowledgements

We thank all our lab members for help and enthusiastic discussions. We are grateful to the CRL Flow Cytometry Facility, the Vincent J. Proteomics/Mass Spectrometry Laboratory (NIH S10RR025622), and HTS Facility (NIH S10OD021828) at UCB. We thank the staff of Stanford Synchrotron Radiation Lightsource beamlines 14-1. This work was supported by Siebel Foundation postdoctoral award to PJ, an American Cancer Society award to SW, a Jane Coffin Childs postdoctoral award to HR. MR is an HHMI investigator.

## Author contributions

PJ performed most experiments. SW and ZY performed the structural analyses, and DM, HR, and AJ helped with experiments or data interpretation. All authors analyzed the data. MR supervised the work. PJ, SW and MR wrote the manuscript.

## Inclusion and Diversity

We support inclusive, diverse, and equitable conduct of research, and one or more of our authors identifies as an underrepresented minority in science.

## Supplementary Figures

**Figure S1:**
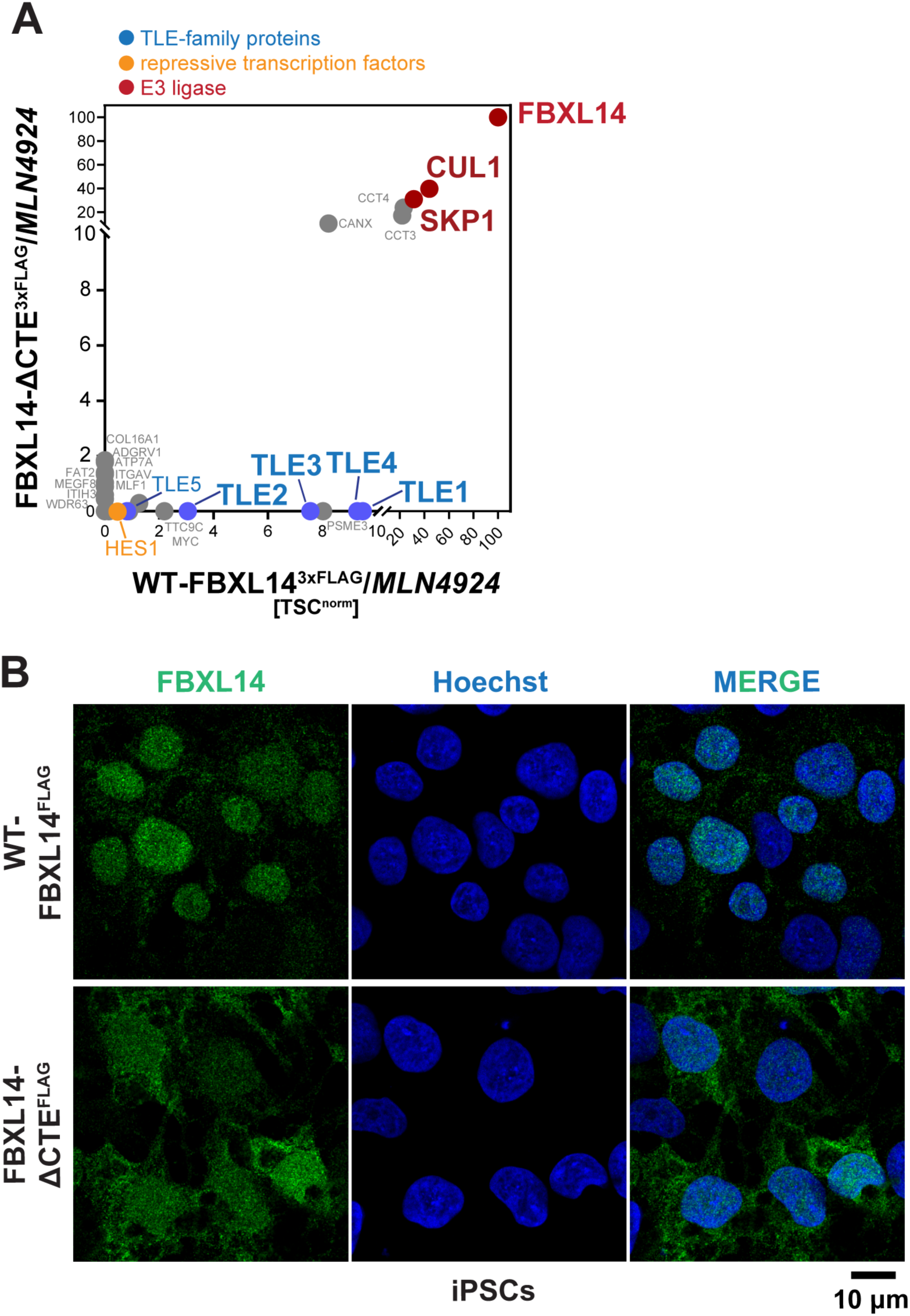
The SCF substrate adaptor FBXL14 engages all TLE co-repressors. **A.** FBXL14 binds all members of the TLE family of co-repressors dependent on its CTE. Full-length FBXL14^3xFLAG^ or FBXL14^ΔCTE/3xFLAG^ were affinity-purified from lysates of cells treated with the NEDD8-E1 inhibitor MLN4924 and binding partners were determined by mass spectrometry. **B.** FBXL14 localizes to the nucleus dependent on its interaction with TLEs. Localization of stably expressed FBXL14^3xFLAG^ or FBXL14^ΔCTE/3xFLAG^ was determined by αFLAG immunofluorescence microscopy (green: FBXL14; blue: Hoechst).

**Figure S2:**
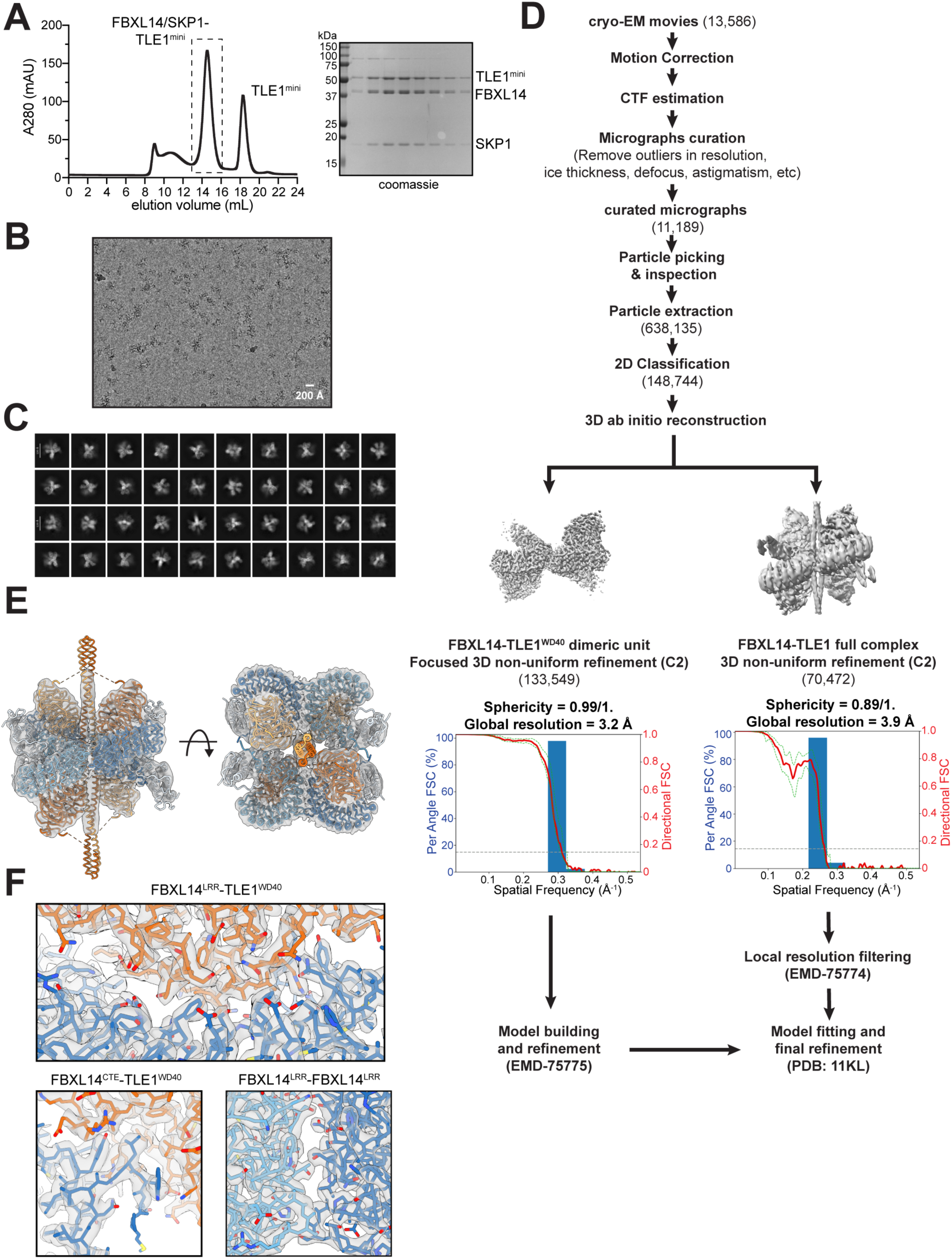
Cryo-EM structure of the mini-TLE1/FBXL14/SKP1 complex. **A.** Size exclusion chromatography trace and Coomassie-stained gel of FBXL14/SKP1/mini-TLE1 complex used for cryo-EM analysis. **B.** Motion-corrected and denoised representative micrograph. **C.** Representative 2D-class averages. **D.** Cryo-EM data processing particle flow and map resolution estimates. All processing steps were performed in CryoSparc. **E.** Full FBXL14/SKP1/mini-TLE1 model fit into the main cryo-EM map (EMD-75774) **F.** Model fit into the FBXL14/SKP1/TLE1^WD40^ high-resolution map obtained from a focused refinement of the region (EMD-75775).

**Figure S3:**
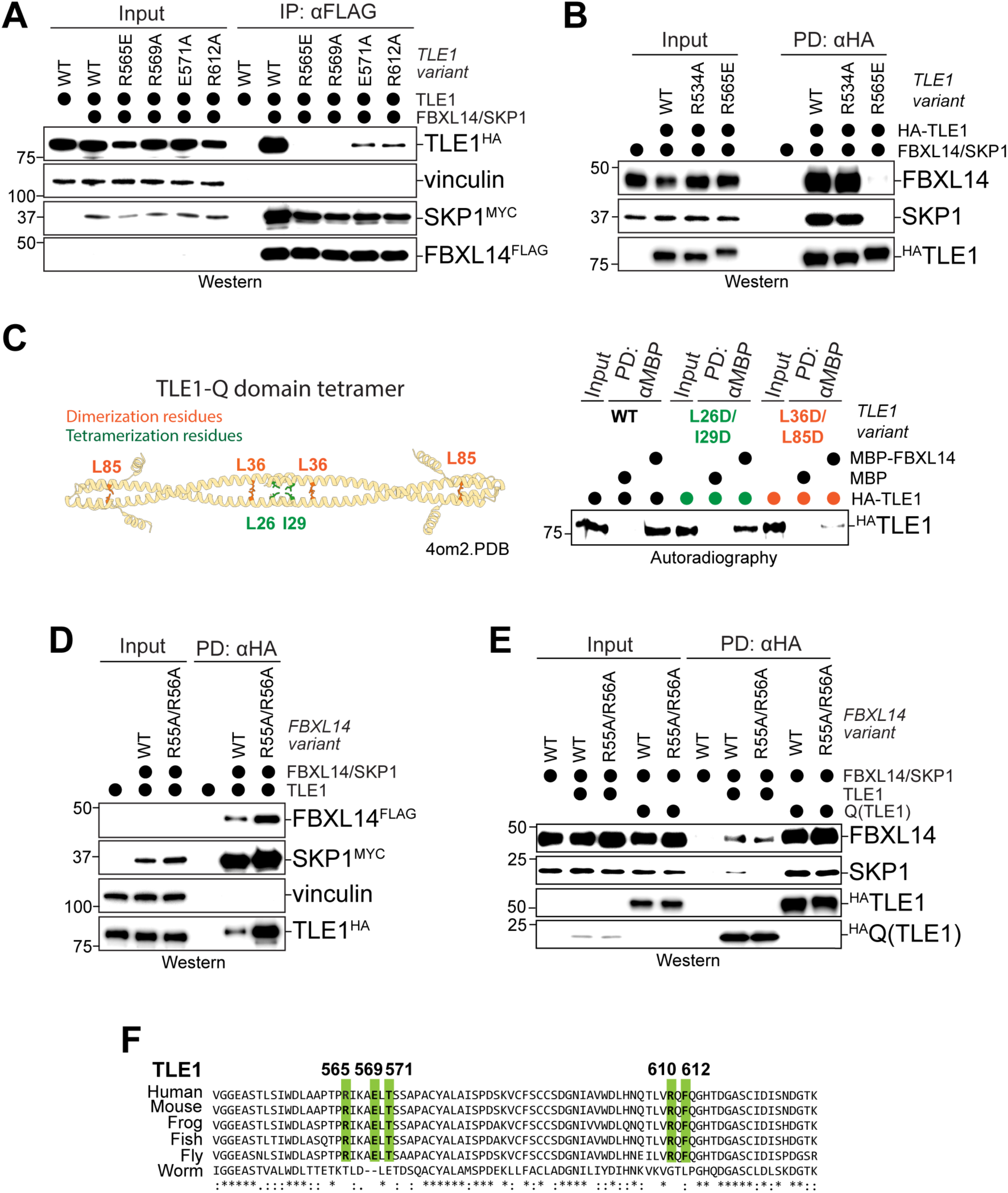
Validation of cryo-EM structure of the complex between FBXL14 and mini-TLE1. **A.** Mutations in TLE1 WD40-domain at the interface with the FBXL14 LRR-domain disrupt complex formation in cells. WT or mutant TLE1^3HA^ was co-expressed with FBXL14^3xFLAG^/SKP1^3xMYC^. FBXL14 was affinity-purified through αFLAG agarose, and co-precipitated TLE1 was detected by Western blotting. **B.** Mutations in the TLE1 WD40 disrupt binding to FBXL14 *in vitro*. WT- or mutant ^HA^TLE1 were immobilized and incubated with recombinant FBXL14/SKP1 complexes. Binding was visualized after gel electrophoresis by Western blotting. TLE1^R534A^ is defective in transcription factor binding, while TLE1^R565E^ has a mutation at the interface with FBXL14. **C.** Mutations in the TLE1 Q-domain disrupt binding to FBXL14 *in vitro*. MBP or ^MBP^FBXL14 were immobilized on beads and incubated with ^35^S-labeled WT- or mutant TLE1. Binding of TLE1 was monitored after gel electrophoresis by autoradiography. **D.** Arg-based salt bridges are not required for the interaction of FBXL14 with TLE1 in cells. WT or mutant FBXL14^3xFLAG^/SKP1^3xMYC^ was expressed in 293T cells, immobilized on αFLAG-agarose, and binding of endogenous TLE1 was monitored after gel electrophoresis by Western blotting. **E.** Arg-based salt bridges are not required for the interaction of FBXL14 with TLE1 *in vitro*. Recombinant ^HA^ TLE1 was immobilized on αHA-agarose and incubated with recombinant WT- or Arg-mutant (R55A/R56A) FBXL14. Binding of FBXL14 was monitored after gel electrophoresis by Western blotting. **F.** Conservation of residues that are required for the interaction between TLE1 and FBXL14 in organisms that express homologs of both proteins.

**Figure S4:**
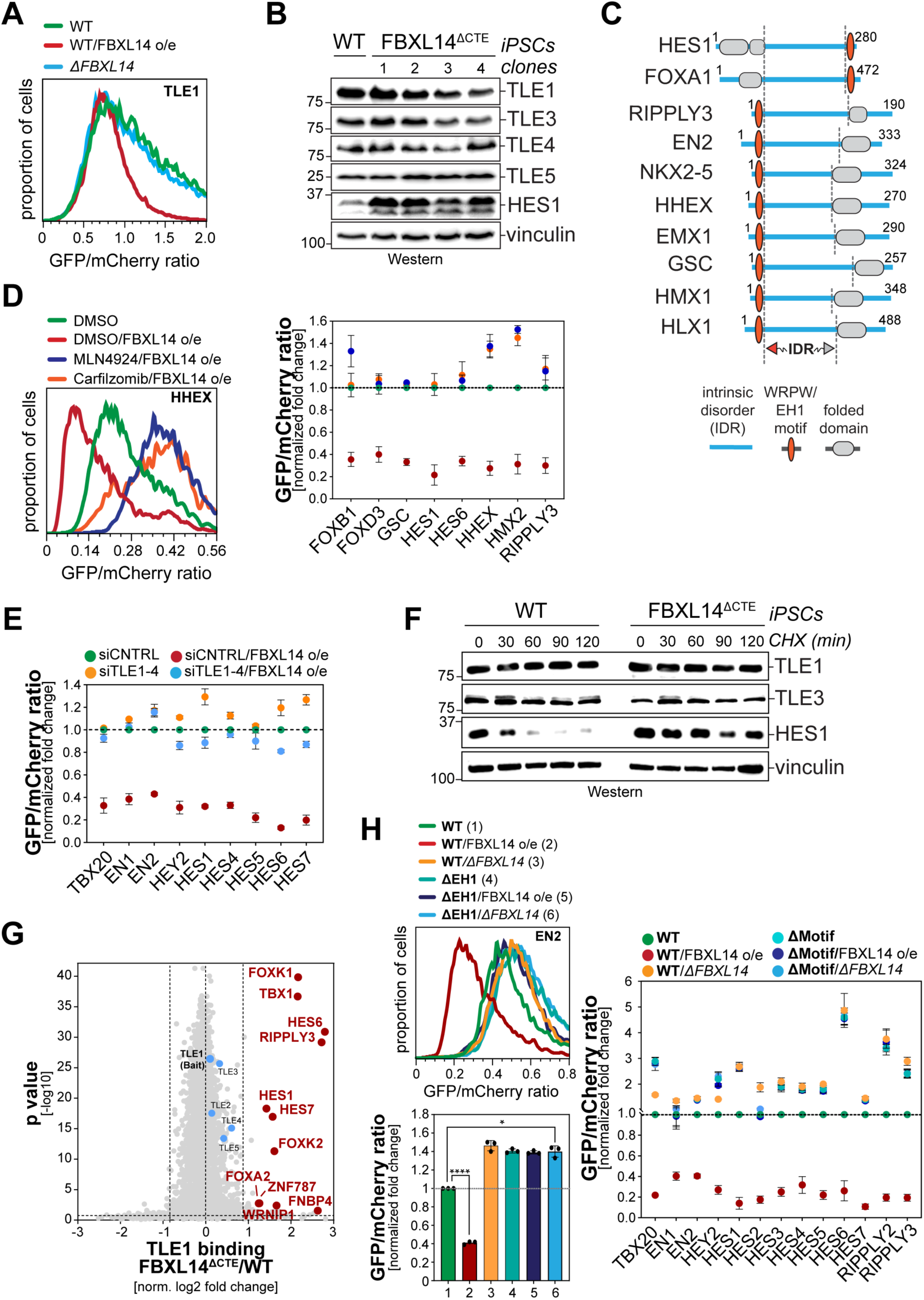
SCF^FBXL14^ targets TLE-bound repressive transcription factors for degradation. **A.** TLE1 is not degraded in an FBXL14-dependent manner in cycling 293T cells. Stability reporters for TLE1 were expressed in WT or *ΔFBXL14* 293T cells and analyzed by flow cytometry. **B.**TLE levels do not change in iPSCs deleted of the CTE in endogenous *FBXL14*. WT or independent clones of *FBXL14^ΔCTE^::*iPSCs were analyzed for endogenous TLE levels by Western blotting using specific antibodies. **C.** Degron motifs in SCF^FBXL14^-substrates are embedded in large regions of intrinsic disorder, shown here for examples of WRPW- or EH1-motif containing transcription factors. **D.** Transcription factor degradation upon FBXL14 overexpression is prevented by Cullin-RING E3 ligase inhibition (MLN4924, blue) or proteasome inhibition (carfilzomib, orange), as shown by flow cytometry. Quantification of three independent experiments for 8 transcription factors is shown on the right. **E.** Depletion of all full length TLEs expressed in human cells (TLE1-4) stabilizes several SCF^FBXL14^-substrates and protects them against degradation induced by FBXL14 overexpression. Stability of indicated transcription factors was measured using reporters in flow cytometry, and quantification of n=3 independent experiments is shown. **F.** Deletion of the CTE in the endogenous *FBXL14* loci, a mutation that prevents TLE binding, stabilizes endogenous HES1, as shown by cycloheximide chase and Western blotting. **G.** Deletion of the CTE in endogenous *FBXL14* of iPSCs traps several WRPW- and EH1-motif containing repressive transcription factors in complexes with endogenous TLE1, as determined by TLE1 affinity purification coupled to data-independent acquisition mass spectrometry. **H.** Deletion of WRPW-or EH1-motifs stabilizes multiple SCF^FBXL14^-substrates and protects them against degradation induced by FBXL14 over-expression. Stability was monitored using reporters by flow cytometry. In all experiments, data are represented from n=3 independent experiments +/- SD.

**Figure S5:**
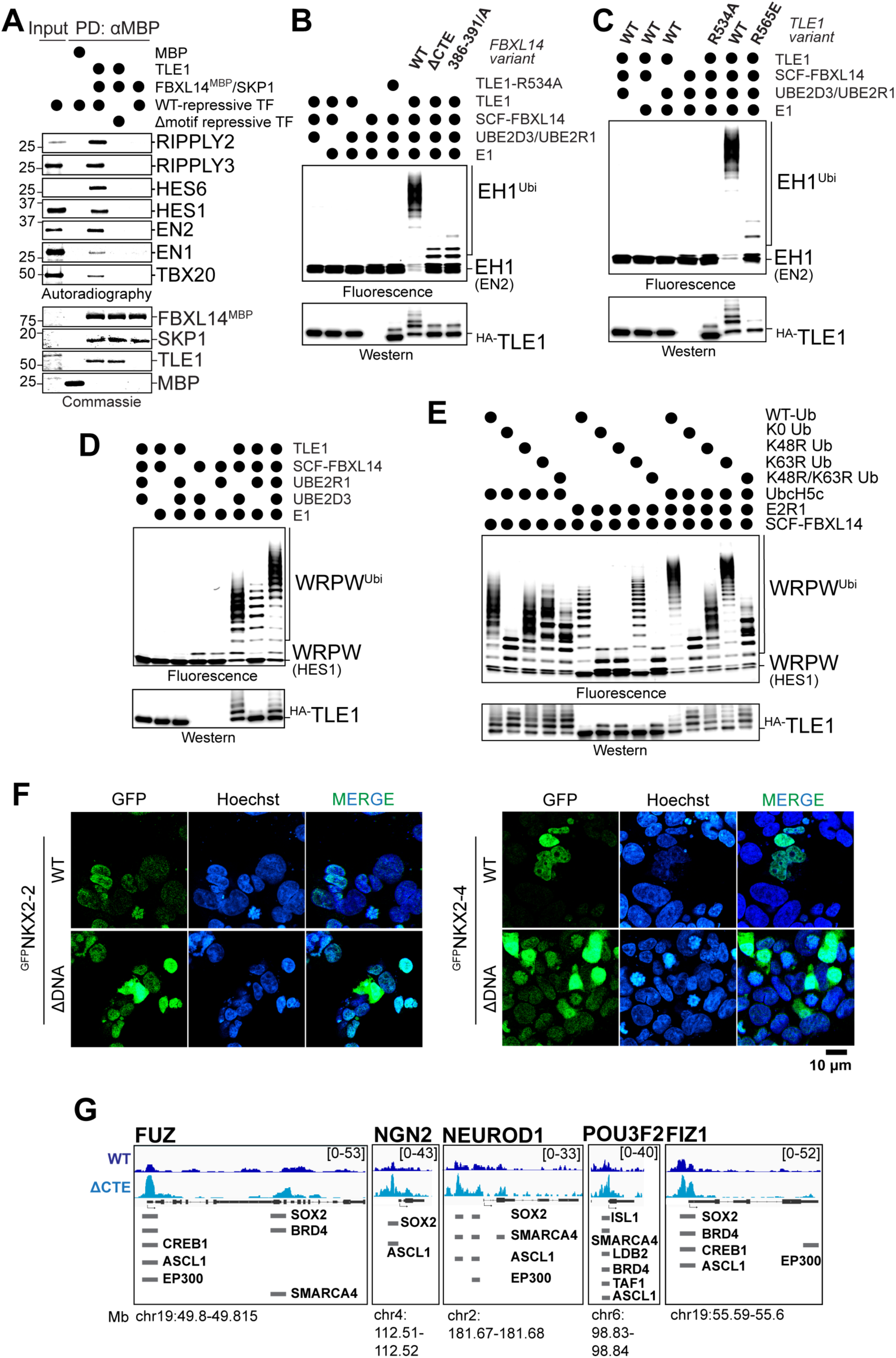
Repressive transcription factors are direct targets of SCF^FBXL14^. **A.** FBXL14 binds repressive transcription factors dependent on TLEs. ^MBP^FBXL14 was immobilized and incubated with recombinant TLE1 as indicated. ^35^S-labeled transcription factors (either WT or a variant with a deleted WRPW- or EH1-motif) were added and binding was detected by gel electrophoresis and autoradiography. **B.** SCF^FBXL14^ ubiquitylates EN2 dependent on its ability to bind TLEs. A TAMRA-labeled EH1 peptide derived from EN2 was incubated with recombinant SCF^FBXL14^, E1, E2s (UBE2D3, UBE2R1) and TLE1, as indicated. TLE1^R534A^, which is unable to recognize EH1 motifs, was added as control, and FBXL14 variants (ΔCTE; 386-391/A), which do not bind TLEs, were added as indicated. Ubiquitylation was monitored after gel electrophoresis by detected the TAMRA fluorescence. **C.** SCF^FBXL14^ ubiquitylates EN2 dependent on TLE1s ability to bind FBXL14. Ubiquitylation of a TAMRA-labeled EN2 peptides by SCF^FBXL14^ was analyzed as described above. WT-TLE1, TLE1^R534A^ (unable to bind transcription factors) and TLE1^R565E^ (unable to bind FBXL14) were added as indicated. **D.** SCF^FBXL14^ ubiquitylates substrate most efficiently when cooperating with both UBE2D3 and UBE2R1. Ubiquitylation of a TAMRA-labeled HES1 WRPW-motif peptide was performed as described above. UBE2D3 and UBE2R1 were added as indicated, and ubiquitylation was monitored after gel electrophoresis by fluorescence detection. **E.** SCF^FBXL14^ decorates substrates with K48-linked ubiquitin chains known to be recognized by proteasomes. Ubiquitylation of a TAMRA-labeled HES1 WRPW-motif peptide was performed in the presence of SCF^FBXL14^ and TLE1 as described above. Ubiquitin mutants (K0: all Lys residues mutated to Arg; K48R: Lys48 mutated to Arg; K63R: Lys63 mutated to Arg) were added as indicated. **F.** Transcription factor mutants lacking their DNA-binding domains localize to the cell nucleus. Localization of GFP-tagged transcription factors (WT or ΔDNA-binding domain variant) was analyzed in iPSCs using immunofluorescence microscopy. **G.** Genome browser tracks of TLE1 binding sites in either WT-iPSCs or *FBXL14^ΔCTE^*::iPSCs show overlap with binding sites for lineage specific activators.

**Figure S6:**
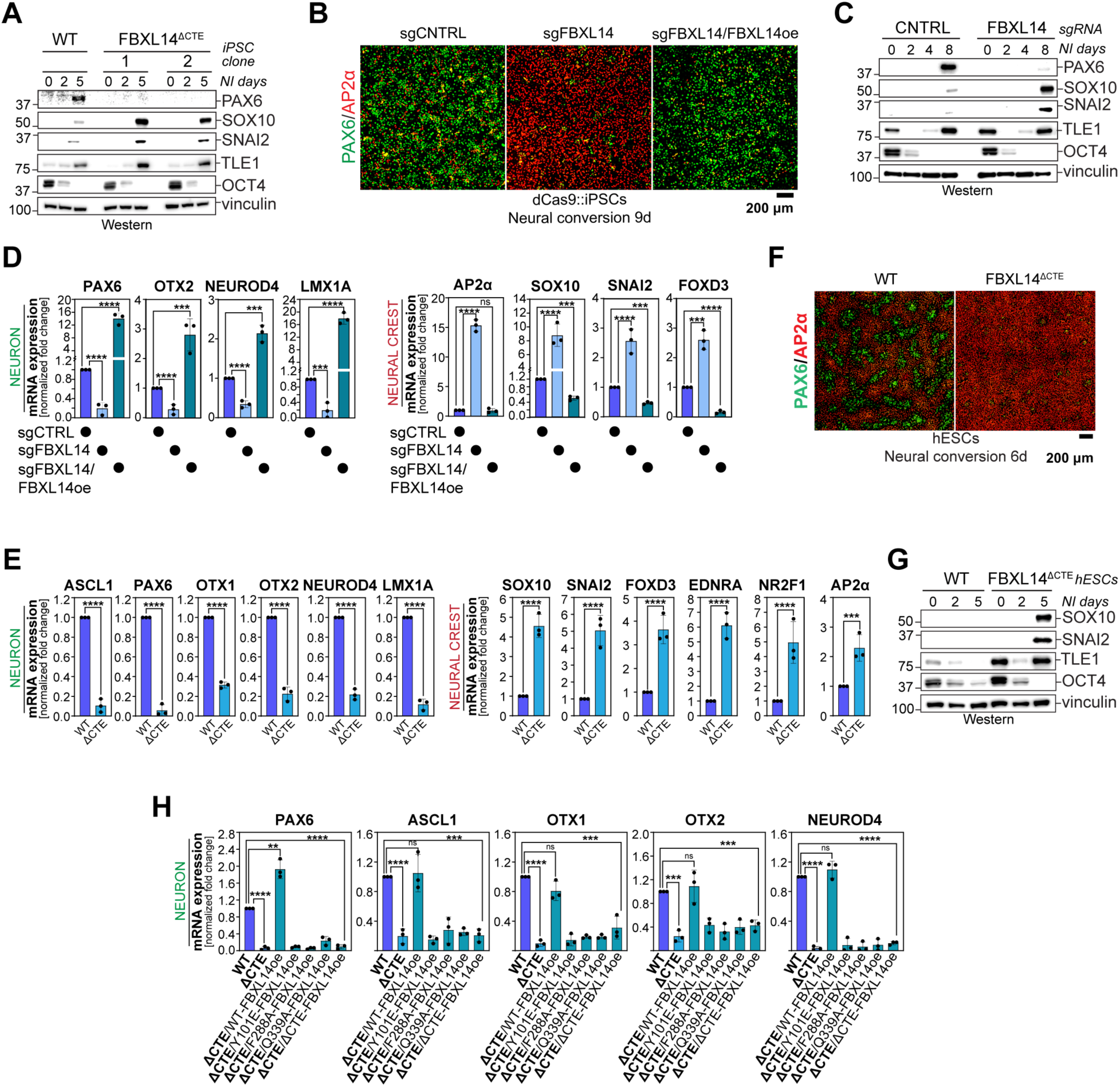
SCF^FBXL14^ controls cell fate. **A.** SCF^FBXL14^ is essential for neuronal differentiation of iPSCs. WT- or *FBXL14^ΔCTE^*::iPSCs were subjected to neural conversion and expression of cell fate markers (PAX6: neuron; SOX10, SNAI2: neural crest) was analyzed by Western blotting. **B.** dCAS9::iPSCs treated with either sgCNTRL or sgFBXL14 were subjected to neural conversion. As indicated, an sgRNA-resistant variant of WT-FBXL14 was expressed in sgFBXL14-treated cells. Expression of cell fate markers (PAX6: neuron; AP2α: neural crest) was analyzed by immunofluorescence microscopy. **C.** dCAS9::iPSCs treated with either sgCNTRL or sgFBXL14 were subjected to neural conversion, and expression of cell fate markers (PAX6: neuron; SOX10, SNAI2: neural crest) was analyzed by Western blotting. **D.** dCAS9::iPSCs treated with either sgCNTRL or sgFBXL14 were subjected to neural conversion. As indicated, an sgRNA-resistant variant of WT-FBXL14 was expressed in sgFBXL14-treated cells. Expression of cell fate markers was analyzed by qRT-PCR. **E.** WT or *FBXL14^ΔCTE^*::hESCs were subjected to neural conversion. Expression of cell fate markers was analyzed by qRT-PCR. **F.** WT or *FBXL14^ΔCTE^*::hESCs were subjected to neural conversion. Expression of cell fate markers (PAX6: neuron; AP2α: neural crest) was analyzed by immunofluorescence microscopy. **G.** WT or *FBXL14^ΔCTE^*::hESCs were subjected to neural conversion. Expression of cell fate markers was analyzed by Western blotting. **H.** WT- or *FBXL14^ΔCTE^*::iPSCs were subjected to neural conversion. As indicated, either WT-FBXL14 or various mutants unable to bind TLEs were re-expressed. and levels of cell fate markers was analyzed by qRT-PCR. In all experiments, data are represented from n=3 independent experiments +/- SD.

## Material and Methods

### Key resources table

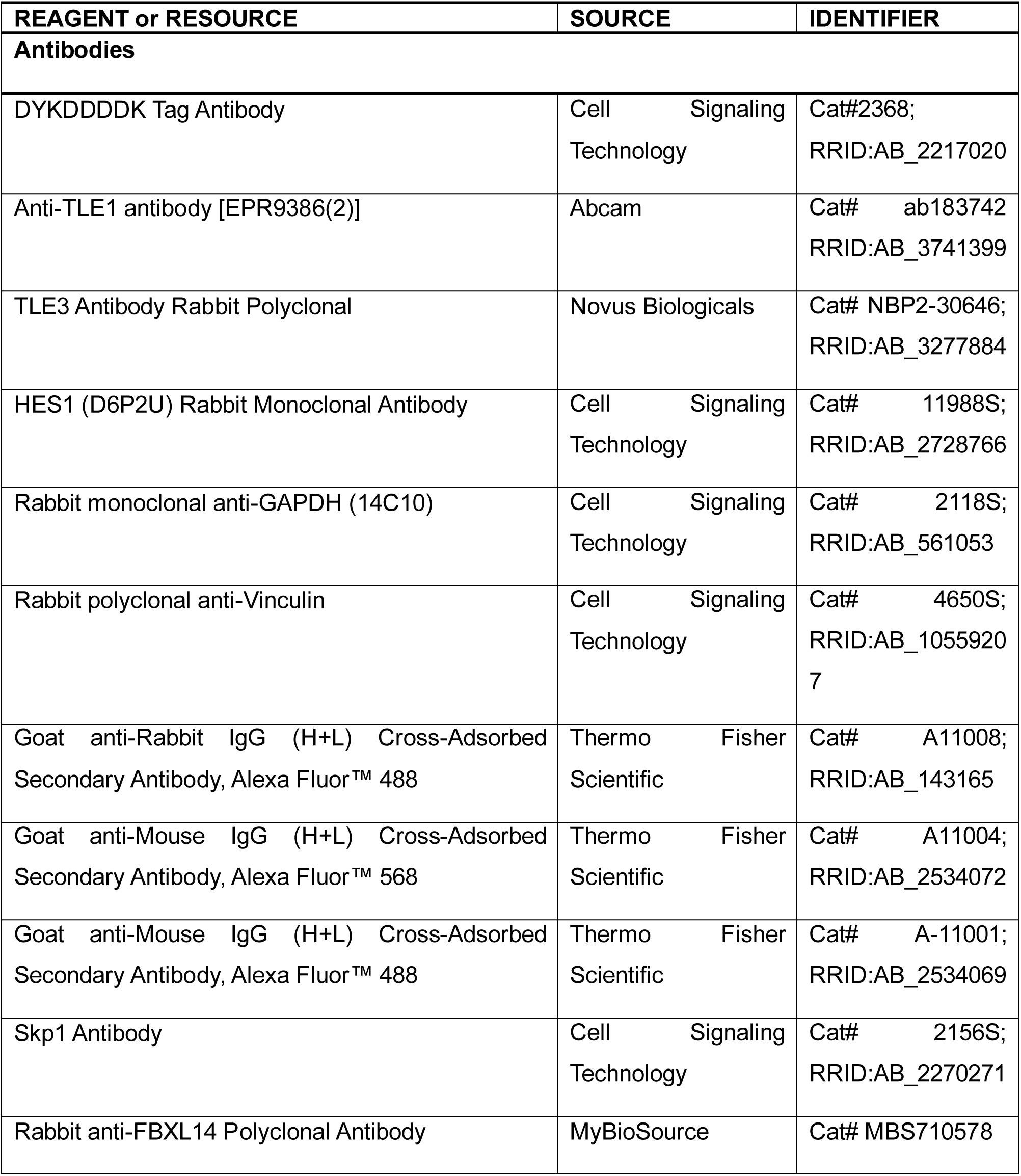
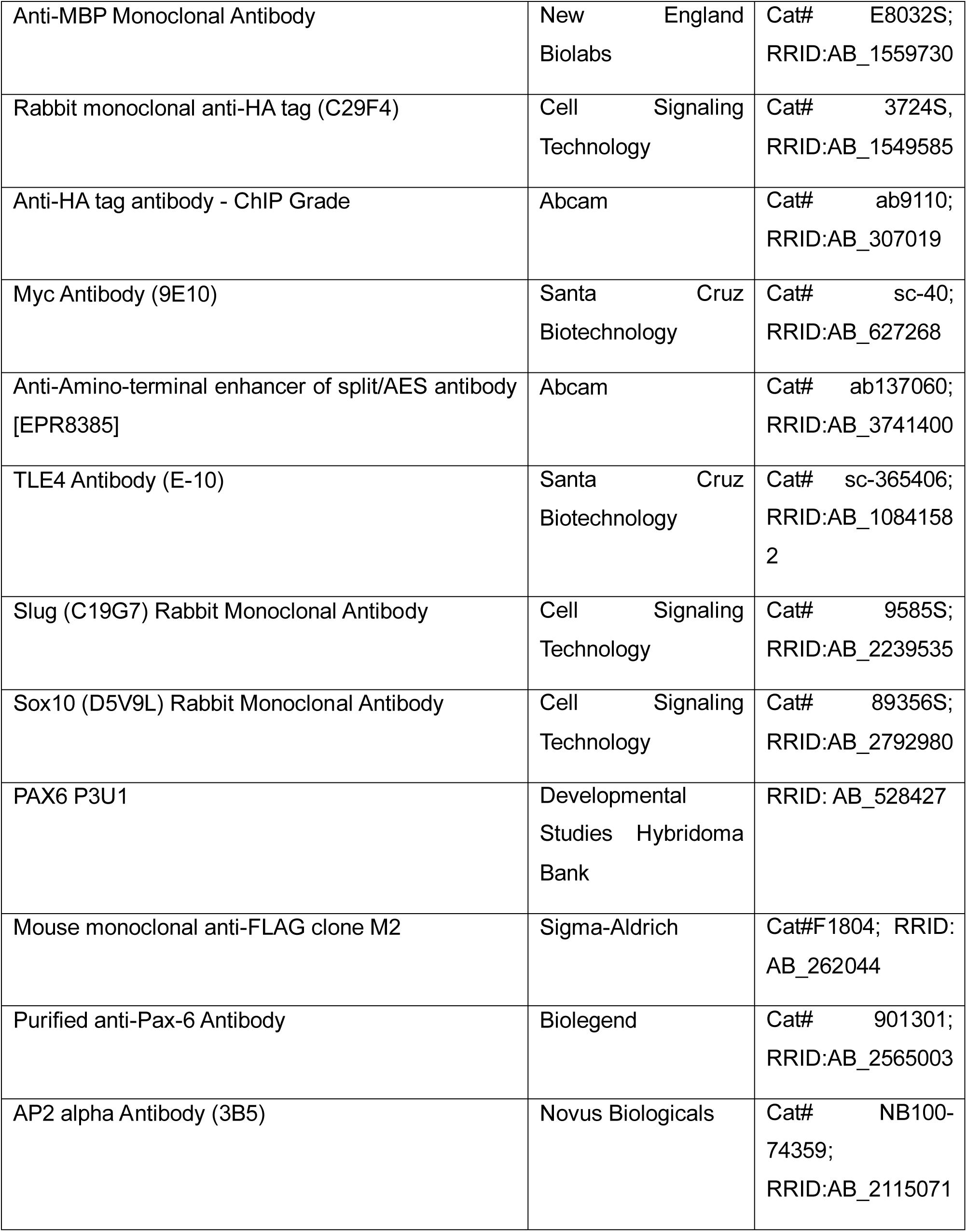
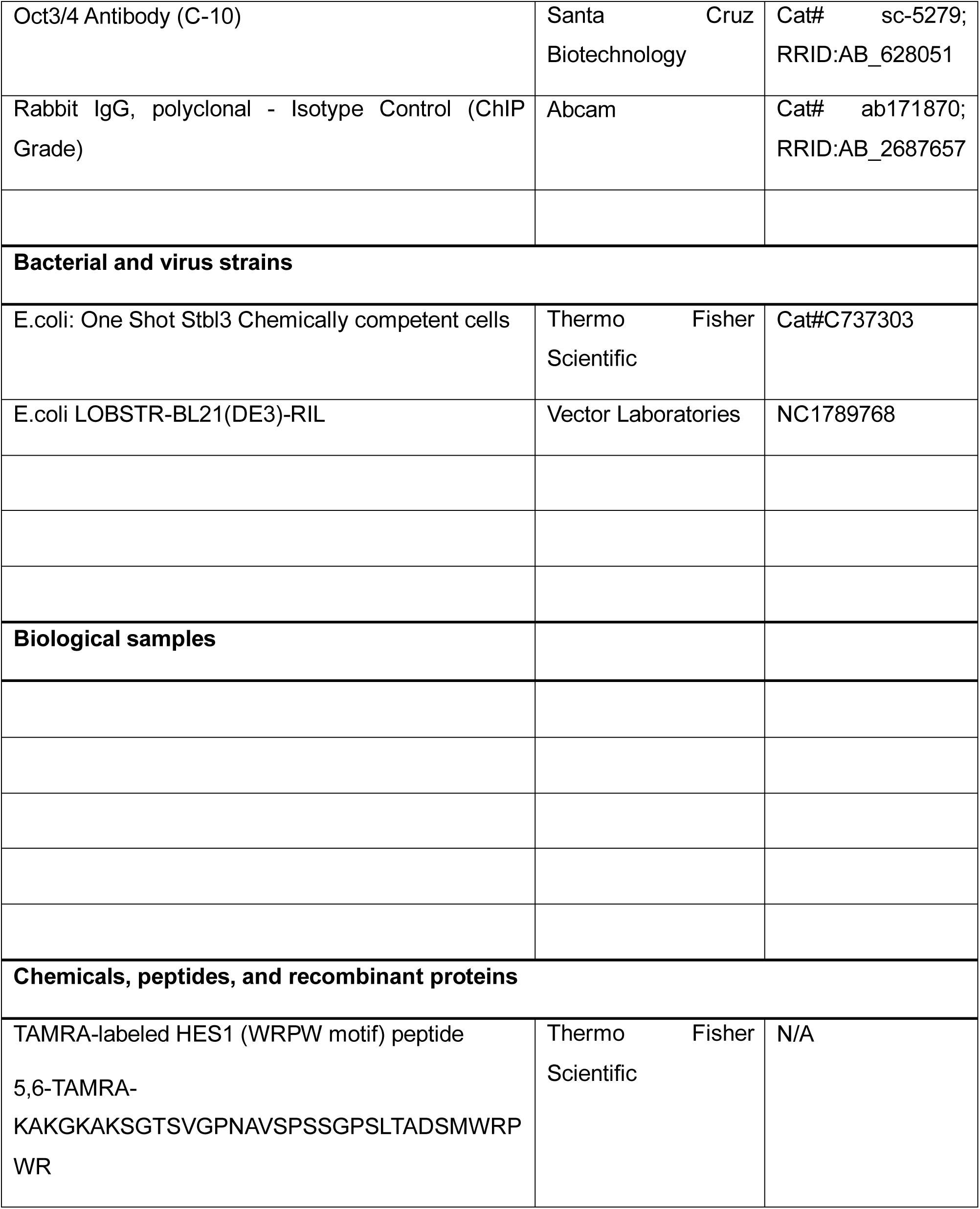
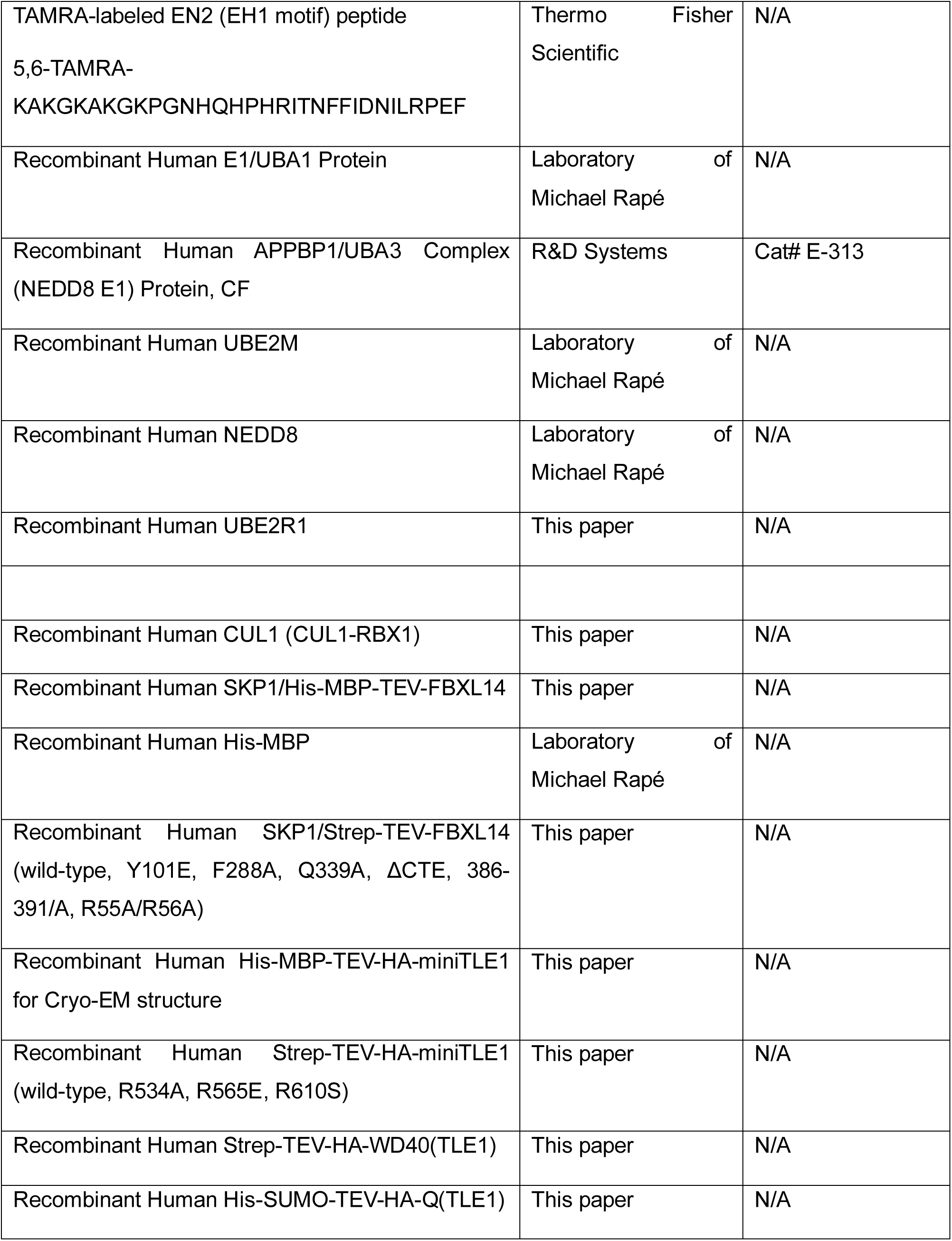
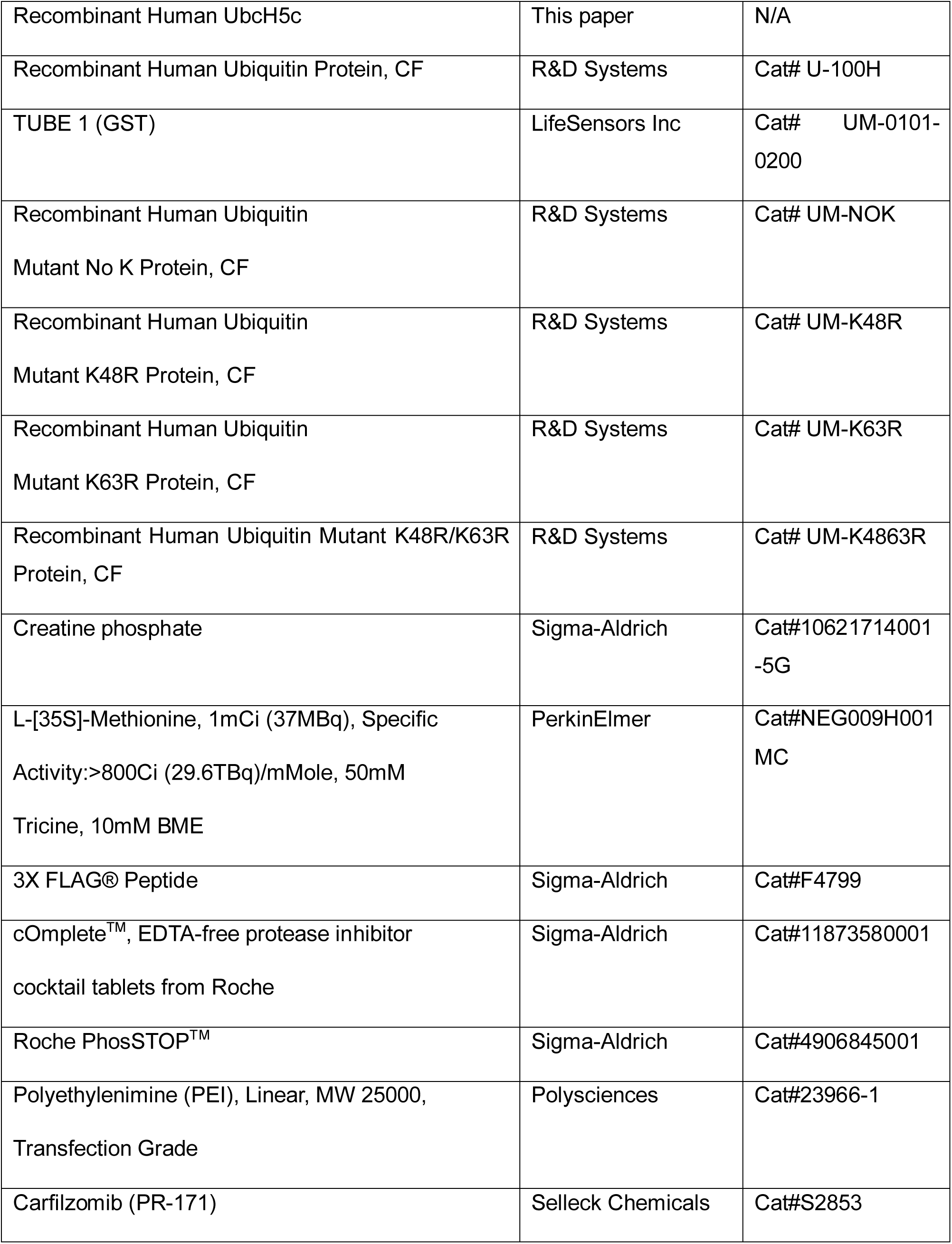
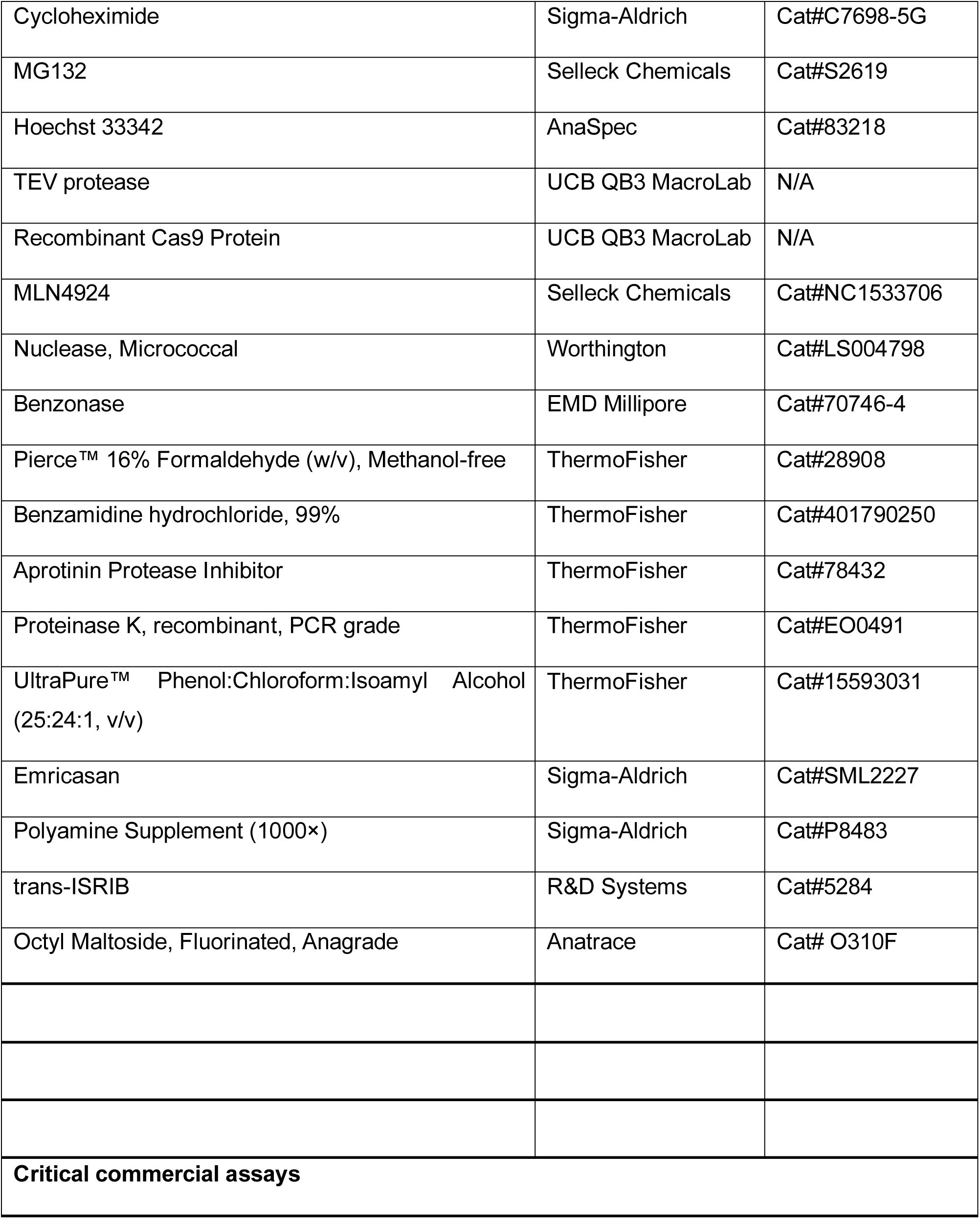
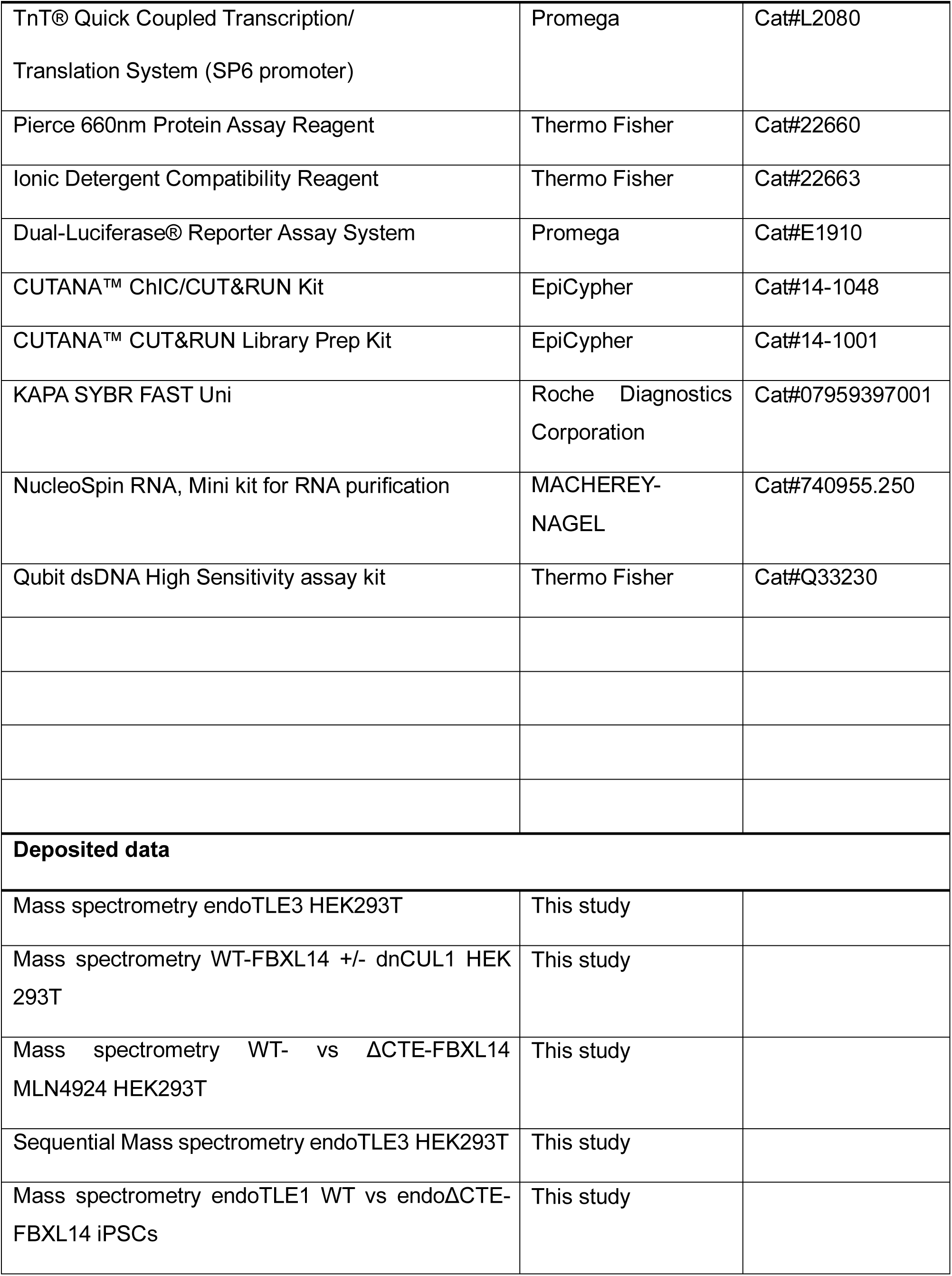
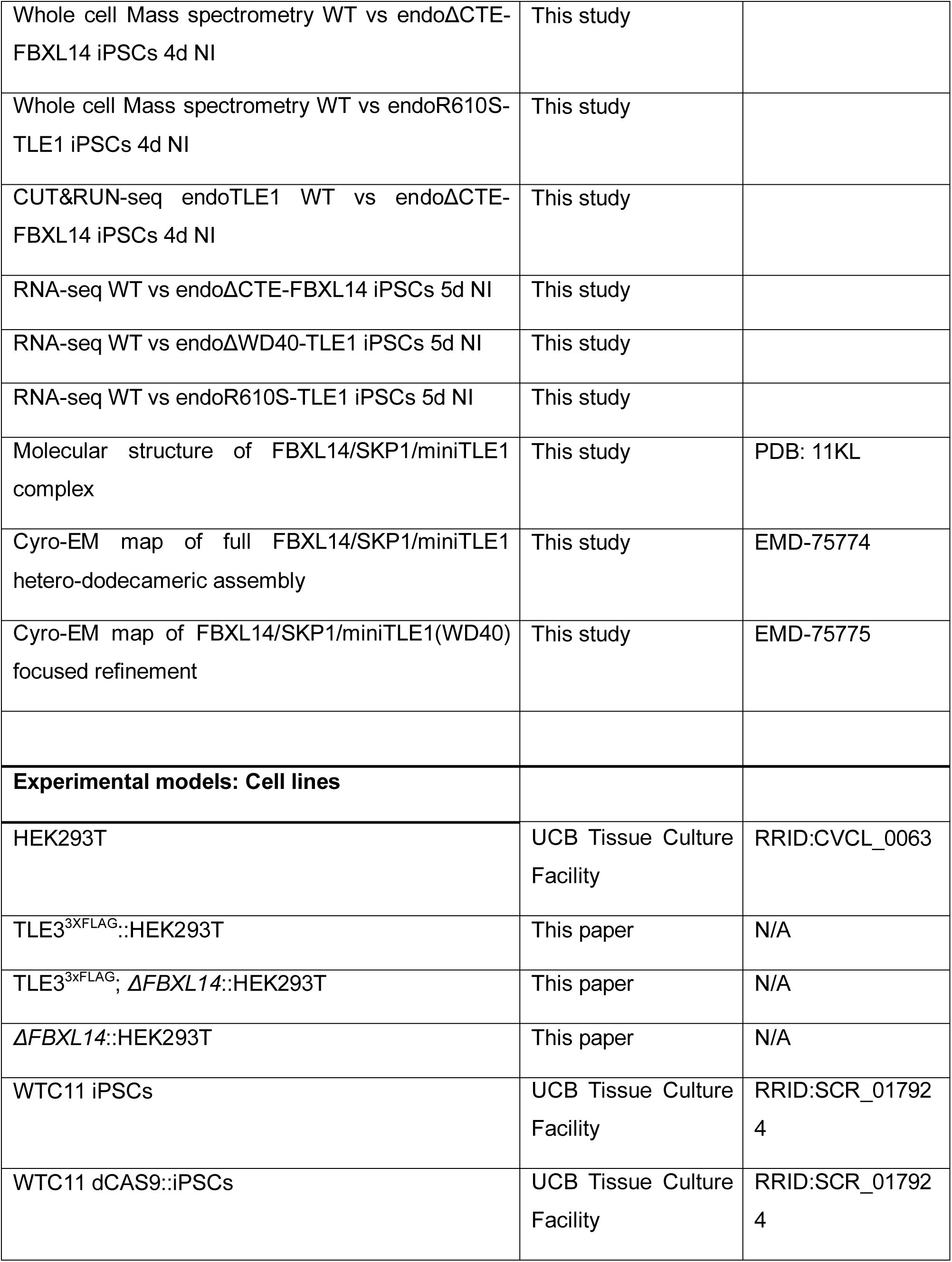
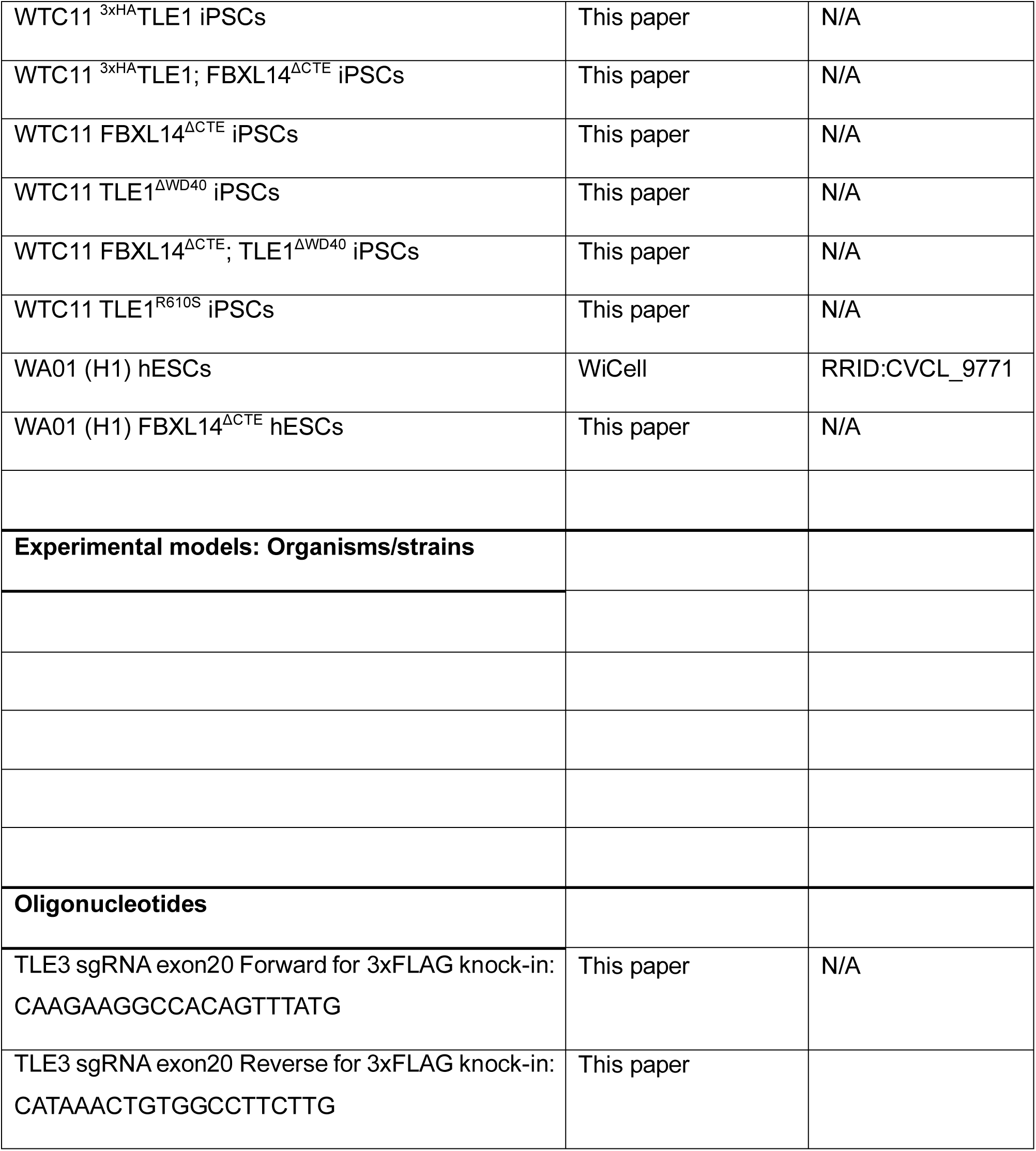
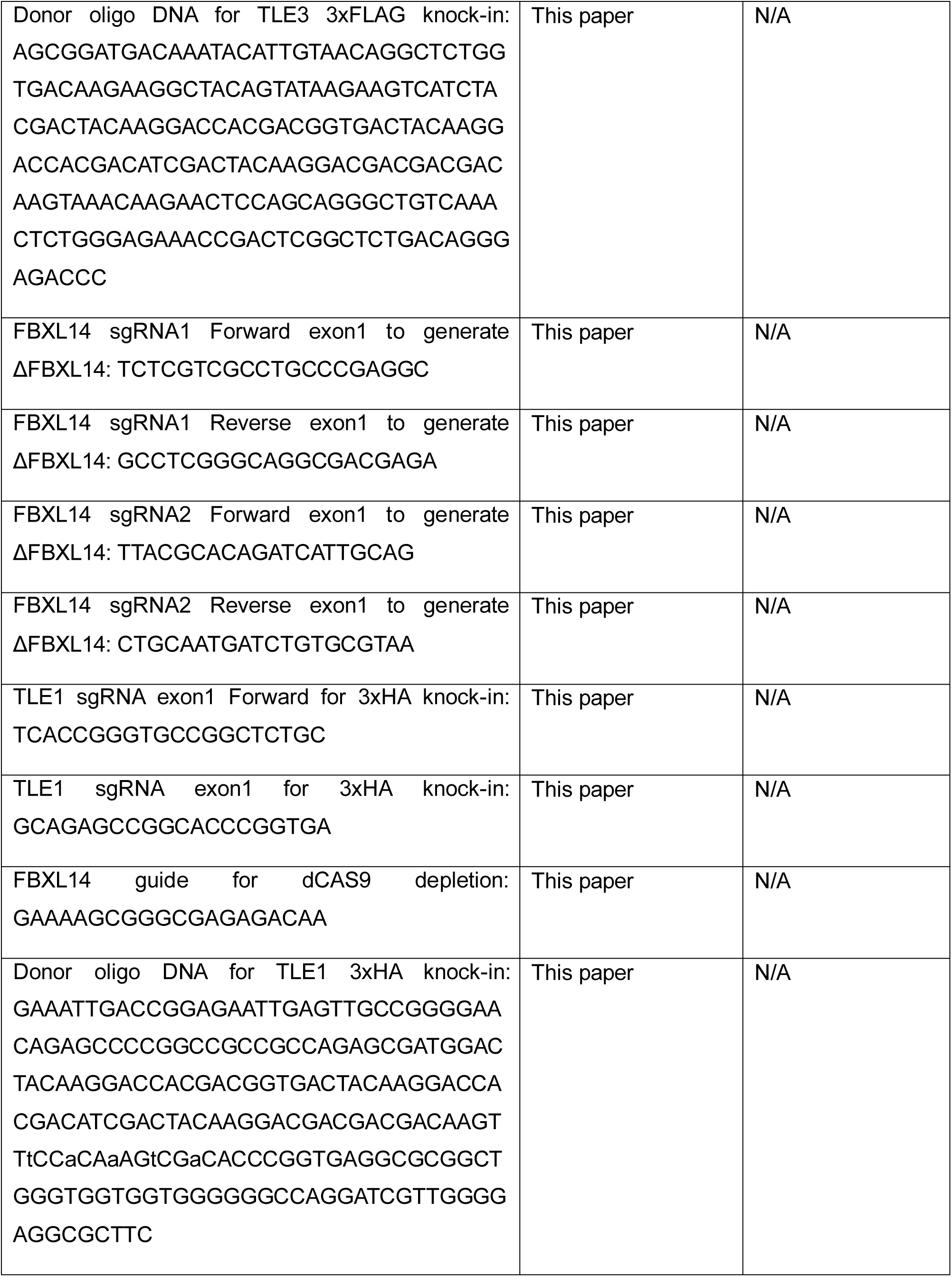
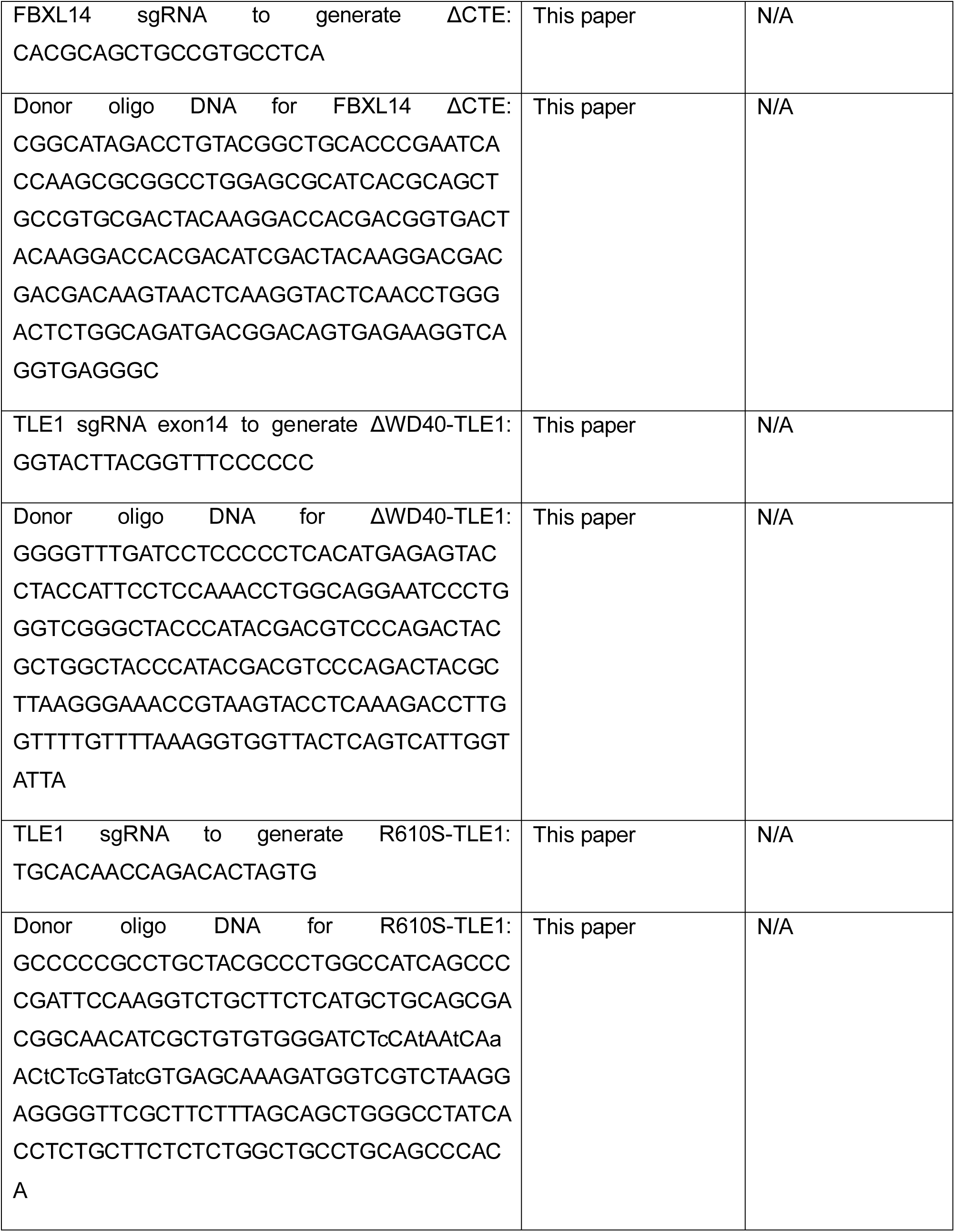
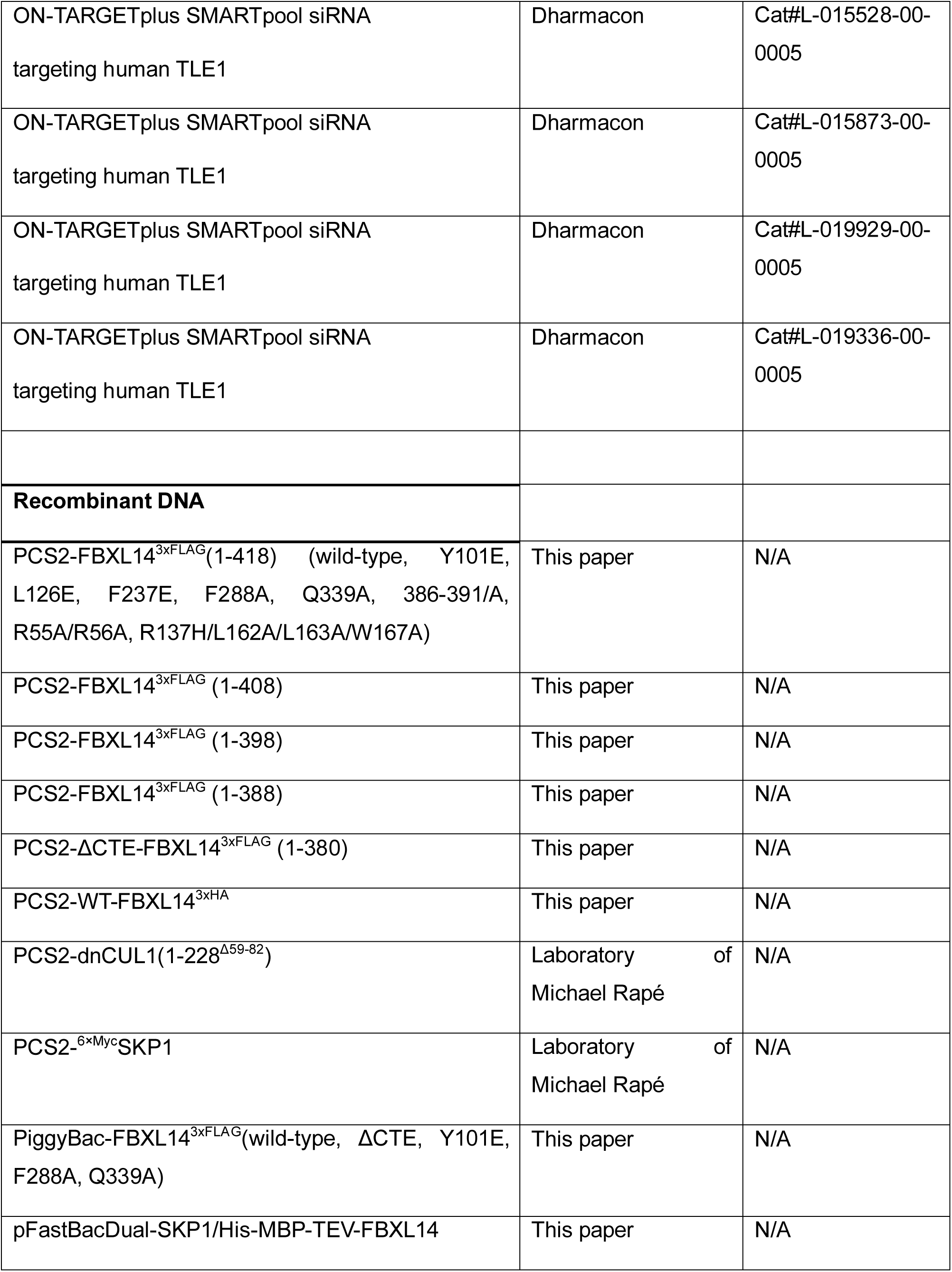
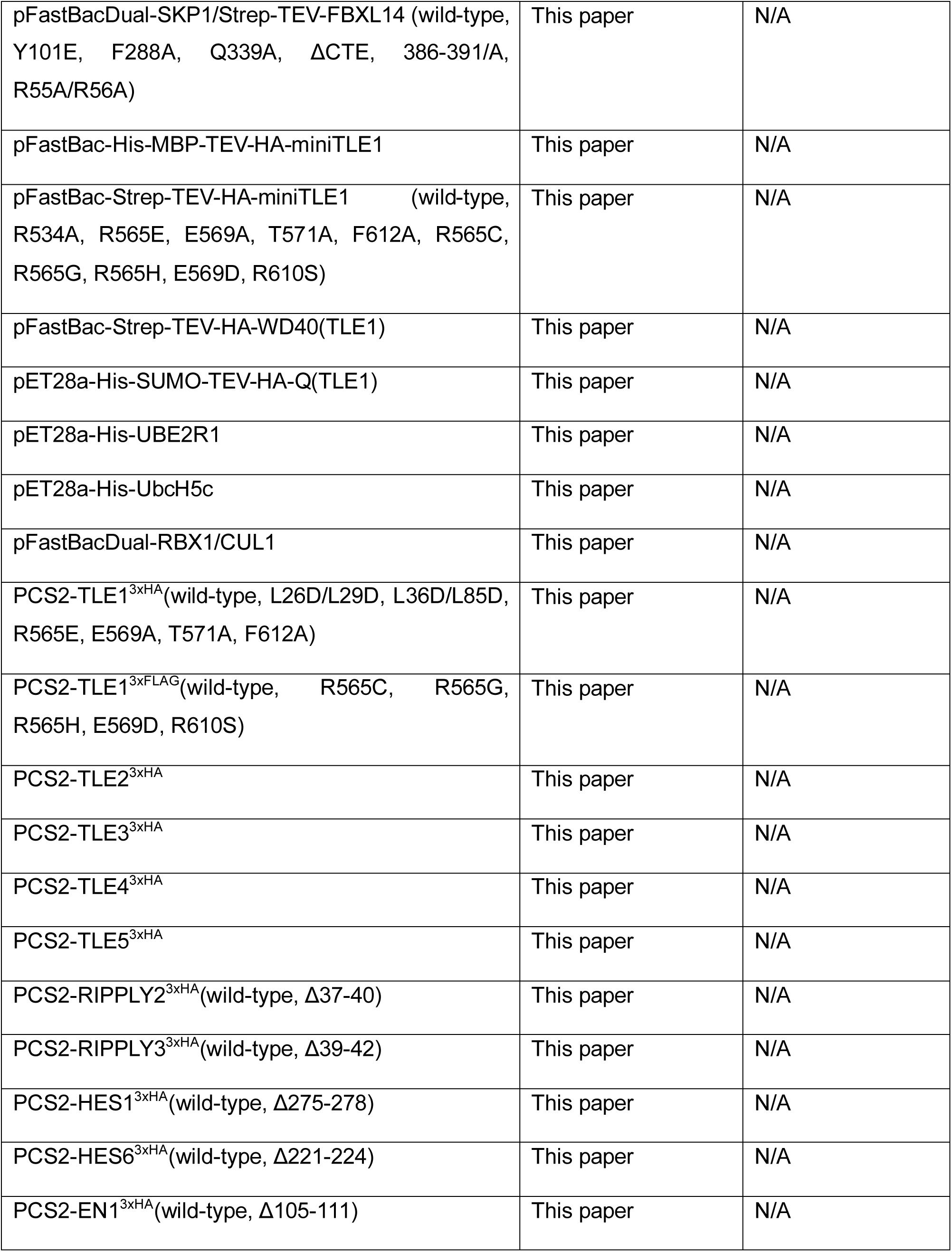
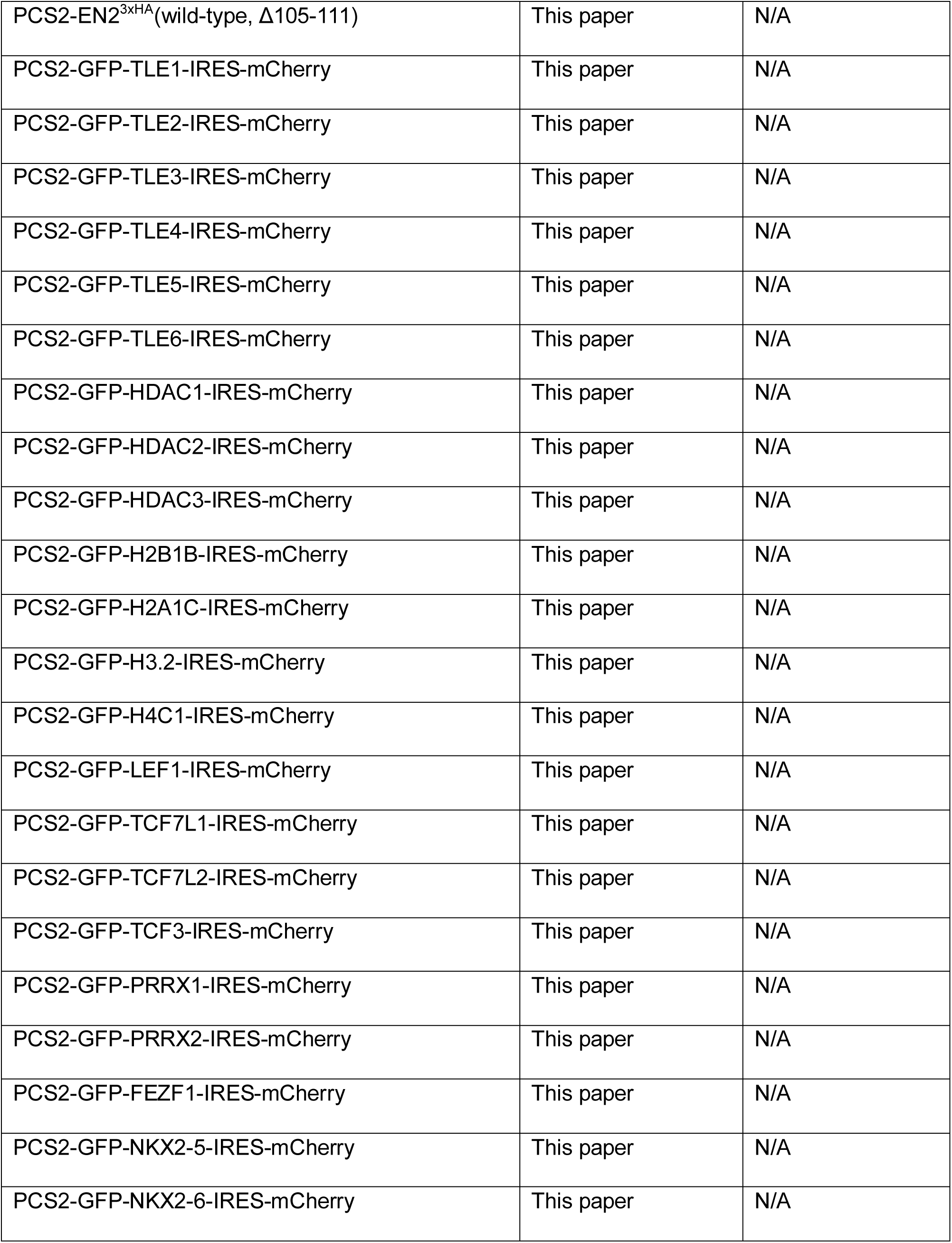
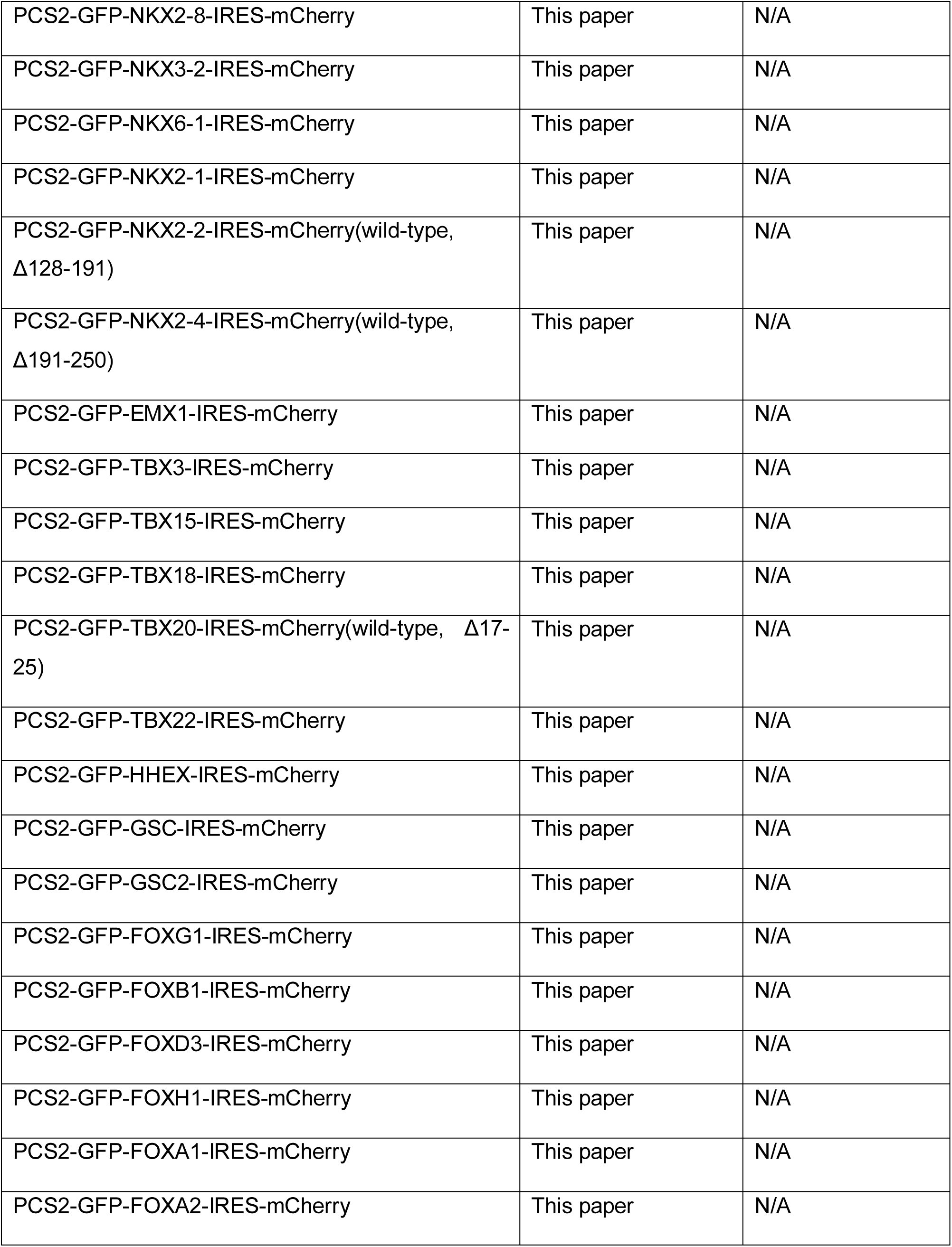
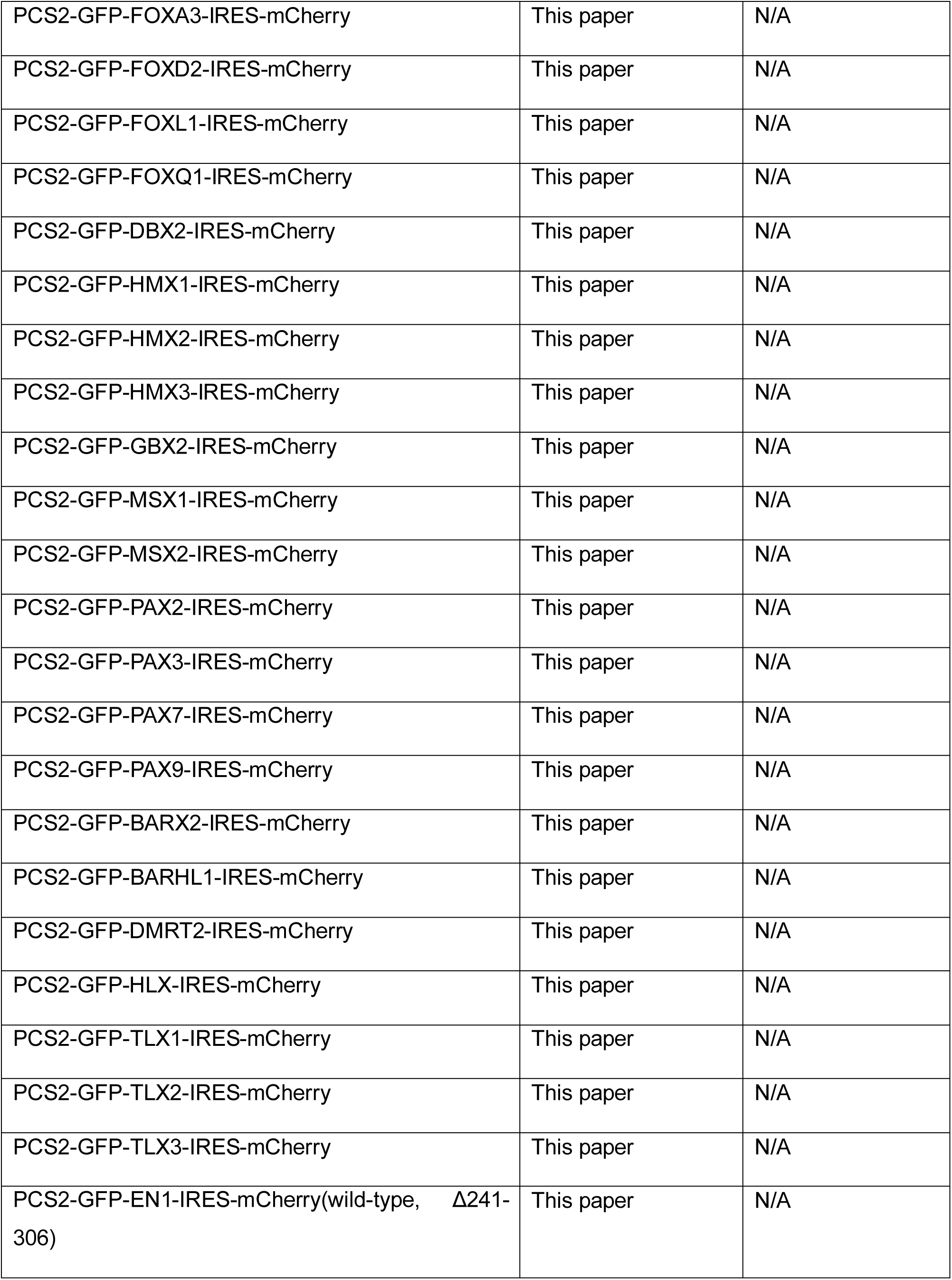
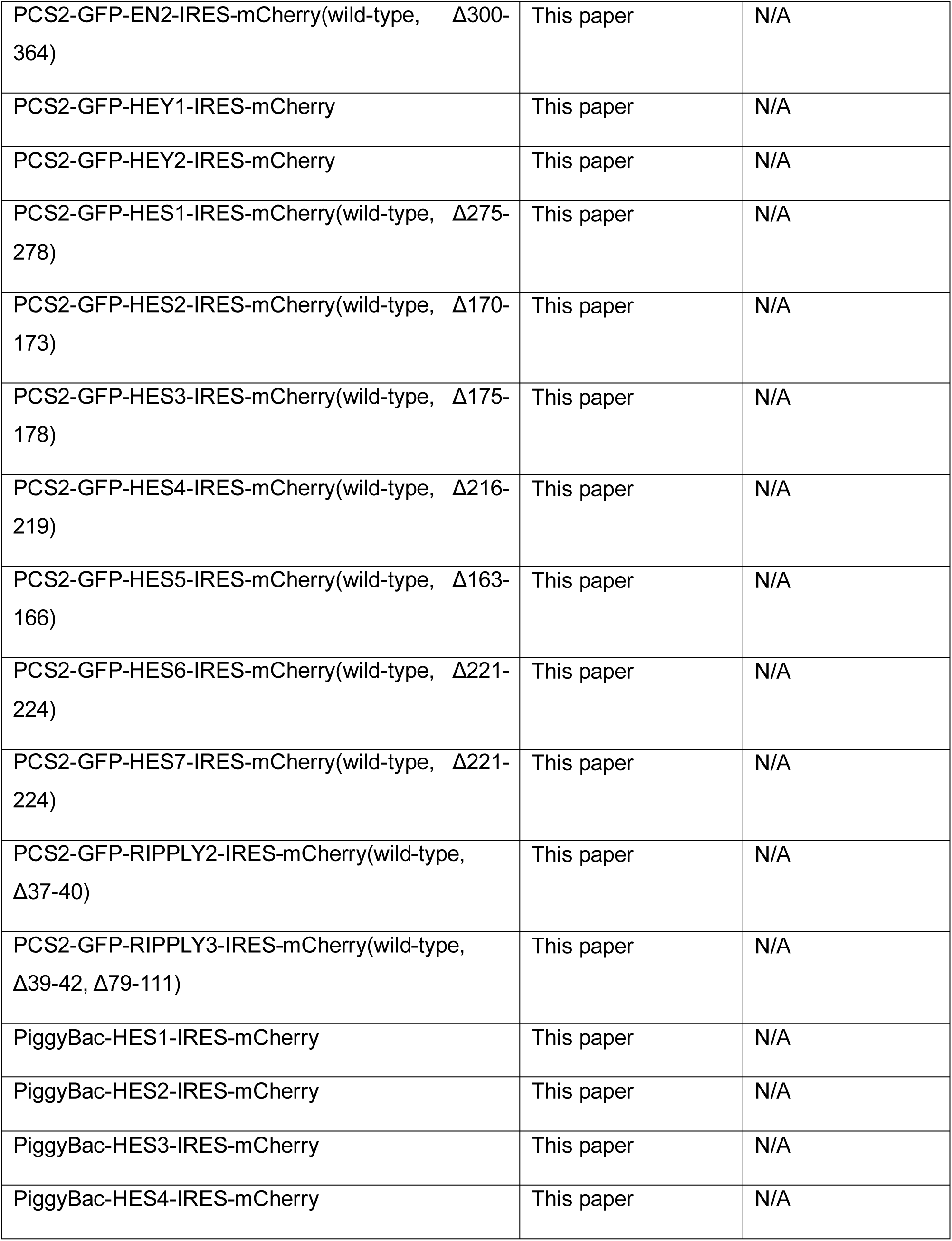
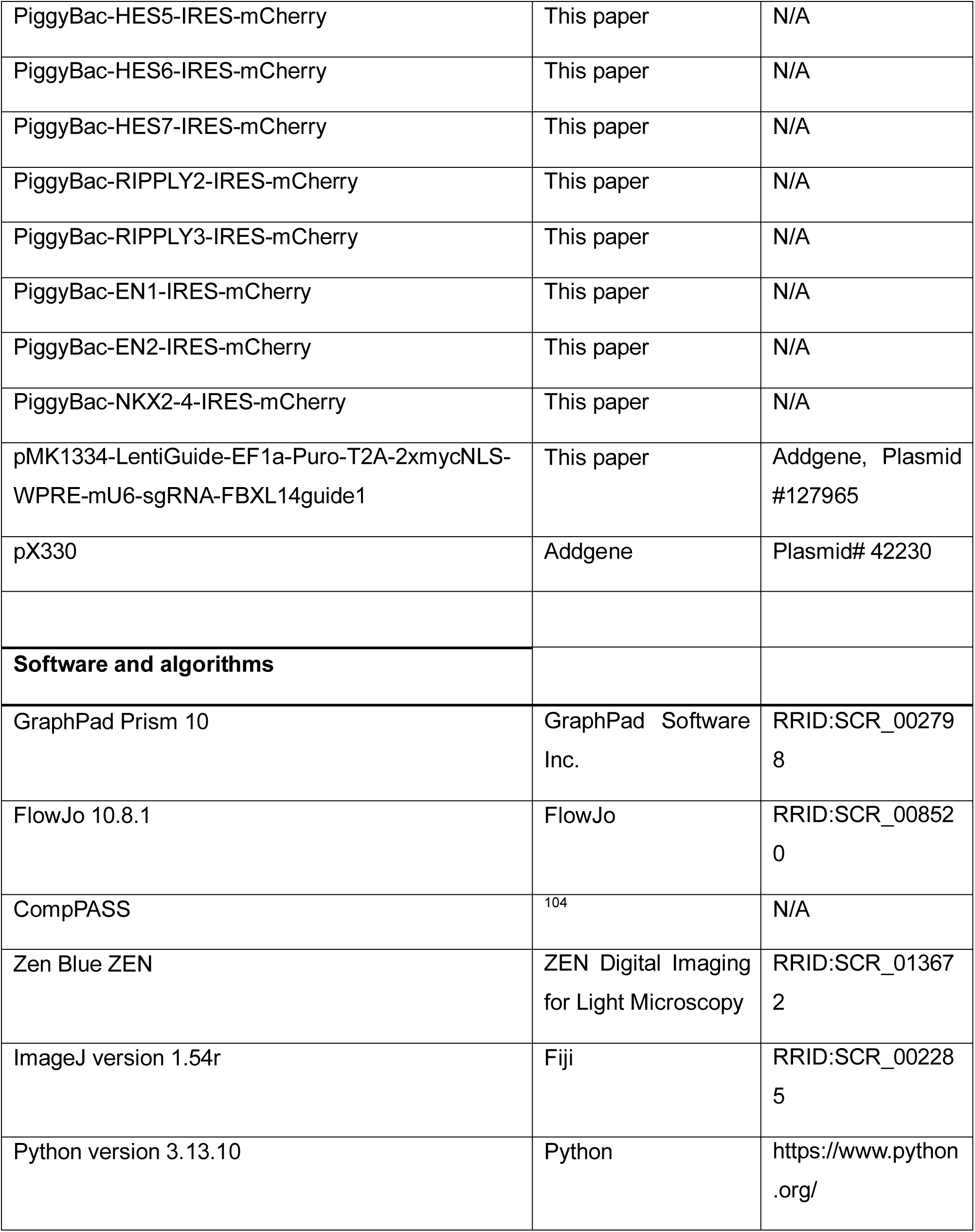
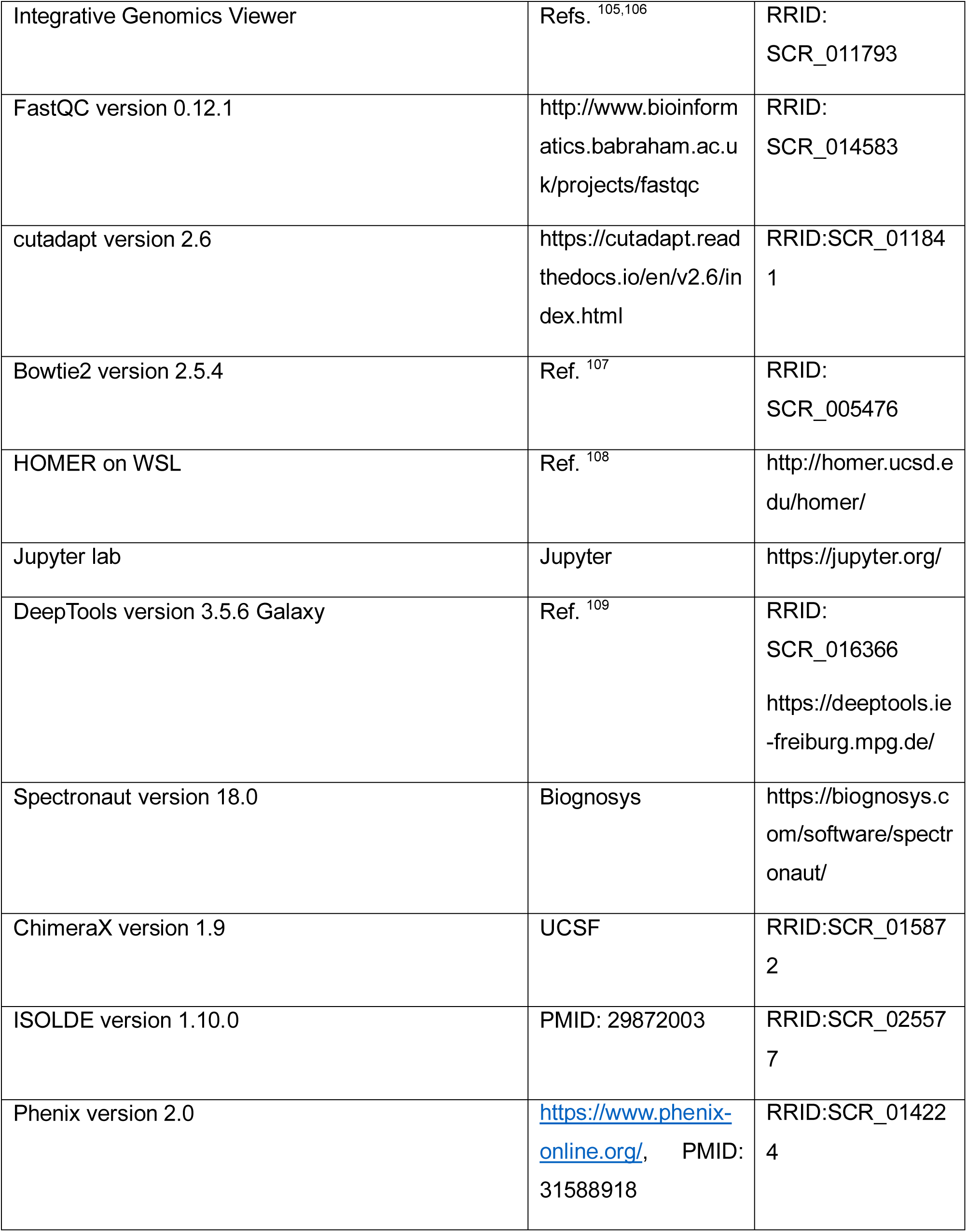
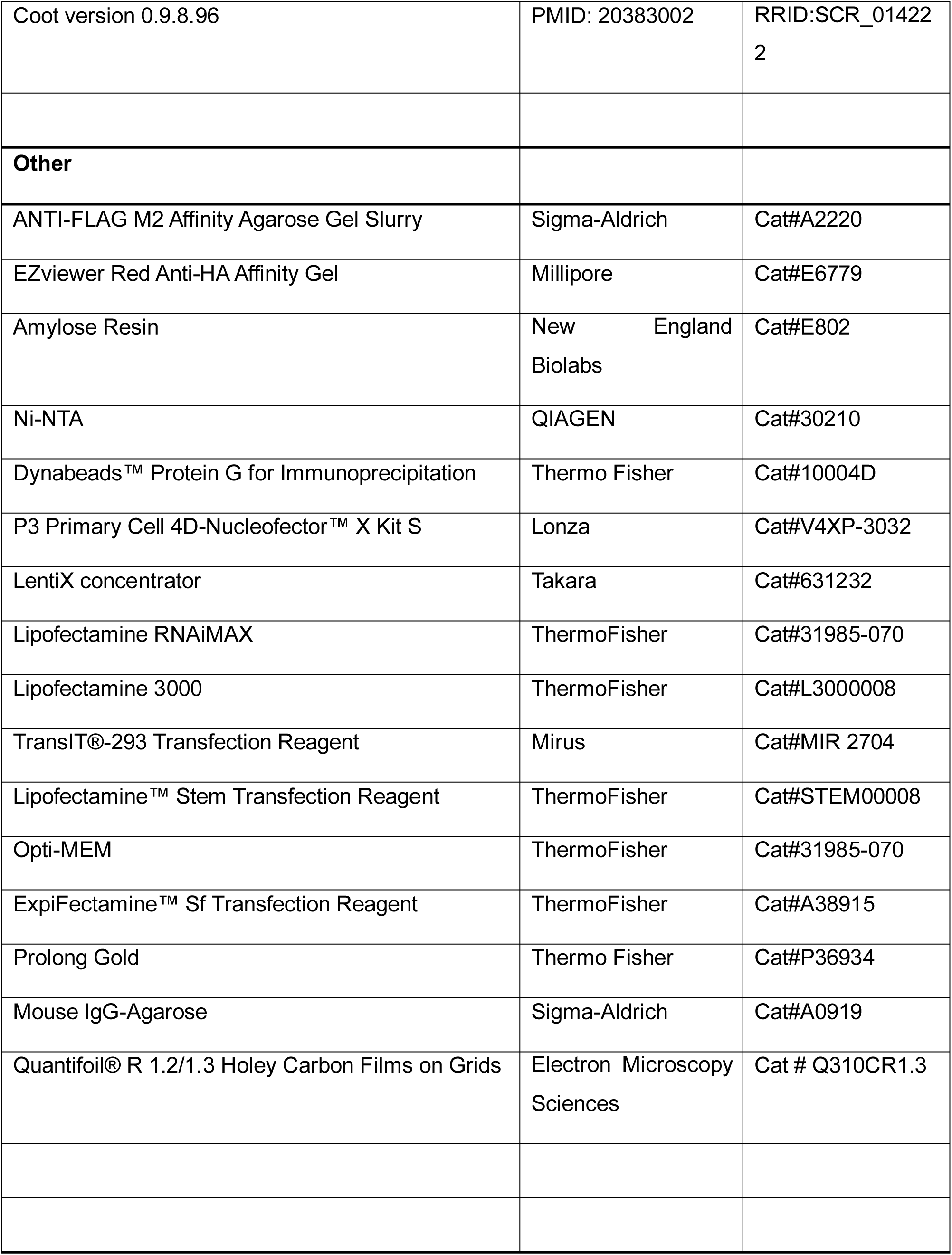
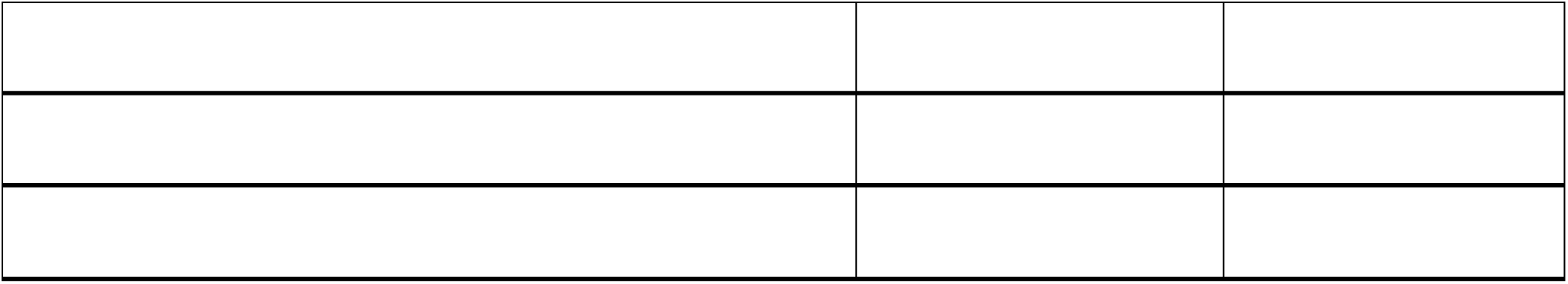

### EXPERIMENTAL MODEL AND STUDY PARTICIPANT DETAILS

Human embryonic kidney (HEK) 293T cells were maintained in DMEM + GlutaMAX (Gibco, 10566-016) supplemented with 10% fetal bovine serum (VWR, 89510-186). Plasmid transfections for immunoprecipitations were performed using polyethylenimine (PEI) at a 1:6 ratio of DNA (in mg) to PEI (in ml at a 1 mgml-1 stock concentration). Plasmid transfections for FACS-based stability assays were performed using using 6ul of TransIT®-293 Transfection Reagent (Mirus, MIR 2704) and 2ug total DNA plasmid per 6-well cell culture plate. siRNA transfections were performed with 20 nM of indicated siRNAs and 5ul of RNAiMAX transfection reagent (ThermoFisher, 13778150) per well in a 6-well cell culture plate. Lentiviruses were produced in HEK 293T cells by co-transfection of lentiviral and packaging plasmids using TransIT®-Lenti Transfection Reagent (Mirus, MIR 6603) according to the manufacturer’s protocol. Viruses were harvested 48 h post transfection, concentrated using the Lenti-X concentrator (Takara, 631232), aliquoted, and stored at 80°C for later use. HEK 293T cells were purchased directly from the Berkeley Cell Culture Facility (authenticated by short tandem repeat analysis).

Recombinant FBXL14/SKP1, miniTLE1, WD40-domain and CUL1-RBX1 for biochemical and structural studies were produced in Spodoptera frugiperda (Sf9) insect cells cultured at 27°C in SF4 Baculo Express Media (BioConcept, 9-00F38-K). Recombinant Q domain, UBE2R1 and UbcH5c were produced in E. coli BL21-CodonPlus(DE3)-RIL cells grown in LB broth media.

Human induced pluripotent stem cells (iPSCs) (WTC11) and human embryonic stem cells (WiCell, WA01/H1) were grown in mTeSR™Plus media (StemCell Technologies, 100-0276) on Matrigel-coated plates (Corning, 354277) with daily media change. Stem cells were passaged by Accutase (StemCell Technologies, 07920) for maintenance, PiggyBac vector transfections or lentiviral infections. Stem cells were dissociated with Accutase and 2x10^6^ cells were seeded on matrigel-coated 6-well plates and subjected to neural conversion using the STEMdiff™ Neural Differentiation medium with neural crest supplement (STEMCELL Technologies, 08610) which gives equal proportion of neural progenitors and neural crest cells. To generate stably expressing cell lines iPSCs were treated with Accutase and 5x10^5^ cells were seeded on a Matrigel-coated well of a 6-well plate with 2 ml mTeSR™Plus and 50 nM Chroman I (MedChem Express, HY-15392). The following day cells were transfected with 1875 ng PiggyBac vector and 625 ng Super PiggyBac Transposase Expression vector (System BioSciences, PB210PA-1) using Lipofectamine Stem Transfection reagent (Thermo Fisher Scientific, STEM00001). Cells were selected for 10 days with 20 µg ml^−1^ blasticidin (Thermo Fisher Scientific, A1113903). For lentiviral infection of guides dCas9-iPSCs were seeded at 3x10^5^ density on a Matrigel-coated well of a 6-well plate with 2 ml mTeSR™Plus and 50 nM Chroman I. The following day medium was removed, and cells were incubated overnight with 200 ul of concentrated virus and mTeSR™Plus up to 1 ml total volume. Cells were selected with 0.8 µg ml^−1^ puromycin (Millipore, P8833) for 5-7 days.

All cell lines were routinely tested for mycoplasma contamination using the Mycoplasma PCR Detection Kit (abmGood, G238).

### METHOD DETAILS

#### Antibodies

The following antibodies were used in this study: DYKDDDDK Tag Antibody (Cell Signaling Technology, 2368), Anti-TLE 1 antibody [EPR9386(2)] (Abcam, ab183742), TLE3 Antibody Rabbit Polyclonal (Novus Biologicals, NBP2-30646), HES1 (D6P2U) Rabbit Monoclonal Antibody (Cell Signaling Technology, 11988S), Rabbit monoclonal anti-GAPDH (14C10) (Cell Signaling Technology, 2118S), Rabbit polyclonal anti-Vinculin (Cell Signaling Technology, 4650S), Goat anti-Rabbit IgG (H+L) Cross-Adsorbed Secondary Antibody, Alexa Fluor™ 488 (Thermo Fisher Scientific, A11008), Goat anti-Mouse IgG (H+L) Cross-Adsorbed Secondary Antibody, Alexa Fluor™ 568 (Thermo Fisher Scientific, A11004), Skp1 Antibody (Cell Signaling Technology, 2156S), Rabbit anti-FBXL14 Polyclonal Antibody (MyBioSource, MBS710578), Anti-MBP Monoclonal Antibody, (New England Biolabs, E8032S), Rabbit monoclonal anti-HA tag (C29F4), (Cell Signaling Technology, 3724S), Myc Antibody (9E10) (Santa Cruz Biotechnology, sc-40), Anti-Amino-terminal enhancer of split/AES antibody [EPR8385] (Abcam, ab137060), TLE4 Antibody (E-10) (Santa Cruz Biotechnology, sc-365406), Slug (C19G7) Rabbit Monoclonal Antibody (Cell Signaling Technology, 9585S), Sox10 (D5V9L) Rabbit Monoclonal Antibody (Cell Signaling Technology, 89356S), PAX6 P3U1 (Developmental Studies Hybridoma Bank, RRID: AB_528427), Purified anti-Pax-6 Antibody (Biolegend, 901301), Oct3/4 Antibody (C-10) (Santa Cruz Biotechnology, sc-5279).

#### Cloning

All genes were cloned from cDNA prepared from HEK293T or stem cells. The list of all constructs used in this study are provided in the Key resources table. Most cloning was performed using Gibson assembly using HIFI DNA Assembly master mix (NEB, E2621L).

#### Flow cytometry

HEK293T cells were seeded at 400,000 cells per well in 6-well plates. The next day, 100 ng of GFP-degron-IRES-mCherry reporters, 500 ng of FBXL14 constructs, 500 ng dnCUL1, 170 ng of SKP1 and empty vector up to maximum 2 ug total were transfected using 6 ul TransIT®-293 Transfection Reagent (Mirus, MIR 2704). After 24 h, cells were harvested for flow cytometry. In experiments using siRNA depletion, siRNAs were reverse transfected (at the time of seeding cells) with 20 nM of indicated siRNAs and 5ul of RNAiMAX transfection reagent (ThermoFisher, 13778150) per well in a 6-well cell culture plate. The next day plasmids were transfected into each well using 5 ul Lipofectamine 3000 (ThermoFisher, L3000008). Cells were treated with the following reagents at indicated times before harvesting: 50 nM Carfilzomib (Selleck, PR-171) overnight or 500 nM MLN-4924 (Selleck, S7109) overnight. 24 h post-transfection, cells were trypsinized and resuspended in DMEM with 10% FBS and analyzed on either BD Bioscience LSR Fortessa or LSR Fortessa X20 and FlowJo. To test substrate stability in stem cells either WT, FBXL14^ΔCTE^ or TLE1^R610S^ iPSCs stably expressing GFP-degron-IRES-mCherry reporters were generated using the PiggyBac system. iPSCs were dissociated in Accutase and resuspended in mTeSR™Plus and analyzed on either BD Bioscience LSR Fortessa or LSR Fortessa X20 and FlowJo.

#### Western blotting

For whole cell lysates analysis, cells pellets were lysed in lysis buffer (40mM HEPES pH 7.5, 1mM MgCl2, 1% Triton X-100 and 150mM NaCl) supplemented with Roche complete protease inhibitor cocktail (Sigma, 11836145001) and benzonase (EMD Millipore, 70746-4) on ice. Samples were then normalized to total protein concentration using Pierce 660 nm Protein Assay reagent (Thermo Fisher, 22660). Next, 2x urea sample buffer (120 mM Tris pH 6.8, 4% SDS, 4 M urea, 20% glycerol and bromophenol blue) was added to the samples. SDS-PAGE and immunoblotting were performed using the indicated antibodies. Images were captured using ProteinSimple FluorChem M device.

#### Small-scale Immunoprecipitations

Transfections for co-immunoprecipitations were performed using polyethylenimine (PEI) (Polysciences, 23966-2) in a 1:6 ratio of μg DNA:μl PEI. ^6×Myc-^SKP1 was also co-transfected with FBXL14 in a 3:1 ratio of FBXL14:SKP1. Cells were transfected for 48 h, pelleted and resuspended in cold swelling buffer (20 mM HEPES-NaOH pH 7.5, 5 mM KCl, 1.5 mM MgCl2) with 0.1% Triton-X100, 2 mM NaF, 0.2 mM Na3VO4 and protease inhibitors (Roche, 11873580001). For small scale co-immunoprecipitations transfections were done in 10-cm plates and 500 μl of swelling buffer was used to resuspend cells. Cells were lysed for 30 min on ice, followed by 40 min centrifugation at 21,000g. Total protein concentration and volume of the lysate were normalized using Pierce 660 (Thermo, 22660). Normalized lysate was supplemented with NaCl to a final concentration of 150 mM and ANTI-FLAG M2 Affinity Agarose resin (Sigma-Aldrich, A2220) was added and incubated at 4 °C for 2 h. After four washes of the bound resin with cold wash buffer (20 mM HEPES-NaOH pH 7.5, 5 mM KCl, 1.5 mM MgCl2, 0.1% Triton-X100, 150 mM NaCl) bound proteins were eluted by addition of sample loading buffer and analyzed by western blotting. Images were captured using a ProteinSimple FluorChem M device.

#### Mass spectrometry

For mass spectrometry of endogenous TLE3^-3xFLAG^ 30 15-cm plates of HEK293T cells were used per condition. For FBXL14 mass spectrometry, HEK293T cells were transfected with 2.5 ug FBXL14^-3xFLAG^, 850 ng ^6×Myc-^SKP1 and 1.7ug dnCUL1 or 1.7 ug empty vector per 15 cm plate in total of 20 15-cm plates per condition. To prepare samples for mass spectrometry, bound proteins were eluted from anti-FLAG resin using 0.5 mg ml^-1^ 3×Flag peptide (Sigma, F4799), and proteins were precipitated overnight by the addition of trichloroacetic acid (Fisher, no. BP555) to a final concentration of 20% (w/v). Protein precipitates were washed three times in cold solution of 10 mM HCl in 90% acetone; resuspended in 8 M urea, 100 mM Tris-HCl, pH 8.5; reduced with 5 mM TCEP; and alkylated with 10 mM iodoacetamide. Samples were diluted with 100 mM Tris-HCl pH 8.5 to a 2 M urea concentration, supplemented with CaCl2 to 1 Mm concentration. Samples were trypsinized with 1 μl of 0.5 mg ml^-1^ trypsin (Promega) overnight at 37 °C, and formic acid was added to 5% final concentration. We used multidimensional protein identification technology (MudPIT) to analyze mass spectrometry samples. The analysis was performed by the Vincent J. Coates Proteomics/Mass Spectrometry Laboratory at UC Berkeley. To identify high-confidence interactors we performed CompPass statistical analysis ^104^ against mass spectrometry results from unrelated Flag immunoprecipitates performed in our laboratory.

For sequential IP-mass spectrometry of endogenous TLE3^-3XFLAG^/FBXL14^3XHA^ 30 15 cm plates of HEK293T cells per condition were transfected with 4 ug FBXL14^3XHA^ and 1.3 ug ^6×Myc-^SKP1 in 30 15-cm plates per condition and incubated for 48h. To prevent substrate degradation cells were treated with 500 nM MLN4924 overnight (Selleck Chemicals, NC1533706) prior to harvest. For endogenous ^3xHA-^TLE1 mass spectrometry 10 confluent 15 cm plates of iPSCs were used per condition. Samples for mass spectrometry were prepared similar as described above except for bound proteins were washed on resin three times with PBS, flash frozen and sent to further processing at UC San Diego Proteomics Facility. For whole-cell label-free quantitative DIA mass spectrometry 20 million iPSCs undergoing neural differentiation for 4 days were dissociated in Accutase, washed in PBS, pelleted, flash frozen in liquid nitrogen and sent to UC San Diego Proteomics Facility. Protein samples were diluted in TNE (50mM Tris pH8.0, 100mM NaCl, 1mM EDTA) buffer. RapiGest SF reagent (Waters) was added to the mix to a final concentration of 0.1%, and the samples were boiled for 5 min. TCEP was added to 1 mM (final concentration) and the samples were incubated at 37 °C for 30 min. Subsequently, the samples were carboxymethylated with 0.5 mg ml−1 of iodoacetamide for 30 min at 37 °C followed by neutralization with 2 Mm TCEP (final concentration). The proteins samples were then digested with trypsin (trypsin:protein ratio, 1:50) overnight at 37 °C. RapiGest was degraded and removed by treating the samples with 250 mM HCl at 37 °C for 1 h followed by centrifugation at 14,000 rpm for 30 min at 4 °C. The soluble fraction was then added to a new tube, and the peptides were extracted and desalted using C18 desalting columns (Thermo Fisher Scientific, PI-87782). Peptides were quantified using BCA assay and a total of 1 μg of peptides were injected for LC–MS analysis. Trypsin-digested peptides were analysed by ultra-high-pressure liquid chromatography (UPLC) coupled with tandem mass spectroscopy (LC–MS/MS) using nano-spray ionization. The nanospray ionization experiments were performed using a TimsTOF 2 pro hybrid mass spectrometer (Bruker) interfaced with nanoscale reversed-phase UPLC (EVOSEP ONE).

The Evosep method of 30 samples per day was performed using a 10 cm × 150 μm reversed-phase column packed with 1.5 μm C18-beads (PepSep, Bruker) at 58 °C. The analytical columns were connected with a fused silica ID emitter (10 μm inner diameter, Bruker Daltonics) inside a nanoelectrospray ion source (captive spray source, Bruker). The mobile phases comprised 0.1% formic acid as solution A and 0.1% formic acid/99.9% acetonitrile as solution B. The MS settings for the TimsTOF Pro 2 were as follows: the DIA-PASEF method for proteomics. The values for mobility-dependent collision energy were set to 10 eV. No merging of TIMS scans was performed. The ion mobility (IM) was set between 0.85 (1/k0) and 1.3 (1/k0) with a ramp time of 100 ms. Each method includes one IM window per DIA-PASEF scan with variable isolation window at 20 amu segments; 34 PASEF MS/MS scans were triggered per cycle (1.38 s) with a maximum of 7 precursors per mobilogram. Precursor ions in an m/z range of between 100 and 1,700 with charge states ≥3+ and ≤8+ were selected for fragmentation. Protein identification and label-free quantification were performed using Spectronaut 18.0 (Biognosys).

#### Immunofluorescence and confocal microscopy

iPSCs stably expressing either FBXL14^-3xFLAG^ or FBXL14^ΔCTE-3xFLAG^ were seeded on Matrigel coated 12mm glass coverslips at 500,000 cells/well in a 24-well plate with 500 ul mTeSR™Plus media (StemCell Technologies, 100-0276) supplemented with 50 nM Chroman I (MedChem Express, HY-15392). iPSCs subjected to neural conversion for 5 days using the STEMdiff™ Neural Differentiation medium with neural crest supplement (STEMCELL Technologies, 08610) were dissociated with Accutase and 400,000 cells were seeded on Matrigel coated chambered coverglass chambers (Lab-Tek, 155411) in 300 ul neural induction medium. The following day cells were fixed in a solution of 4% paraformaldehyde in 1X PBS for 20 min, followed by permeabilization with 0.5% Triton X-100 in 1X PBS for 15 min, and finally blocked with 10% fetal bovine serum (FBS) (VWR, 89510-186) in 1X PBS containing 0.05% Tween-20 for 30 min. Samples were probed with either anti-Flag M2 antibody (1:500) (Sigma-Aldrich, F1804), or AP2 alpha Antibody (1:200) (Novus Biologicals, NB100-74359) and anti-Pax-6 Antibody (1:500) (Biolegend, 901301) overnight at 4°C in 1X PBS containing 0.05% Tween-20 and 10% FBS. Following primary incubation, samples were incubated with either Goat anti-Mouse IgG (H+L) Cross-Adsorbed Secondary Antibody, Alexa Fluor™ 488 (Thermo Fisher Scientific, A11001) (1:500) or Goat anti-Rabbit IgG (H+L) Cross-Adsorbed Secondary Antibody, Alexa Fluor™ 488 (Thermo Fisher Scientific, A11008) (1:500) and Goat anti-Mouse IgG (H+L) Cross-Adsorbed Secondary Antibody, Alexa Fluor™ 568 (Thermo Fisher Scientific, A11004) (1:500) and staine with Hoechst 33342 (AnaSpec, 83218) (1:1000) for 1h. Coverslips were mounted onto microscope slides with ProLong gold antifade reagent (Thermo Fisher, P36934) and imaged using a Zeiss LSM 900 with Airyscan 2 microscope. Images were captured with a 20x air objective and Airyscan SR. Images were processed using Zen Blue (Zeiss) airyscan processing and Fiji.

#### Protein expression and purification

##### FBXL14^Strep^/SKP1 (WT and mutants)

N-terminally strep-tagged FBXL14/SKP1 was expressed in SF9 insect cells using the bac-to-bac baculovirus system according to manufacturer’s instructions (Thermo Fisher Scientific). Briefly, 1 µg of bacmid was transfected into SF9 cells using ExpiFectamine™ Sf Transfection Reagent (Thermo Fisher Scientific, A38915). After 5 days, virus was harvested (P1) and amplified twice further to generate high-titer P3 virus. 20 mL of P3 virus was used to infect 1L of SF9 cells, and cells were harvested ∼72 hours after infection. Purifications were conducted from 2-6L of cells. Cell pellets were resuspended in lysis buffer (50 mM Tris-HCl pH 8.0, 500 mM NaCl, 10% glycerol, 1 mM DTT) supplemented with EDTA-free protease inhibitor cocktail (Millipore Sigma, 04693132001) and benzonase (EMD-Millipore, #70746-4). All steps were carried out on ice or at 4 °C. Cells were lysed by sonication, followed by centrifugation at 17,000 xg for 1 hour at 4 °C. The clarified supernatant was incubated with StrepTactin 4Flow XT resin (IBA, 2-5010) for 1 hour in batch on a nutator in 50 mL conical tubes. Beads were collected by centrifugation, washed once in lysis buffer without protease inhibitors or benzonase, and loaded onto a gravity purification column (BioRad, 7374011). After washing the beads with ∼10-column volumes of lysis buffer, protein was eluted in buffer containing 50 mM biotin. The eluate was concentrated and further fractionated using a Superdex 200 increase (10/300) SEC column (Cytiva, 28990944) on an AKTA Pure 25 HPLC in storage buffer (25 mM HEPES-NaOH pH 7.5, 500 mM NaCl, 10% glycerol, 1 mM DTT). Protein was concentrated, aliquoted, flash-frozen, and stored at −80 °C for future use.

##### TLE1^HA-Strep^ (WD40 domain, miniTLE1; WT and mutants)

The HA-TLE1 WD40 domain and miniTLE1 constructs were expressed and purified as described above for FBXL14/SKP1 above from 1L of SF9 expression, except for the components of the lysis buffer (50 mM Tris-HCl pH 8.0, 150 mM NaCl, 1 mM DTT) and storage buffer (25 mM HEPES-NaOH pH 7.5, 150 mM NaCl, 1 mM DTT).

##### CUL1^6xHis^/RBX1

His-tagged CUL1 and RBX1 were expressed in SF9 insect cells using the bac-to-bac baculovirus system according to manufacturer’s instructions. Cells were lysed in buffer (50 mM Tris-HCl pH 8.0, 500 mM NaCl, 10 mM imidazle) supplemented with EDTA-free protease inhibitor cocktail (Millipore Sigma, 04693132001) and benzonase. Clarified supernatant was applied to a 1 mL HisTrap FF crude column (Cytiva, 11000458) and washed with lysis buffer containing 10 mM and then 30 mM imidazole. Protein complexes were eluted in buffer containing 300 mM imidazole, concentrated, and further fractionated using a Superdex 200 increase (10/300) SEC column (Cytiva, 28990944) in storage buffer (25 mM HEPES-NaOH pH 7.5, 500 mM NaCl, 1 mM DTT).

##### TLE^HA^ (Q-domain), UBE2R1, UbcH5c

N-terminally 6xHis-SUMO-TEV-tagged constructs were expressed in E. coli (BL21-CodonPlus(DE3)-RIL) cells grown in LB broth media. Cells were lysed in buffer (50 mM Tris-HCl pH 8.0, 500 mM NaCl, 10 mM imidazle) supplemented with EDTA-free protease inhibitor cocktail (Millipore Sigma, 04693132001) and benzonase. The clarified supernatant was applied to a 5 mL HisTrap FF crude column (Cytiva, 11000458), and washed with lysis buffer containing 10 mM and then 30 mM imidazole. Protein complexes were eluted in buffer containing 300 mM imidazole. TEV cleavage (produced in-house, UC Berkeley MacroLab; ∼1:50 w/w) was performed overnight at 4 °C in dialysis buffer (25 mM Tris-HCl pH 7.5, 200 mM NaCl, 2 mM β-mercaptoethanol). Cleaved product was filtered over a HisTrap column, and the flow-through concentrated and applied to a HiLoad™ Superdex 75 column (Cytiva, 28989333) in storage buffer (25 mM HEPES-NaOH, 150 mM NaCl, 1 mM DTT).

##### FBXL14^Strep^/SKP1-miniTLE1 complex for structure determination

Strep-TEV-FBXL14/SKP1 and 6xHis-MBP-TEV-^HA^TLE1^mini^ were co-expressed by co-infecting SF9 cells with two baculovirus (15 mL each/liter) as described above. Cell pellets were resuspended in lysis buffer (50 mM Tris-HCl pH 8.0, 150 mM NaCl, 1 mM DTT) supplemented with EDTA-free protease inhibitor cocktail (Millipore Sigma, 04693132001) and benzonase. Clarified supernatant was incubated with StrepTactin 4Flow XT resin (IBA, 2-5010) for 1 hour in batch on a nutator in 50 mL conical tubes. Beads were collected by centrifugation, washed once in lysis buffer without protease inhibitors or benzonase, and loaded onto a gravity purification column (BioRad, 7374011). After washing the beads with ∼10-column volumes of lysis buffer, protein was eluted in buffer containing 50 mM biotin. TEV cleavage (produced in-house, UC Berkeley MacroLab; ∼1:50 w/w) was performed overnight at 4 °C in dialysis buffer (25 mM Tris-HCl pH 7.5, 150 mM NaCl, 1 mM DTT). Cleaved product was concentrated and applied to a Superose 6 10/300 column (Cytiva, 29091596) in cryo-EM buffer (25 mM HEPES-NaOH, 150 mM NaCl, 1 mM DTT). Fractions containing the FBXL14/SKP1/TLE1^mini^ complex were pooled, concentrated to ∼10 mg/mL and immediately used for cryo-EM grid preparation.

#### Cryo-EM sample preparation, data collection, and processing

Cryo-EM samples were mixed with a final concentration of 0.02% (w/v) fluorinated octylmaltoside (Anatrace, O310F) immediately before cryo-freezing to prevent protein denaturation at the air–water interface. Then, 2.6 μL of the sample was applied to a glow-discharged 300-mesh Quantifoil R1.2/1.3 grid and incubated for 15 s before being blotted and plunge-vitrified in liquid ethane cool-protected by liquid nitrogen. Grid freezing was performed using a Mark IV Vitrobot (Thermo Fisher Scientific) system operating at 12 °C and 100% humidity.

Cryo-EM data were collected using a 300 kV Titan Krios G2 microscope (Thermo Fisher Scientific) equipped with a BIO Quantum energy filter (slit width 20 eV). Data were collected using SerialEM software at a nominal magnification of 105,000x with a pixel size of 1.05 Å per pixel. Movies were recorded using a 6k x 4k Gatan K3 Direct Electron Detector operating in super-resolution CDS mode. Each movie was composed of 40 subframes with a total dose of 50 e-/A2, resulting in a dose rate of ∼1.25 e-/A2. Data processing, including motion correction, CTF estimation, particle picking, 2D class averaging, and 3D refinement, was performed using cryoSPARC v.4.3 workflow, using mostly default settings. All movies were 2x binned and patch motion corrected. After particle picking and several iterations of 2D class averaging, the initial 3D volume was calculated using several rounds of ab initio 3D reconstruction. For the cryo-EM map containing the full FBXL14/SKP1/miniTLE1 tetrameric structure, 3D classes containing this structure were curated at the ab initio 3D reconstruction stage. For the higher-resolution focused refinement of the dimeric FBXL14/SKP1/TLE1-WD40 region, a soft mask was generated of this region using ChimeraX. A larger subset of particles than were used for the tetrameric structure were utilized for focused refinement. A full workflow of cryo-EM data processing is outlined in Supplemental Figure 2. Selected particles were then used for non-uniform 3D refinement. Default B-factor sharpening in CryoSparc was used to generate the final deposited maps.

#### Model building and structural analysis

The initial models for the full FBXL14/SKP1/TLE1^mini^ complex were generated using AlphaFold3 (FBXL14/SKP1 and TLE1 Q-tetramer) using UniProt sequences Q8N1E6, P63208, and Q04724 or an existing crystal structure for the TLE1 WD40 domains (PDB: 2CE9). First, a model of the dimeric FBXL14/SKP1/TLE1^WD40^ complex was built into the high resolution cryo-EM map (EMD-75775) by manually fitting the protomers into the density in ChimeraX (version 1.9) and deleting regions without corresponding density. Initial refinement was performed using ISOLDE (version 1.10.0) within ChimeraX, followed by iterative rounds of model building/adjustment and refinement in Coot (version 0.9.8.96) and Phenix (version 2.0). Next, FBXL14/SKP1/TLE1^WD40^ (2 copies) were fit into the cryo-EM density map for the full FBXL14/SKP1/TLE1^mini^ complex (EMD-75774), the TLE1 Q-domain tetramer was added, and chains merged with the WD40 domains. A final round of refinement was performed in Phenix on the full complex, constraining side-chains to best represent the high-resolution data. All structural figures were assembled using ChimeraX. For the composite model of FBXL14/SKP1/TLE1^mini^/CUL1/RBX1/E2-Ub shown in Figure 4A, a crystal structure containing CUL1 and SKP1 (PDB: 1LDK) was aligned to the complex via SKP1. From there, an AlphaFold3 model of a RBX1/UbcH5c/Ub complex that resembles experimental structures of RING/E2-Ub complexes was aligned to the RBX1 protomer in PDB: 1LDK.

#### In vitro transcription/translation (IVT/T)

Full-length TLE proteins or substrates in pCS2+ were synthesized using the TnT quick coupled rabbit reticulocyte lysate in vitro transcription and translation (IVTT) system (Promega, no. L2080). Each 12.5-μl in vitro reaction contained 10 μl of rabbit reticulocyte lysate, 0.3 μl of ^35^S-Methionine (11 μCi μl−1, Promega, no. NEG009H005MC), and 600 ng of pCS2+ construct. Reactions were mixed and incubated at 30°C for 1 h. Approximately 10% of the IVTT was saved as input.

#### In vitro binding assays

IVTT reactions containing synthesized TLEs or substrates were mixed with 500 μl PBST (PBS (Gibco, 14190144) + 0.1% Triton X-100), 2 uM carfilzomib, 15 μl amylose resin (NEB, E8021L), 8 μg of ^His-MBP-^FBXL14/SKP1 complex or ^His-^MBP and with or without recombinant ^HA-^miniTLE1 and incubated at 4°C for 3 h. Resin was washed five times with cold PBST supplemented with 300 mM NaCl and eluted with the sample loading buffer. Pulldowns were analyzed by SDS-PAGE and autoradiography.

For in vitro HA-binding assays 8 μg of WT or interface mutant recombinant FBXL14/SKP1 proteins were mixed with 500 μl PBST, 10 μl EZviewer Red Anti-HA Affinity resin (Millipore, E6779) with or without WT or interface mutant recombinant ^HA-^ TLE1 proteins and incubated for 3 h at 4 °C. Resin was washed five times with cold PBST supplemented with 300 mM NaCl and eluted with the sample loading buffer. Pulldowns were analyzed by SDS-PAGE and western blot.

#### In vitro ubiquitylation assays

To activate SCF complexes recombinant Cul1 was first subjected to neddylation reaction. For each neddylation reaction 8 μM recombinant CUL1/RBX1 was mixed with 0.5 μM UBA3/APPBP1, 1 μM UBE2M, 25 mΜ Nedd8, 1x energy mix (10x: 150 mM creatine phosphate, 20 mM ATP, 20 mM MgCl2, 2 mM EGTA, pH to 7.5 with KOH) in 1x neddylation buffer (10X: 500 mM Tris Ph 7.5, 1000 mM NaCl and 25 mM MgCl2). Reaction was incubated at 25°C for 25 min and was quenched with 1 mM DTT.

In vitro ubiquitylation assays were performed in a 10 μl reaction volume: 0.5 μl of 10 mM E1 (250 nM final), 0.5 μl of 25 μM UBE2R1 (1.25 μM final), 0.5 μl of 25 μM UbcH5c (1.25 μM final), 0.5 μl of 1 mg ml^-1^ TUBE1 (LifeSensors, UM101), 1 μl of 10mg ml^-1^ ubiquitin (1 mg ml^-1^ final) (R&D Systems, U-100H), 1 μl of 100 mM DTT, 1 μl of 10x energy mix, 1 μl of 10x ubiquitylation assay buffer (250 mM Tris 7.5, 500 mM NaCl and 100 mM MgCl2), 1.5 μl of neddylated CUL1 (1.2 μM final), 0.5 μl FBXL14/SKP1 (1.2 μM final), 1 μl miniTLE1 (1.2 μM final) and 1 μl of 20 μM TAMRA-labeled substrate peptide. Reactions were incubated at 30°C for 30 min and eluted in 2X urea sample buffer and resolved on SDS-acrylamide 4-20% gradient gels and imaged using a ProteinSimple Fluorchem M imager. To test for polyubiquitin chain specificity recombinant ubiquitin mutants were used (R&D Systems, UM-NOK, UM-K48R, UM-K63R, UM-K4863R).

#### CRISPR-Cas9 genome editing

All CRISPR-Cas9 edited HEK-293T cell lines are generated using an overexpression pX330 (Addgene, 42230) vector containing a specific guide and encoding for spCas9. sgRNA sequences were designed using the online tool CRISPOR (https://crispor.gi.ucsc.edu/crispor.py). sgRNA DNA oligonucleotides were phosphorylated (NEB, M0201), annealed and ligated (NEB, M0202) into pX330 plasmid. 5x10^5^ cells were cultured in a 6-well plate and next day 2 μg of px330 plasmid and 0.6 μl of 100 μM single stranded donor oligo for knock-ins using 6 μl Mirus TransIT-293 Transfection reagent (Mirus, MIR2704). Two days post transfection cells were subjected to clonal selection in 96-well plates. Homozygous clones were confirmed by PCR genotyping, DNA sequencing, western blot and qPCR analysis when applicable. DNA guide oligonucleotide and donor sequences are listed in the Key resources table section.

All CRISPR-Cas9 edited stem cell lines were generated using Cas9-RNP complexes, the Lonza 4D-Nucleofector X Unit (program CA137) and the P3 Primary Cell 4D-Nucleofector™ X Kit S (Lonza, V4XP-3032). For each reaction, 2 µl of recombinant Cas9 (20 µM) was mixed with 3 µl of sgRNA (100 µM) and incubated for 10 min at room temperature, after which 0.7 µl of ssODN (100 µM) was added. Stem cells were disassociated in Accutase, washed in PBS and 5x10^5^ cells were resuspended in 20 µl Lonza P3 Nucleofector™ solution, mixed with the RNP complex. Immediately after nucleofection cells were transferred to one Matrigel coated 6-well with 2 ml mTeSR™Plus and 50 nM Chroman I. 72 H post nucleofection bulk editing efficiency was determined by PCR and cells were subjected to single clonal selection in 6-well plate format with

2 ml mTeSR™Plus, 50 nM Chroman I, 5 μM Emericasan (Sigma-Aldrich, SML2227), 1x Polyamine Supplement (1000×) (Sigma-Aldrich, P8483) and 700 nM trans-ISRIB (R&D Systems, 5284). After 5-7 days each individual colony originated from a single clone was manually picked and transferred to a single Matrigel-coated well of 24-well plates with 500 µl mTeSR™Plus and without Chroman I. Homozygous clones were confirmed by PCR genotyping, DNA sequencing, western blot and qPCR analysis when applicable. DNA guide oligonucleotide and donor sequences are listed in the Key resources table section.

#### MNase ChIP-qPCR

iPSCs were crosslinked in one 15 cm plate per IP with PBS containing 1% paraformaldehyde (ThermoFisher, 28908) for 10 min at room temperature and quenched with 125 mM glycine for 5 min. Cells were washed twice in cold 1x PBS and harvested in 1x PBS plus 0.25 mM PMSF, 10 μg/mL of aprotinin (ThermoFisher, 78432) and 1 mM benzamidine (ThermoFisher, 401790250). Cell pellets were frozen in liquid nitrogen and stored at −80°C for later use. Frozen pellets were resuspended in 1 ml of lysis buffer (5 mM PIPES pH 8.0, 85 mM KCL 0.5% NP4, 1 × cOmplete protease inhibitor cocktail and 1× phosSTOP) for 10 min on ice and spun at 4000rpm for 10 mins at 4°C to pellet the nuclei. Nuclei were resuspended in dilution buffer (1% Triton X-100, 150 Mm NaCl, 20 mM Tris 8.0, 2.5 mM CaCl2, 1× cOmplete protease inhibitor cocktail and 1× phosSTOP), digested with 45 units of MNase (Worthington, LS004798) per 100 μl of pellet volume for 5 min at 37 °C, quenched with 6 mM EDTA and 6 mM EGTA, spun at 20,000*g* and frozen in liquid nitrogen and stored at −80 °C for later use. 10-20 ug DNA was used per IP. For FBXL14^-3xFLAG^ ChIP-qPCR MNase-digested chromatin was incubated overnight at 4 °C with 5 ug of anti-FLAG clone M2 antibody (Sigma-Aldrich, F1804). Flag antibodies were immunoprecipitated by addition of BSA-blocked ANTI-FLAG M2 Affinity resin (Sigma-Aldrich, A2220) and control samples were incubated with BSA-blocked Mouse IgG-Agarose resin (Sigma-Aldrich, A0919) for 2 hr at 4°C. For endogenous ^3XHA-^TLE1 ChIP-qPCR MNase-digested chromatin was incubated overnight at 4 °C with 5 ug anti-HA tag antibody (Abcam, ab9110) or unspecific IgG antibody (Abcam, ab171870). Next day samples were incubated with BSA-blocked Protein G Dynabeads™ (ThermoFisher, 10004D) for 2 hr at 4°C. Beads were washed twice with low salt wash buffer (20 mM Tris 8.0, 150 mM NaCl, 2 mM EDTA, 1% Triton X-100 and 0.1% SDS), twice with high salt wash buffer (20 mM Tris 8.0, 500 mM NaCl, 2 mM EDTA, 1% Triton X-100 and 0.1% SDS), once with LiCl buffer (20 mM Tris 8.0, 250 mM LiCl, 1 mM EDTA, 1% deoxycholate and 1% Nonidet P-40), and twice with 1× TE. Samples were eluted twice at 30 °C with 1% SDS buffered in 1× TE. Eluates were pooled, treated with RNase A and reverse crosslinked overnight at 65 °C. Samples were then treated with proteinase K (ThermoFisher, EO0491), phenol:chloroform (ThermoFisher, 15593031) extracted, isopropanol precipitated and eluted in 10 mM Tris 8.

Following primers were used in this paper:

ASCL1_F_ GTAGGAGAGGAACGCGAGACGC

ASCL1_R_ CCGCTGGCGCCTTCTTGTTTCT

FOXB1_F_TCTCGCTCAGTCCCCTCCCTCT

FOXB1_R_ GAAAAGACCCCTGTAGCGGCGC

PAX6_F_ GCACTTAGTCAACAAATGGCACGTGGG

PAX6_R_ GCGGAGCAGAGGCACAGCTC

OTX1_F_ CTCGCGTTCACATACCCGGGGA

OTX1_R_ CGGAGTGGCTTGTCTTGCTCGG

OTX2_F_ AGAGGAGCATGGCGGCCTGTAA

OTX2_R_ ACCAGTTGCTGTGTCCTCGCTC

SOX2_F_ TGACAGCCCCCGTCACATGGAT

SOX2_R_ AGGCAGCAAACTACTTTCCCCCT

#### Cleavage Under Targets and Release Using Nuclease (CUT&RUN)

Endogenous ^3xHA^TLE1 CUT&RUN experiments from either WT or FBXL14^ΔCTE^ iPSCs undergoing neural conversion for 4 days were performed in biological triplicates using CUTANA™ ChIC/CUT&RUN Kit (EpiCypher, 14-1048) according to the manufacturer’s protocol (User Manual Version 6.0). Cells at 4 days of neural conversion were dissociated in Accutase, washed in PBS and 6x10^5^ cells was used per condition. Cells were permeabilized using a final working concentration of 0.015% digitonin. Endogenous ^3xHA^TLE1 was bound using 1 μg anti-HA tag antibody (Abcam, ab9110) and 1 µl of manufacturer supplied IgG antibodies were used as negative control and incubated at 4 °C overnight. Samples were washed with cell permeabilization buffer and pAG-MNase was added to each sample and incubated at room temperature for 15 min. Targeted chromatin digestion was initiated by adding calcium chloride to each sample and samples were digested at 4 °C for 1 h. Reactions were quenched with stop buffer followed by incubation at 37 °C for 10 min to release digested chromatin fragments. DNA was purified using SPRIselect beads, washed with 85% ethanol and eluted in 0.1X TE Buffer. DNA concentration was determined using the Qubit dsDNA High Sensitivity assay kit (Thermo Fisher, Q33230). Libraries were prepared using the CUTANA™ CUT&RUN Library Prep Kit (EpiCypher, 14-1001). Library DNA size distribution was assessed using a Fragment Analyzer (Agilent). Final libraries were pooled and sequenced by QB3 Genomics, UC Berkeley, Berkeley, CA (RRID:SCR_022170) on an NextSeq P1 150PE (Illumina). Pair-end reads were quality-checked with FastQC (version 0.12.1), trimmed with cutadapt (version 2.6) and aligned onto the human genome (hg38 assembly) using Bowtie2 (version 2.5.4). Peaks were called with MACS3 (version 3.0.3) and annotated using Homer on WSL (Windows Subsystem for Linux) platform. To generate read density clustered heat maps downstream bioinformatic analyses were performed using Python (version 3.13.10) in Jupiter Notebook-Lab and deepTools in Galaxy (web-based platform for data analysis). Peak scores for target genes were averaged and concatenated between three biological replicas and different conditions aligned against each other based on their peak scores. Ultimately, three clusters were identified: 1^st^ cluster showing genomic sites with similar enrichment in both conditions (−1< z-score <1), 2^nd^ cluster with higher enrichment in ΔCTE-FBXL14-iPSCs

(z-score ≥1) and 3^rd^ cluster with higher enrichment in WT-FBXL14-iPSCs (z-score ≤ −1). Heat maps and plot profiles of the corresponding clusters were generated by aligning coverage reads (bigwig files) from WT (first heat map panel in Figure 5E) and ΔCTE (second heat map panel in Figure 5E) to genomic sites identified as peaks with MACS3. For more transparency, the same clusters were represented as log2 ratio of the coverage reads from ΔCTE-FBXL14-iPSCs and WT-FBXL14-iPSCs in identified enrichment sites (third heat map panel in Figure 5E). GO analysis was performed using Homer on WSL.

To quantify change in target region binding CUT&RUN-qPCR was performed. CUT&RUN libraries were diluted 20-fold and samples from three biological replicas were run in technical triplicates and normalized to their own IgG controls. Fold changes in binding region enrichment were calculated using the ΔΔCt method.

Following primers were used in this paper:

FWD_cr_NEUROD1_ TTGGCCCTGTGAATGCTTCGCC

REV_cr_NEUROD1_ AGGGGAAGGGGGAGGCATGT

FWD_cr_ATOH1_TTTTGGTTGAGCTGGTGTCCCA

REV_cr_ATOH1_TGGCGATGGTGGTCTCCCAACT

FWD_cr_NEUROG2_TCACCCGGGAGTTAGGAGGG

REV_cr_NEUROG2_AGCTCATTAGGGGCCCGAGC

FWD_cr_FIZ1_AGGAAAGCCGACGGACTTCCCC

REV_cr_FIZ1_GCCCCTAAAAGGTGCAGCCCTG

FWD_cr_LMX1A_TGCCAGTCTAGTCCAGGGCGAA

REV_cr_LMX1A_TGTTTCTGCGCCTGGCCTCTTG

FWD_cr_RTN4_ACCCCACCCCAACTCACGGTTA

REV_cr_RTN4_CCGGGCCAACAGAAAGGGAAGG

#### qPCR

Total RNA was purified using a nucleospin RNA kit (Macherey-Nagel, 740955). cDNA was generated using a RevertAid First Strand cDNA Synthesis kit (Thermo Fisher Scientific, K1622) and RT-qPCRs were performed using 2×KAPA SYBR Fast qPCR master mix (Roche, KK4602) on a LightCycler 480 II Instrument (Roche) using. Fold changes in expression were calculated using the ΔΔCt method.

#### RNAseq sample preparation and analysis

Wild-type, FBXL14^ΔCTE^, TLE1^ΔWD40^ or TLE1^R610S^ iPSCs were subjected for neural conversion for 5 days and collected for transcriptomic analysis. Total RNA was extracted from cells using the NucleoSpinRNA kit (Macherey-Nagel, no. 740955) according to the manufacturer’s instructions. Library preparation, sequencing and bioinformatic statistical analysis was performed by Novogene Bioinformatics Technology Co., Ltd. (Beijing, China). qPCR validations were done from each sample cDNA in biological and technical triplicates. For each condition fold changes in gene expression were first derived from their day 0 and day 5 time points of neural conversion and then fold differences in gene expression for FBXL14^ΔCTE^, TLE1^ΔWD40^ or TLE1^R610S^ conditions were compared to WT cells.

Following primers were used in this paper

PAX6_F_CACATGAACAGTCAGCCAATG

PAX6_R_GGCCAGTATTGAGACATATCAGG

ASCL1_F_AGCTTCTCGACTTCACCAAC

ASCL1_R_ CAACGCCACTGACAAGAAAG

NEUROD4_F_AGACGCAGATTCACAGAGTTC

NEUROD4_R_GAGCCCAGACCTTTATCCATC

FOXB1_F_GCGCAACTTGAAGCAACT

FOXB1_R_CTTCTGGTCGCTGTACGTG

OTX1_F_GATCCAGGTAGATGGTGAACG

OTX1_R_ACCACGCAGTCCTCCAG

OTX2_F_GCTGAGTCTGACCACTTCG

OTX2_R_CATTCTGCTGTTGTTGCTGTT

LMX1A_F_AGAACTTCCAAAGCGCGAT

LMX1A_R_AGAAACCTGTCCAAGATGACC

POU4F1_F_CTCACTTTGCCATGCATCC

POU4F1_R_AGCAGCGTCTCGTCCAG

FBXL14_F_TGCGCTCCTGTGACAACATC

FBXL14_R_ TGGGCTATGTAAGCCAGACTC

SOX10_F_ CTTTCTTGTGCTGCATACGG

SOX10_R_ AGCTCAGCAAGACGCTGG

SNAI2_F_ TGACCTGTCTGCAAATGCTC

SNAI2_R_ CAGACCCTGGTTGCTTCAA

FOXD3_F_ TTGACGAAGCAGTCGTTGAG

FOXD3_R_ TCTGCGAGTTCATCAGCAAC

EDNRA_F_ CATGACTTGTGAGATGTTGAACAG

EDNRA_R_ CTGTTTTTGCCACTTCTCGAC

NR2F1_F_ CTCAAGAAGTGCCTCAAAGTG

NR2F1_R_ AGATGTAGCCGGACAGGTAG

TFAP2A_F_ATTGACCTACAGTGCCCAGC

TFAP2A_R_ ATGCTTTGGAAATTGACGGA

HOXB1_F_TAAGAAAACCCACCCAAGAC

HOXB1_R_ACTCCTTTTCCAGTTCTGTCAG

HOXB2_F_TCGGATCGCCTGCAGAT

HOXB2_R_CGGCACAGGTACTTATTAAAGTG

GAPDH_F_ACATCGCTCAGACACCATG

GAPDH_R_TGTAGTTGAGGTCAATGAAGGG

#### Quantification and statistical analysis

All quantifications are presented as the mean ± standard deviation. Significance was determined by two-tailed t test, ns p > 0.05 and significant p<0.05. Flow cytometry, RNA-seq, qPCR and CHIP experiments were done in biological triplicates and shown as the mean ± standard deviation.

